# Lateralized cerebellar connectivity differentiates auditory pathways in echolocating and non-echolocating whales

**DOI:** 10.1101/2024.09.18.609772

**Authors:** Sophie Flem, Gregory Berns, Ben Inglis, Dillon Niederhut, Eric Montie, Terrence Deacon, Karla L. Miller, Peter Tyack, Peter F. Cook

## Abstract

We report the first application of diffusion tractography to a mysticete, which was analyzed alongside three odontocete brains, allowing the first direct comparison of strength and laterality of auditory pathways in echolocating and non-echolocating whales. Brains were imaged post-mortem at high resolution with a specialized steady state free precession diffusion sequence optimized for dead tissue. We conducted probabilistic tractography to compare the qualitative features, tract strength, and lateralization of potential ascending and descending auditory paths in the mysticete versus odontocetes. Tracts were seeded in the inferior colliculi (IC), a nexus for ascending auditory information, and the cerebellum, a center for sensorimotor integration. Direct IC to temporal lobe pathways were found in all animals, replicating previous cetacean tractography and suggesting conservation of the primary auditory projection path in the cetacean clade. Additionally, odontocete IC-cerebellum pathways exhibited higher overall tract strength than in the mysticete, suggesting a role as descending acousticomotor tracts supporting the rapid sensorimotor integration demands of echolocation. Further, in the mysticete, contralateral right IC to left cerebellum pathways were 17x stronger than those between left IC and right cerebellum, while in odontocetes, the laterality was reversed, and left IC to right cerebellum pathways were 2-4x stronger than those between right IC and left cerebellum. The stronger left IC-right cerebellum connectivity observed in odontocetes corroborates the theory that odontocetes preferentially echolocate with their right phonic lips, as the right phonic lips are likely innervated by left-cortical motor efferents that integrate with left-cortical auditory afferents in right cerebellum. This interpretation is further supported by the reversed lateralization of IC-cerebellar tracts observed in the non-echolocating mysticete. We also found differences in the specific subregions of cerebellum targeted by the IC, both between the mysticete and odontocetes, and between left and right sides. This study establishes foundational knowledge on mysticete auditory connectivity and extends knowledge on the neural basis of echolocation in odontocetes.

## Introduction

### Cetacean brains and behavior

Cetaceans present a fascinating set of species for studying changes in brain and behavior following their evolutionary adaptation to an aquatic environment some 60 million years ago (Gatesy et al., 2013). Observations of both wild and captive cetaceans suggest they possess a wide range of complex sensory and cognitive capabilities (Marino et al., 2007). Many species engage in extensive communication with conspecifics and other cetaceans, facilitating their rich and intricate social lives, and possibly enabling the development of learned individual identity signals (Janik et al., 2006) and rudimentary culture (Fox et al., 2017; Pettit & McCulloch, 2023). The sensory, behavioral, social, and cognitive complexity observed in cetaceans has motivated scientific study of their similarly complex, large, and highly convoluted brains (Ridgway et al., 2017). Though there are 94 known living cetacean species (Committee on Taxonomy, 2023), most studies on neuroanatomy and behavior have involved but a small number of these species, particularly bottlenose dolphins (*Tursiops truncatus*), orcas (*Orcinus orca*), and beluga whales (*Delphinapterus leucas*) (Pettit & McCulloch, 2023). Notably, there is a particular dearth of work on baleen whales (parvorder Mysticeti). Given the evolutionary split between odontocetes and mysticetes 38.8 million years ago (Marx & Fordyce, 2015), and their divergent sensorimotor capabilities, neuroanatomical comparisons between these closely-related suborders may be particularly fruitful.

Though recent work has included important advances such as multimodal methods that integrate analyses of the gross neuroanatomical and cellular levels (Orekhova et al., 2022; Raghanti et al., 2018), the practical and ethical difficulty of carrying out non-invasive functional neuroimaging in large, fully-aquatic mammals like cetaceans has prevented researchers from clearly and conclusively elucidating the *specific functions* of the myriad regions in the great, convoluted mass of the cetacean cortex. Current understanding relies predominantly on decades-old invasive functional studies (Ladygina & Supin, 1977) patched together with more recent cytoarchitectural, gross anatomical, and behavioral characterization (Cook et al., 2024; De Vreese et al., 2024; Hof et al., 2005; Marino et al., 2002; Morgane et al., 1980). In other words, though the behavioral and cortical complexity of cetaceans is well-established (Marino et al., 2007), the question of *how* the complex brain functions to subserve complex behavior remains largely undetermined.

### Cetacean auditory system

The cetacean auditory system serves as an ideal starting point for understanding the mechanisms by which cetacean brains subserve their behavior. Because of the long range over which sound travels through seawater and the reduced range of light in much of the ocean, the cetacean nervous system evolved to specialize in auditory processing (Oelschläger et al., 2007). There is clear evidence of rapid evolutionary adaptation of external auditory structures in the cetacean lineage (Nummela et al., 2004), and the need to process complex auditory information may have contributed heavily to the dramatic encephalization seen in cetaceans (Ridgway, 1990). Further, different suborders of cetacea have evolved to do very different things with sound; while many utilize passive audition in service of complex communication, navigation, foraging, and hunting (Kremers et al., 2016), toothed whales (suborder Odontoceti) additionally evolved the capability for a more active form of audition: echolocation (Au, 1993). Echolocation, also known as sonar, occurs when an organism emits sound waves in order to gain information about features of its environment from the echoic return signals (Moss et al., 2023). Paleontological evidence suggests that echolocation emerged in the earliest diverging odontocetes during the Oligocene, approximately 30 million years ago (Park et al., 2016), and evolved rapidly following its initial emergence due to its affordance of plentiful access to prey across otherwise acoustically complex environments (Coombs et al., 2022; Marx et al., 2016). Across mammal species, subcortical auditory integration is time-pressured and requires very rapid processing (Frisina, 2001), time demands that are likely compounded by the high frequencies of echolocation signals in dolphins. New capacities to hunt in muddy rivers, shallow icy waters, and high-pressure ocean depths likely provided the caloric payoffs requisite to carry out this energetically-demanding, time-pressured processing (Coombs et al, 2020; Laeta et al., 2023). The continuous refinement of echolocation capacities in odontocetes has persisted to the present day, as can be seen in neuroanatomical studies reporting the immense hypertrophy of the modern-day odontocete subcortical auditory system. The absolute size of the dolphin medial geniculate nucleus (MGN) is seven times larger, the inferior colliculi (IC) twelve times larger, and the lateral lemniscus *250* times larger than in the comparably-sized human brain (Bullock & Gurevich, 1979; Cozzi et al., 2018). The bulk of prior research on auditory adaptations in cetaceans has focused on the diencephalon, mesencephalon, or myelencephalon, where brain structures and their connectivity patterns are more conserved across clades. However, as discussed above, difficulties in collecting functional data on the cetacean neocortex has rendered the task of identifying the cortical components of the auditory system quite difficult.

Foundational studies establishing the locations of sensory projection fields in cetacean neocortex employed invasive electrophysiological methods that directly measured which regions of cortex responded to different types of sensory input. These studies initially localized the primary auditory projection field (A1) in a long “belt” of dorsal cortex in the suprasylvian gyrus, near the vertex of the brain, and directly lateral to the primary visual cortex (Ladygina et al., 1978; Sokolov et al., 1972; Supin et al., 1978). Despite one prior study with opportunistic evidence of electrophysiological response to sound in the temporal lobe (Bullock & Ridgway, 1972) and subsequent tracing identifying medial geniculate projections to the temporal operculum (Revishchin & Garey, 1990), later studies tended to accept the localization of a suprasylvian A1 (Morgane et al., 1988; Hof et al., 2005; Cozzi et al., 2018). This convention was challenged in Berns et al. (2015), a first-of-its-kind study that assessed auditory connectivity in opportunistically-collected post-mortem dolphin brains. Berns et al. used a diffusion-weighted steady state free precession (DW-SSFP) protocol specialized for post-mortem imaging (Miller et al., 2012) and conducted diffusion tensor imaging (DTI) tractography on the resulting diffusion-weighted images to allow analysis and visualization of cerebral white matter tracts (Basser et al., 2000). Critically, diffusion tractography allows some inference of functional connectivity from structural connectivity, thus granting researchers a window into cetacean brain function, albeit an indirect one (Hermundstad et al., 2013; Reddi, 2017; Wang et al., 2020). Berns et al. reported that in the dolphins studied, the ascending auditory pathway projected not to the suprasylvian gyrus, but instead to a region in the deep temporal lobe, i.e. the more typical location of mammalian A1. This suggested that there may be multiple topographically distinct cortical centers for auditory processing in the dolphin. Indeed, the dorsal electrode insertions used in the original electrophysiology studies on cortical auditory processing may have simply missed this site due to its harder-to-reach location in deep temporal lobe. In this case, it is possible that earlier invasive studies were in fact measuring suprasylvian auditory processing signals that were “downstream” of temporal A1. Odontocete auditory processing may thus parallel the case in bats, in which temporal A1 sends auditory information to postero-dorsal centers that process finer temporal and spatial features of sound (Kössl et al., 2014). In the mysticetes, meanwhile, neither electrophysiological nor tractographical methods have yet been applied to study ascending auditory pathways.

### Lateralization

Following Berns et al. (2015), Wright et al. (2018) used diffusion-weighted MRI, though with a standard echoplanar protocol, to examine the lateralization observed in the major white matter tracts in the brain of a *Tursiops truncatus* specimen. Wright et al. found evidence for substantial cortical asymmetry, particularly in putative auditory and vocal tracts. This finding is in line with prior evidence of sparse interhemispheric connectivity in cetaceans and, relatedly, unihemispheric sleep (Lyamin et al., 2002, 2008; Tarpley & Ridgway, 1994). In addition, in at least some species of echolocating odontocetes, including beluga whales and dolphins, individuals have two sets of lateralized phonic lips that are employed differentially depending on function: while the left lips are responsible for social communication, the right lips are responsible for high-frequency echolocative clicks (Ames et al., 2020; Madsen et al., 2013). Importantly, Ames et al. (2020) report that *function*, not *frequency*, determines which phonic lips are employed to produce sound, as even high-frequency social buzzes are typically performed by the left lips, in addition to low-frequency social whistles.

The lateralization of phonic lip function, also known as the functional laterality hypothesis, provides a new foundation upon which to bridge cetacean behavior with brain function. Despite limited direct data on functional lateralization in cetacean cortex, the principle of contralateral organization, in which each half of the forebrain and thalamus has efferent and afferent connections to the opposite side of the body (Whitehead & Banihani, 2013), applies broadly to vertebrate species all the way back to the Ordovician period (Janvier, 2002). It is safe to presume that cetacean nervous systems are organized contralaterally as well. If the left phonic lips are employed for social functions and the right phonic lips for echolocation, then the contralateral cortical hemispheres may be accordingly specialized to process social and echolocative information, too– and such functional neural lateralization is particularly plausible given the previous evidence of hemispheric independence in cetaceans. Such a hypothesis may be investigated by studying efferent motor pathways from the central nervous system to the phonic lips, and by studying the pathways prior to these that prepare the appropriate motor information via the integration of auditory, premotor, and social or navigational information. In other words, investigation of this hypothesis may require analysis of not only ascending auditory pathways, but also descending acoustimotor pathways, the latter of which have been relatively unexplored in cetaceans thus far.

### Descending acousticomotor pathways

Although the classical representation of the auditory system follows an ascending pathway from brainstem nuclei to inferior colliculi and then medial geniculate nucleus before reaching A1, recent data from human and animal models shows strong evidence of dense reciprocal connections, ascending and descending between every major waystation on the auditory pathway (Huffman & Henson, 1990; Malmierca, 2004). The inferior colliculus is central in this reciprocal architecture, serving as the termination point for fibers ascending from the brainstem auditory nuclei while also receiving heavy top-down innervation from primary auditory cortex (Malmierca, 2004). The top-down connectivity allows for feedback from auditory cortex onto the IC and to the brainstem auditory nuclei, with potentially faster transit time from cortex to brainstem than vice versa (Huffman & Henson, 1990). This suggests that top-down cortex to IC pathways could play an essential role in modulating early auditory processing. In addition, descending acoustic information from the cortex and IC has been shown to densely target the cerebellum, likely to facilitate motor response to auditory information (Powell & Hatton, 1969; Saldaña & Merchan, 1992). Along with sensorimotor processing, the cerebellum has also been found to play a role in general audition (Baumann et al., 2014) and speech processing in humans (McLachlan & Wilson, 2017).

The cerebellum is of particular interest in cetacean neurobiology due to its clear hypertrophy, particularly among delphinids (Ridgway et al., 2017). In common, bottlenose, and Atlantic white-sided dolphins, the cerebellum comprises an average of approximately 15% of total brain volume, as compared to the 10% and 9% of total brain volume observed in humans and non-human primates, respectively (Marino et al., 2000; Montie et al., 2008; Ridgway, 1990). While generally understudied, one explanation for cerebellar hypertrophy in dolphins is that it is involved in some way in vocal and echolocation processing (Oelschlager, 2008). This hypothesis would parallel findings in bats, which also have relatively large cerebella compared to evolutionarily related non-echolocating rodents (Alvarez et al., 2023). Electrophysiological recording in bats has identified cells in bat cerebellum that play a specific role in echolocation, helping predict target location (Jen & Schlegel, 1980). The role of descending auditory tracts in dolphins requires further exploration, and particularly with respect to the expanded cerebellum, given its potential role in auditory processing and echolocation. Furthermore, dolphins can apparently remain continuously vigilant for *weeks* at a time by using echolocation during unihemispheric sleep (Branstetter et al., 2012), suggesting that sufficient echolocation processing is possible in either hemisphere (despite functional laterality at the level of the phonic lips), *or* that subcortical and cerebellar mechanisms may support the capability for echolocation during unihemispheric sleep (Ridgway et al., 2015).

In sum, while DTI tractography has been used to investigate ascending auditory pathways (Berns et al., 2015; Orekhova et al., 2022), prefrontal cortex (Gerussi et al., 2023), and lateralization (Wright et al., 2018) in *toothed whales*, to date, it has not been employed in the analysis of descending acoustimotor pathways that may be critical for echolocation, and it has never been employed to study baleen whale brains. Baleen whales have a partially conserved laryngeal sound production mechanism (Elemans et al., 2024), which is quite different from the nasal sound production mechanisms of the odontocetes (Cranford et al., 1996). These evolutionary differences in form and function may be linked to differences in descending auditory pathways in particular. Application of DTI tractography to mysticetes could provide foundational knowledge on ascending *and* descending auditory pathways of this elusive, understudied parvorder. Furthermore, study of the mysticete auditory system may offer clues as to what features of the odontocete auditory system are specialized for echolocation, as the capability of odontocetes for echolocation is one of the most salient features that differentiates them from the closely related mysticetes.

In the present study, the successful acquisition and diffusion-weighted MRI imaging of an intact sei whale brain (*Balaenoptera borealis*) enabled the first application of DTI tractography to a mysticete. The *B. borealis* specimen was analyzed alongside brains opportunistically collected from odontocetes, including a common dolphin (*Delphinus delphis*), pantropical spotted dolphin (*Stenella attenuata*), and Atlantic white-side dolphin (*Lagenorhynchus acutus*). In order to establish foundational knowledge on the structural connectivity of mysticete brains, as well as search for differences from odontocetes that may shed light on the neural basis of the closely-related parvorder’s echolocative capabilities, this study employed probabilistic tractography to compare the qualitative features, tract strength, and lateralization of potential ascending and descending auditory paths in mysticetes and odontocetes.

## Results

### Whole brain metrics: Volume and fractional anisotropy

The whole brain volume of *D. delphis* was 1,141.51 cm^3^, while *S. attenuata* was 933.20 cm^3^, *L. acutus* was 1067.08 cm^3^, and *B. borealis* was 2,864.37 cm^3^ (Table 1). The inferior colliculi make up 0.28% of the total brain volume in *D. delphis*, 0.37% in *S. attenuata*, 0.34% in *L. acutus*, and 0.28% in *B. borealis* (Table 1). The cerebella make up 17.47% of the total brain volume in *D. delphis*, 16.54% in *S. attenuata*, 12.74% in *L. acutus*, and 13.21% in *B. borealis* (Table 1). There were no grossly obvious differences between the relative volumes of the right and left inferior colliculi or cerebella in any of the animals studied. In *D. delphis*, the mean fractional anisotropy (FA) value was 0.093; in *S. attenuata*, mean FA was 0.100; in *L. acutus,* 0.123; and in *B. borealis,* 0.170 (Table 1).

**Table 1:** Mean FA and volumetrics. Mean fractional anisotropy (FA), whole brain volume (WBV), percentage of whole brain volume contained in inferior colliculi (IC / WBV) and percentage of whole brain volume contained in cerebella (Cerebella / WBV) for each specimen.

|  | *D. delphis* | *S. attenuata* | *L. acutus* | *B. borealis* |
| --- | --- | --- | --- | --- |
| *Mean FA* | 0.093 | 0.100 | 0.123 | 0.170 |
| *WBV* | 1,141.51 | 933.20 | 1,067.08 | 2,864.37 |
| *IC / WBV* | 0.28% | 0.37% | 0.34% | 0.28% |
| *Cerebella / WBV* | 17.47% | 16.54% | 12.74% | 13.21% |

### Tract strength

Each tractography output, abbreviated in this paper as “trace,” included a *waytotal* value denoting the number of streamlines that satisfied the tractography algorithm by originating in the seed region of interest (ROI) and met waypoint and/or exclusion criteria (Thaploo et al., 2022). We then calculated the absolute tract *strength* of each trace by dividing its waytotal by the total brain volume of the respective specimen (Fig 1).

**Fig 1.**
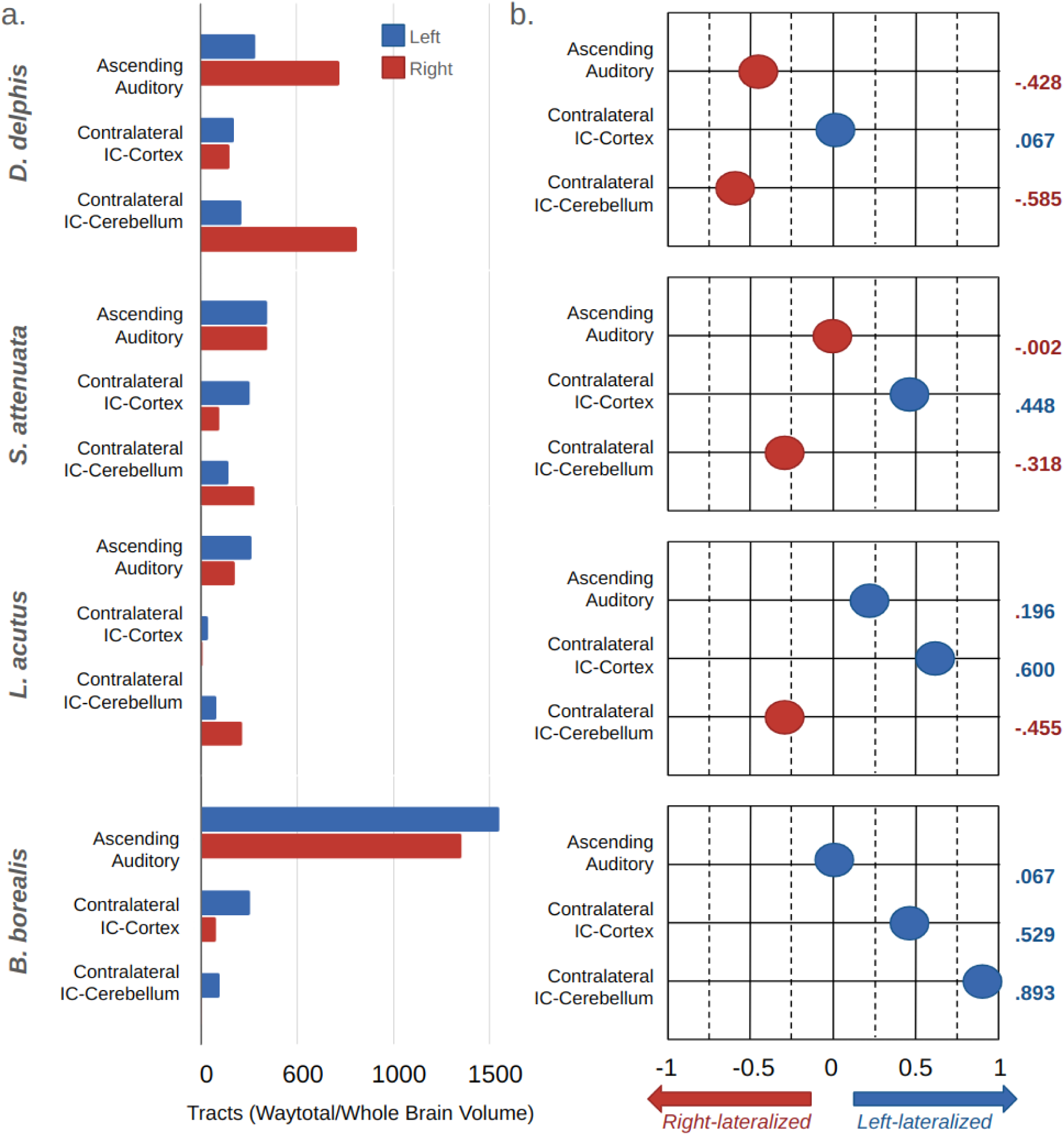
Tract strength and laterality index. Panel (a) contains bar graph representations of the corrected tract strength of each trace in each specimen. See Table 2 for exact values. Panel (b) contains graphical representations of the laterality index (LI) of each trace in each specimen (LI formula and LI graph design adapted from Wright et al. (2018)). LI values range from −1, maximally right-lateralized, to 1, maximally left-lateralized, and are given to the right of each graph. IC= inferior colliculi.

**Table 2.** Tract strength and laterality factor. Tract strength (i.e., waytotal corrected by whole brain volume) and lateralization factor (i.e., how many times more tracts were observed in one hemisphere versus the other) for each trace in each specimen. Ipsi= ipsilateral, Contra= contralateral, IC= inferior colliculi, and LF= lateralization factor. Right versus left designations denote which cortical or cerebellar hemisphere was *targeted* more strongly by a respective inferior colliculus, rather than which the side of the IC was seeded.

|  | *D. delphis* |  |  | *S. attenuata* |  |  | *L. acutus* |  |  | *B. borealis* |  |  |
| --- | --- | --- | --- | --- | --- | --- | --- | --- | --- | --- | --- | --- |
|  | Left | Right | LF | Left | Right | LF | Left | Right | LF | Left | Right | LF |
| IC - Ipsi. Cortex | 287.911 | 719.000 | 2.497x Right | 349.273 | 350.586 | 1.004x Right | 262.506 | 176.421 | 1.488x Left | 1546.153 | 1352.539 | 1.143x Left |
| IC - Contra. Cortex | 175.296 | 153.161 | 1.145x Left | 257.329 | 98.088 | 2.623x Left | 37.535 | 9.374 | 4.004x Left | 253.613 | 78.065 | 3.249x Left |
| IC - Contra. Cerebellum | 212.354 | 811.316 | 3.820x Right | 145.350 | 281.023 | 1.933x Right | 79.636 | 212.374 | 2.667x Right | 92.614 | 5.213 | 17.767x Left |

Among all specimens, the greatest within-specimen tract strength was observed in IC to ipsilateral cortex traces, with the exception of *D. delphis*, in which the left IC-right cerebellum trace showed the strongest within-specimen tract strength. Among all four specimens, the greatest *bilateral* tract strength, and the greatest tract strength overall, was observed in the IC to ipsilateral cortex traces of *B. borealis*, where the volume-adjusted tract count for left and right side traces totaled 1546.15 and 1352.53, respectively. Between all specimens, the weakest observed tract was between the left IC and right cerebellum in *B. borealis*. After dividing by whole-brain volume, we found a mere 5.21 pathways in this trace for *B. borealis*, as opposed to the 811.31 calculated for the same trace in *D. delphis*. With the exception of the left IC to right cerebellum trace in *B. borealis*, the weakest within-specimen tract strength was found in contralateral IC to cortex traces. The weakness of contralateral IC to cortex traces was particularly pronounced in the *L. acutus* specimen, though notably, the tract strength observed in this specimen generally tended to be lower than the other specimens.

### Quantitative lateralization

The *laterality* of each trace was quantified by two methods: simple division of the larger waytotal by the smaller, and calculation of the *laterality index* for each (Wright et al., 2018; Vernooij et al., 2007) (Table 2, Fig 1). In the following, “right-lateralized” versus “left-lateralized” designations reflect which cortical or cerebellar hemisphere was *targeted* more strongly by a respective inferior colliculus, rather than designating whether right or left IC were seeded.

In simple traces of IC connectivity, no lateralization effects were observed (S8 Tables), in accordance with the lack of volumetric differences found between left and right IC in any of the specimens (Table 1). Followup traces examining connectivity between each IC and respective ipsilateral cortex– i.e., excluding both cerebella and the respective cortical hemispheres contralateral to each IC– yielded only mild lateralization effects in all specimens other than *D. delphis* (Table 2). In *D. delphis*, a more notable lateralization effect was observed, with 2.50 times more connections found between the right IC and its ipsilateral cortex than the left (Table 2). In traces between each IC and its respective *contralateral* cortex– i.e., with both cerebella excluded and the sagittal-plane midpoint of each brain set as a waypoint–a common pattern of quantitative lateralization was found in all specimens, echolocator or not (Table 2). Comparatively robust contralateral cortical connectivity of the right IC was observed in all specimens, most strongly in *L. acutus* and *B. borealis*, more moderately in *S. attenuata*, and most weakly in *D. delphis* (Table 2). In *L. acutus*, the contralateral connectivity of the right IC to left cortex was 4.00 times greater than the left IC’s connectivity to right cortex (Table 2).

The strongest lateralization effects were found in traces between inferior colliculi (IC) and the respective cerebellar hemispheres contralateral to each (Table 2). In these traces, we observed a lateralization pattern that was consistent between odontocetes yet robustly reversed in the mysticete (Table 2). Indeed, the most strongly lateralized value in this dataset comes from the *B. borealis* specimen that displayed connectivity between its right IC and left cerebellum that was 17.77 times stronger than between its left IC and right cerebellum (Table 2). By contrast, all odontocetes exhibited greater connectivity between their left IC and right cerebellum (Table 2). In *D. delphis*, this right-lateralization effect was most pronounced, with 3.82 times more connections found between left IC and right cerebellum than right IC and left cerebellum (Table 2). Meanwhile, in *L. acutus* and *S. attenuata*, left IC-right cerebellar tracts were 2.67 and 1.93 times stronger, respectively, than their contralateral counterparts (Table 2).

### Qualitative features

#### Ascending auditory tracts

In all specimens, we found bilateral pathways from inferior colliculi (IC) to temporal lobe that passed through a ventral-caudal portion of the thalamus, likely the medial geniculate nucleus (Figs 2a-d, Fig 3).

**Fig 2.**
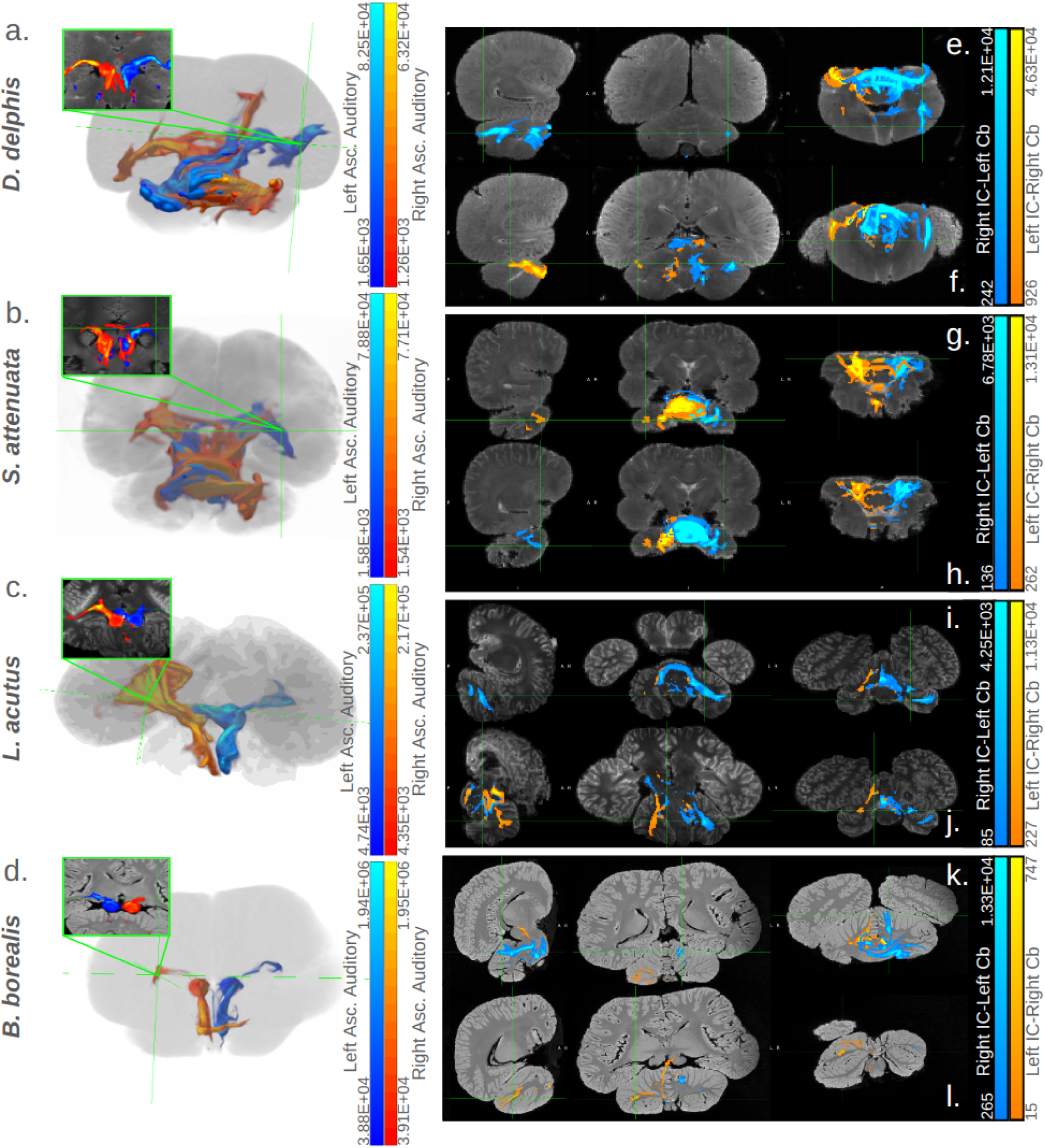
Cortical and cerebellar targets of inferior colliculi. Panels (a)-(d) contain 3D representations of ascending auditory tracts for each specimen. Tracts seeded in the right inferior colliculi (IC) are colored red, while those seeded in the left are colored blue. Adjacent to the 3D view, bright green boxes display coronal cross-section views of these traces in each specimen, highlighting their path through the ventral thalamus (putative medial geniculate nucleus; for a more detailed view see Fig 3.) Panels (e)-(l) contain orthographic views of some main cerebellar targets of traces between the IC and cerebella in each specimen. Tracts targeting the right cerebellum are colored orange, while those targeting the left are colored turquoise. For each specimen, two orthographic-view images are given: one highlighting a prominent projection site in the left cerebellum, the other highlighting a prominent projection site in the right. IC= inferior colliculi, Cb= cerebellum. For more details on cerebellar projection sites, see Fig 6, S9 Table, S1 Text, S6 Figures or S7 Figures. In all panels, the correspondence between the number of predicted tracts and the colors displayed is given in the gradient color bar on the right edge, with brighter colors indicating a higher number of tracts predicted in a given voxel and vice versa.

**Fig 3.**
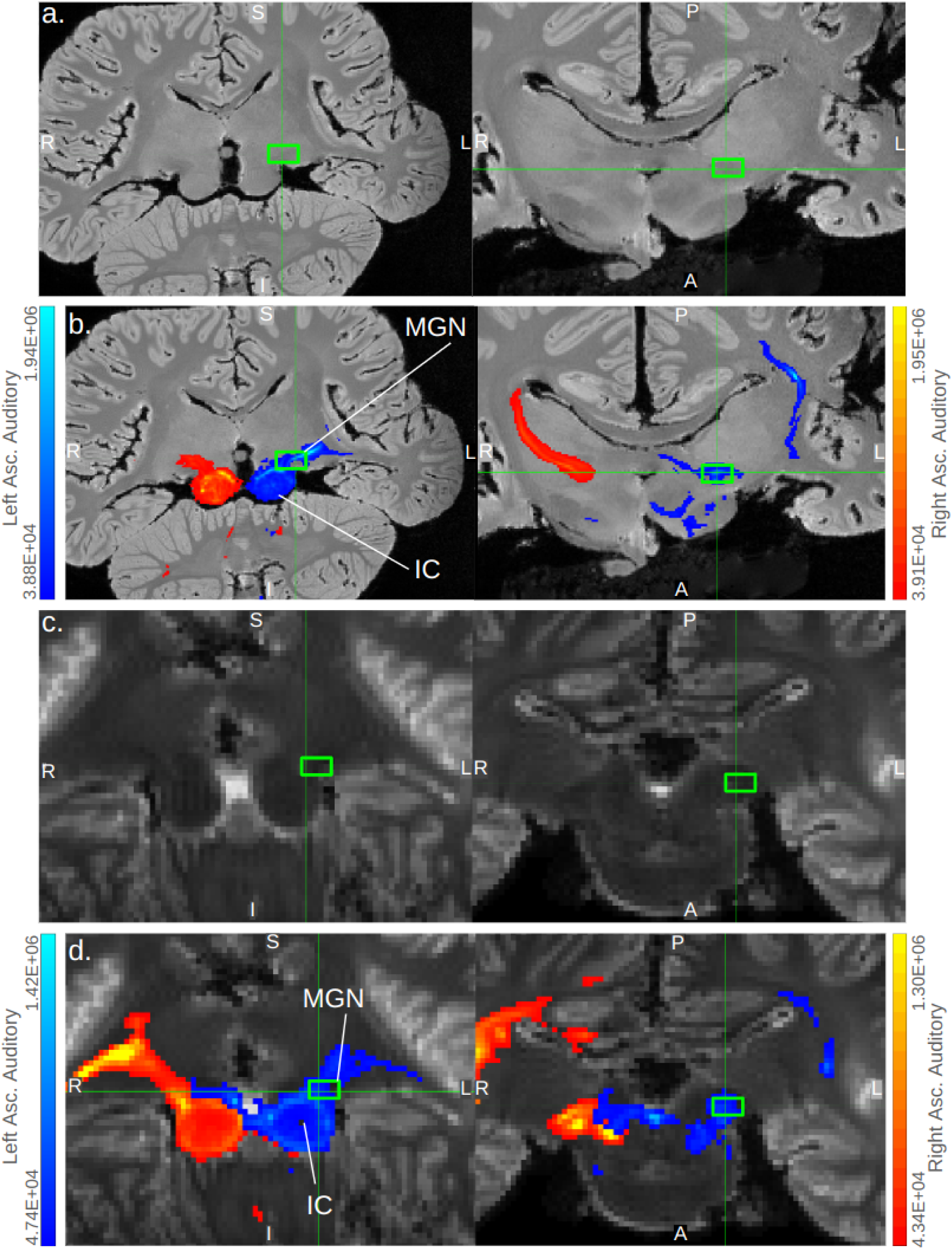
Ascending auditory pathways transit putative MGN. Coronal and axial cross-sections depicting the ventral thalamic transit of ascending auditory pathway traces in *B. borealis* (a, b) and *L. acutus* (c, d). Cross sections without (a, c) and with (b, d) tracts are shown; tracts are thresholded with minimum and maximum values of 0.1% and 5% of the trace waytotal, respectively, in *B. borealis* (b) and with minimum and maximum values of 1% and 30% of the trace waytotal, respectively, in *L. acutus* (d). The correspondence between predicted number of tracts in a given voxel and displayed colors of tracts is given in the gradient color bar on the right edge of panels b and d. Bright green boxes mark putative medial geniculate nucleus in each panel. MGN= medial geniculate nucleus and IC= inferior colliculi.

In *B. borealis*, left and right IC tracts proceeded from the deep temporal lobe adjacent to the IC to an equally ventral but more caudal-medial region of cortex in a fairly linear projection (Fig 3b). In all odontocete IC traces, this deep temporal lobe to ventro-medial-caudal cortical projection was mirrored to at least some extent, most robustly so in *S. attenuata* (Fig 4d). However, the odontocetes also showed multiple other cortical projection sites in addition to the sole ventro-caudal one observed in the mysticete (Fig 5). Consistently, all odontocetes showed bilateral projections from the IC-adjacent deep temporal lobe to a more rostral though equally ventral cortical region (Fig 4b). In *D*. *delphis* in particular, these rostro-ventral projections were robust, particularly on the left side (Fig 4b). These rostral-ventral tracts appeared to reach the caudate head in all odontocetes, and while the left side of the caudate was more strongly implicated across all three, in *S. attenuata*, the contralateral right IC appeared to target the left caudate more prominently than the left IC (Fig 4b). In *L. acutus*, at a fork between the rostral-bound and caudal-bound projections from the deep temporal lobe, a third type of projection ran perpendicular to these toward a more lateral region of cortex, most robustly in the right hemisphere (Fig 4a). In *D. delphis*, these lateral projections were also seen, though more prominently in the left hemisphere (Fig 4a). Additionally, a dorsal-rostral region in the left cerebral cortex of *D. delphis* exhibited uniquely robust projections from both left and right IC, though the projections from the contralateral right IC were particularly pronounced (Fig 4c).

**Fig 4.**
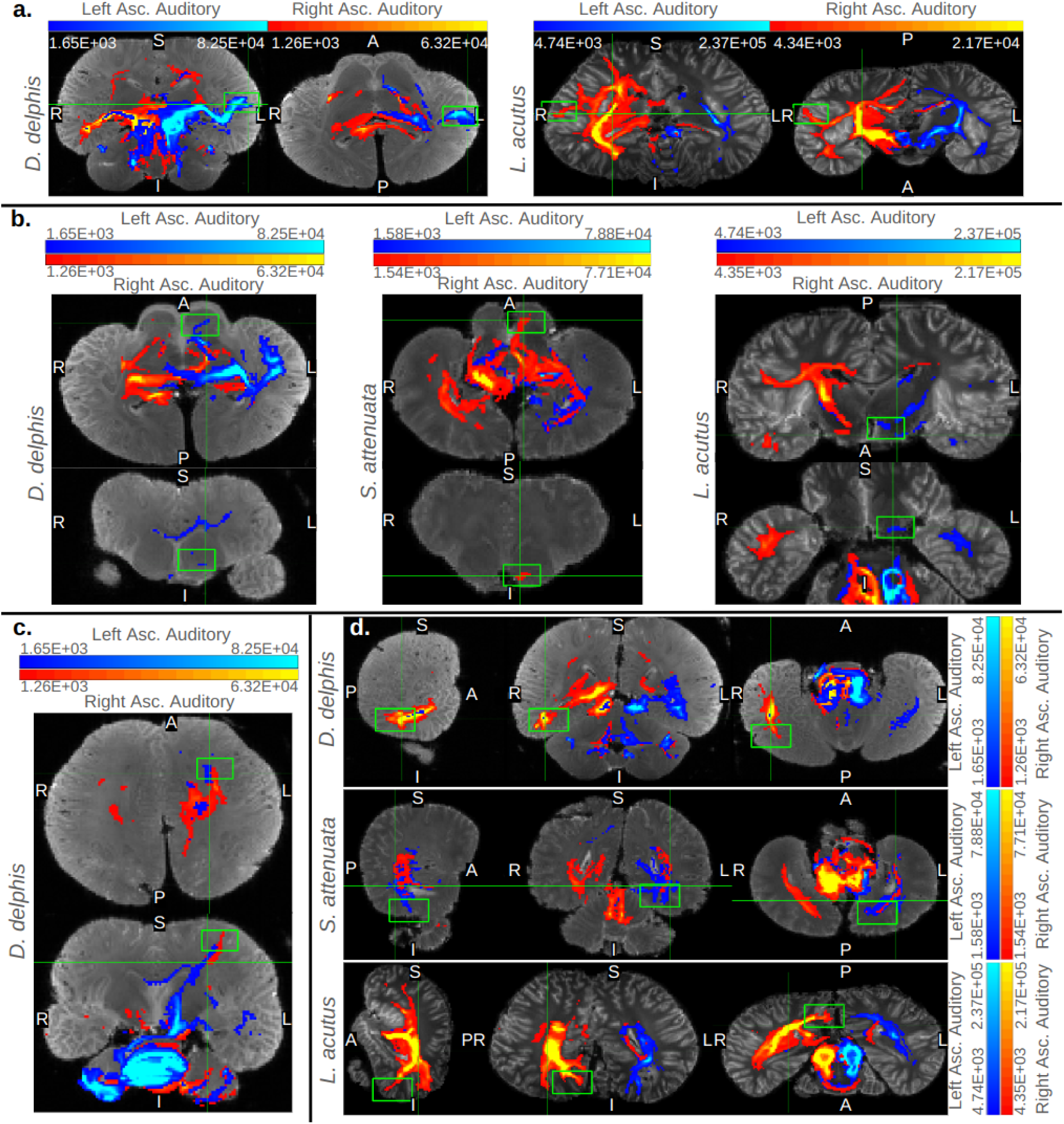
Cortical targets of ascending auditory traces in odontocetes. Orthographic cross-sections displaying key cortical targets of the inferior colliculi in the odontocete specimens. Each trace is displayed with minimum and maximum values set to 0.1% and 5% of its waytotal, respectively. In all panels, traces from right IC are colored red and those from left IC are colored blue. Gradient color bars specify the correspondence between predicted number of tracts in a given voxel and displayed colors of tracts. Panel (a) displays coronal and axial views of projections to lateral extremes of the left and right cortex in *D. delphis* and *L. acutus*, respectively. Panel (b) displays axial and coronal views of tracts transiting the basal ganglia, potentially caudate nucleus, in all odontocetes. Panel (c) displays axial and coronal views of continuous projections between both colliculi and a dorsal-rostral region of left cortex in *D. delphis*. Finally, panel (d) displays sagittal, coronal, and axial views of projections to a caudal, ventral, and medial cortical area in all odontocetes. Bright green boxes mark the notable cortical projection sites mentioned in-text.

**Fig 5.**
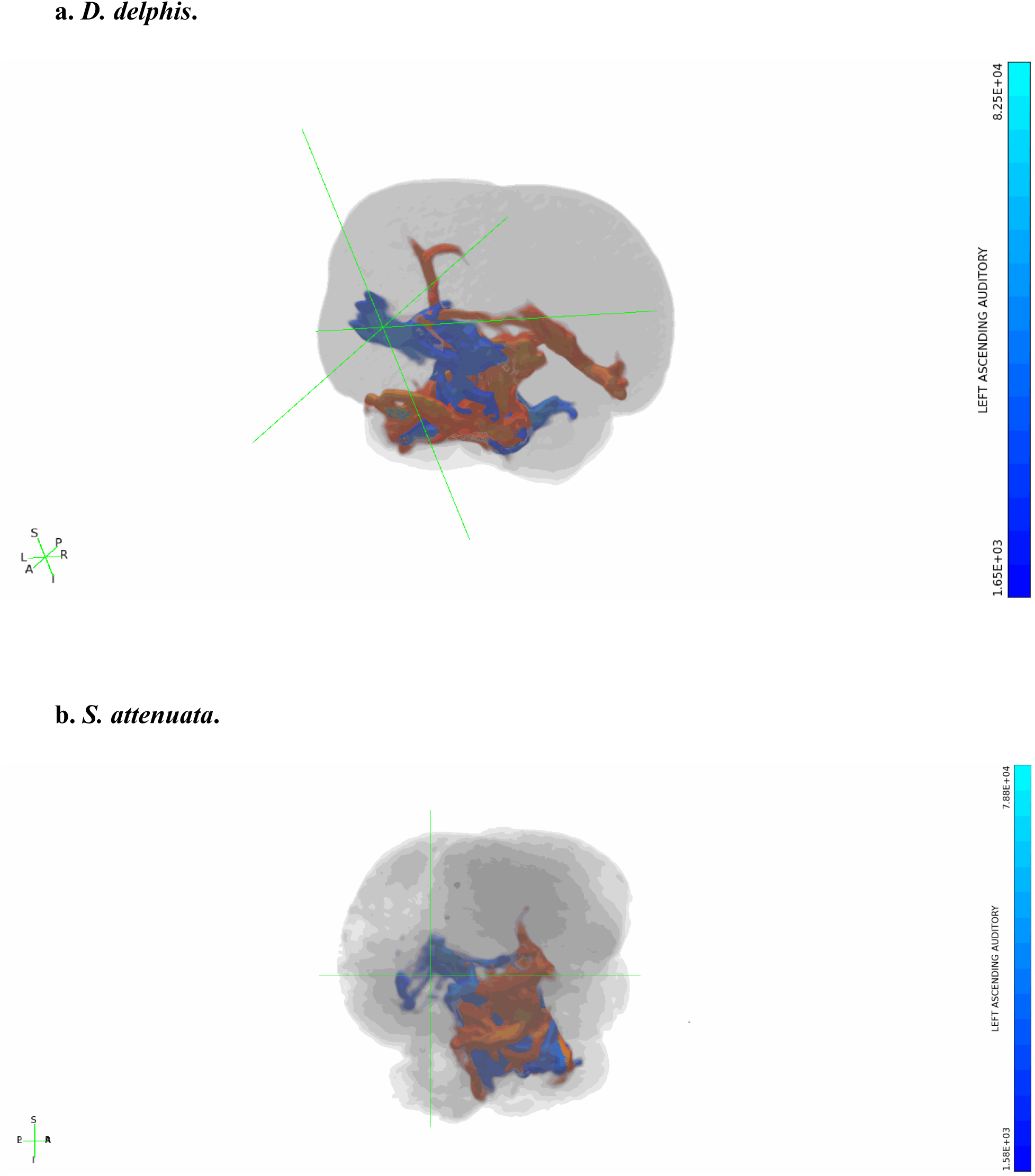
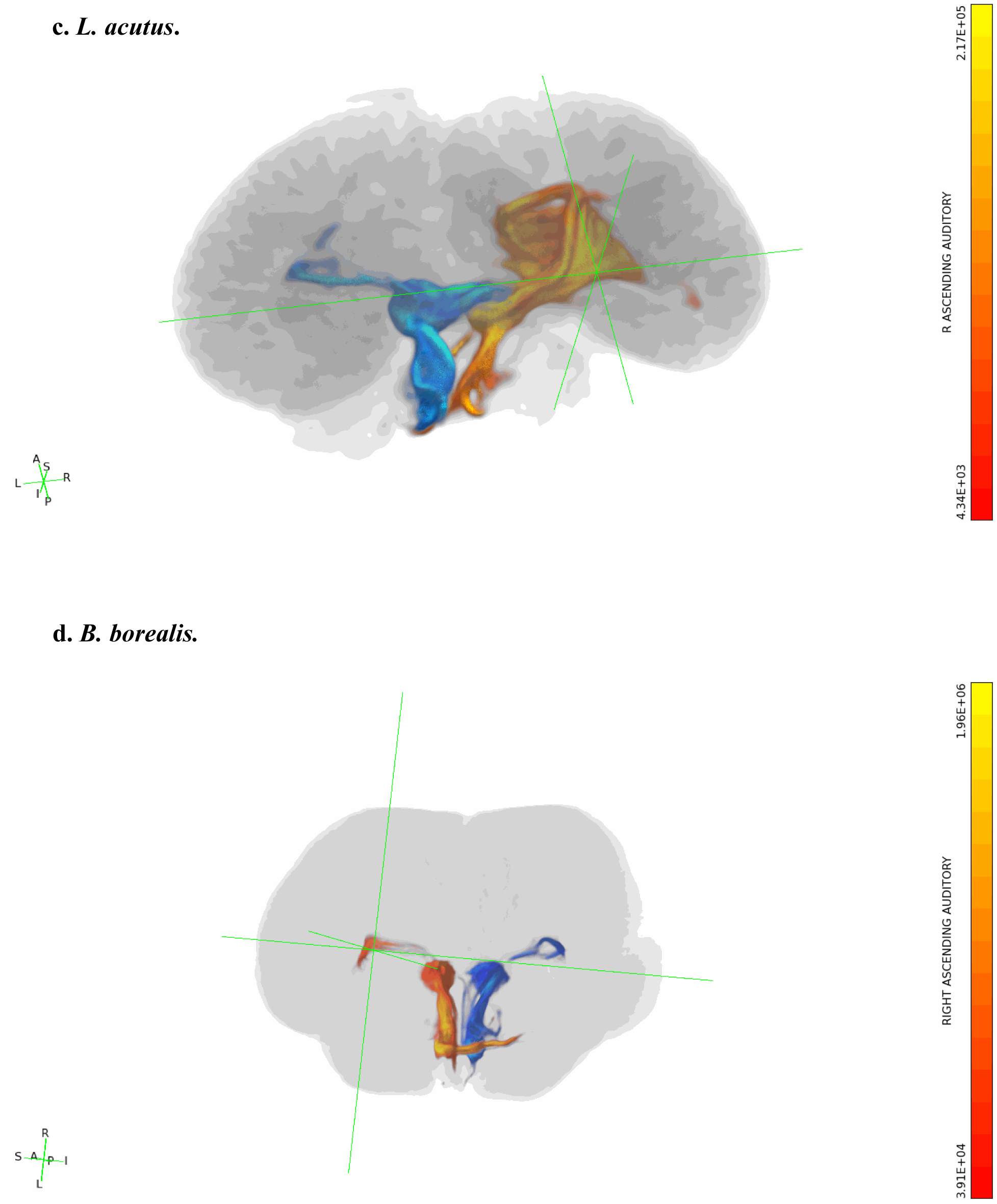
a-d. Rotating 3-dimensional tractograms of ascending auditory pathways in all specimens. All displayed tractograms were seeded in the IC (refer to Fig 9 or S3 Figures to view the masks of IC that served as seeds). These traces are displayed with minimum and maximum values set to 0.1% and 5% of their waytotals, respectively. In all panels, traces from right IC are colored red and those from left IC are colored blue. Gradient color bars specify the correspondence between predicted number of tracts in a given voxel and displayed colors of tracts.

#### Contralateral collicular-cerebellar tracts

We identified cerebellar projection sites for the IC to cerebellum traces in each specimen, using the cerebellar divisions and nomenclature established in Schmahmann et al. (1999) and subsequently applied to a dolphin brain in Hanson et al. (2013) as anatomical guidelines. A summary of the findings is provided below in-text, as well as in Fig 6 and S9 Table, in which the putative functions of each projection site are further specified. For more detailed descriptions of the subcortical and cerebellar targets of the IC-cerebellar traces, refer to S1 Text.

**Fig 6.**
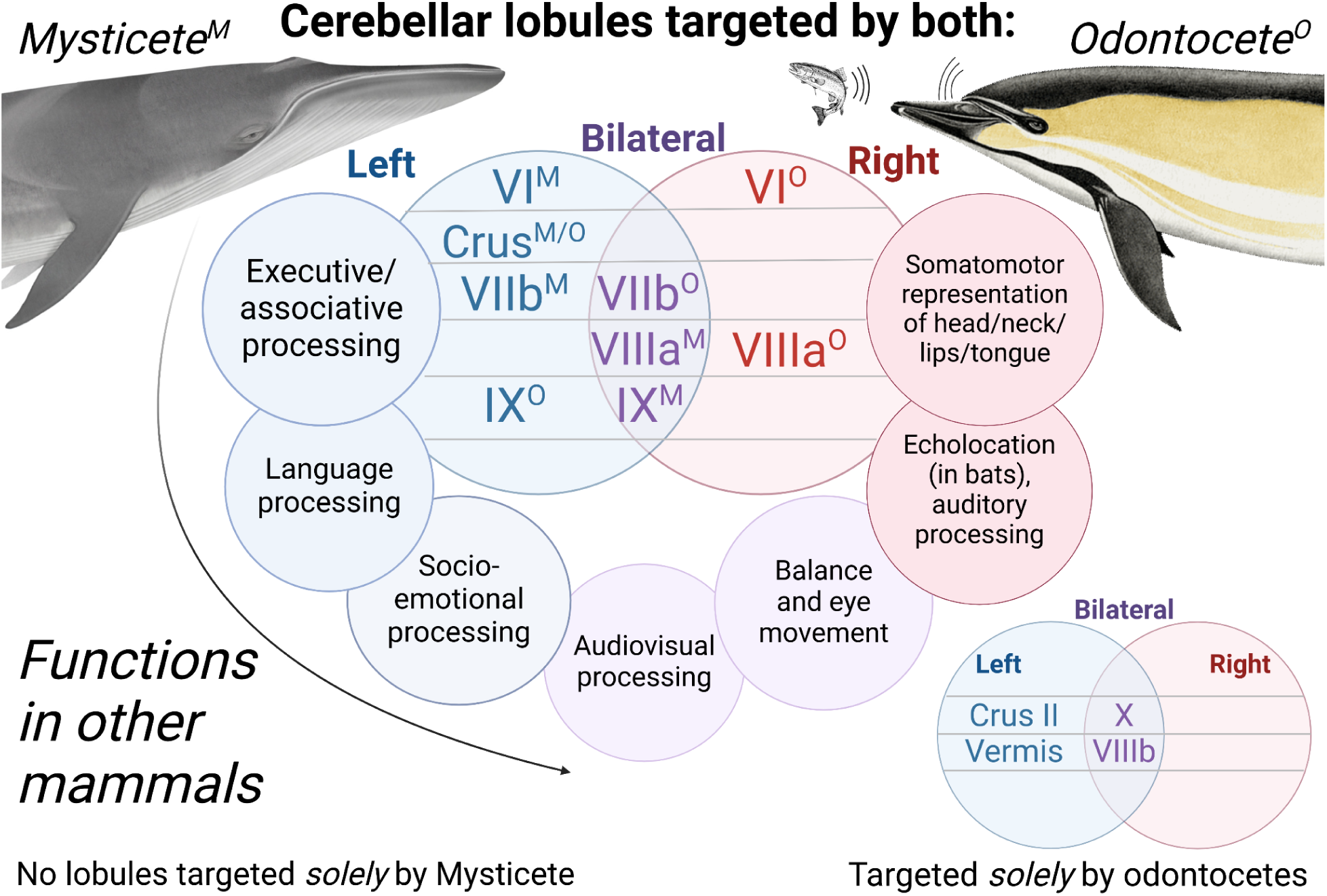
Color-coded diagram displaying cerebellar lobules targeted in the left, right, or bilateral cerebella in odontocete and mysticete specimens. Within the central venn diagram, superscript “M” signifies that a subregion was targeted in the mysticete, while “O” signifies a subregion’s targeting by one or more odontocetes. Subregions that were targeted in the left cerebellar hemisphere are placed in the blue circle on the left, while right-hemisphere subregions are placed in the red circle on the right, and bilaterally-targeted subregions are placed in the overlap region colored purple. The outer ring of the central venn diagram specifies the functions associated with each lobule in other mammals (Azizi et al., 1985; Brissenden et al., 2018; Burne & Woodward, 1983; Cozzi et al., 2018; Hanson et al., 2013; Suga & Horikawa, 1986; O’Reilly et al., 2009; Kirschen et al., 2010; Klein et al., 2016; Metoki et al., 2022; Nakatani et al., 2022; Van Overwalle et al., 2020; Snider & Stowell, 1944). Each function is placed in proximity with the lobule to which it corresponds in the inner venn diagram, and is color-coded in accordance with the subregion to which it corresponds as well. See the discussion subsection Specific cerebellar targets for more details on correspondences between subregions and functions. In the bottom corner, a smaller venn diagram displays the subregions only targeted in odontocetes, and uses the same color-coding scheme to denote whether these regions were targeted on the left side, right side, or bilaterally.

In all specimens, IC to cerebellar traces tended to produce bilaterally robust pathways through the auditory brainstem regions between and proximal to the pons and cerebellum, including the olivary complex, the trapezoid body, and the lateral lemniscus (Figs 2e-l, Fig 7). In all of the dolphins, multiple distinct, seemingly transverse pathways emerged rostrally from the more caudal brainstem at different levels of the pons, typically one at a dorsal extreme of the pons, one at a ventral extreme of the pons, and a third medial to these two in the dorsal-ventral axis (Fig 7, S1 Text). In *S. attenuata*, these three parallel transverse pontine pathways, as well as the dense brainstem pathways caudal to them, tended to be fairly bilateral (Fig 7b, S6 Figures B1-4). Indeed, more so than in any other specimen, both IC to cerebellar traces in *S. attenuata* displayed prominent dorsal-ventral projections through the mylencephalon, possibly traversing the inferior cerebellar peduncle, that proceeded ventrally and laterally from the floor of ventricle IV on the caudal brainstem, potentially contacting the facial colliculi situated here, to the caudal border of the inferior olives and eventually the rostral-most portion of the spinal cord (Fig 7b). Interestingly, though, the left IC to right cerebellum trace, but *not* the right IC to left cerebellum trace, produced a second, similarly continuous and robust projection between the floor of ventricle IV and the caudal edge of the inferior olives, medial and slightly rostral to the bilateral parallel projections described above, closer to the facial motor nucleus (Fig 7b).

**Fig 7:**
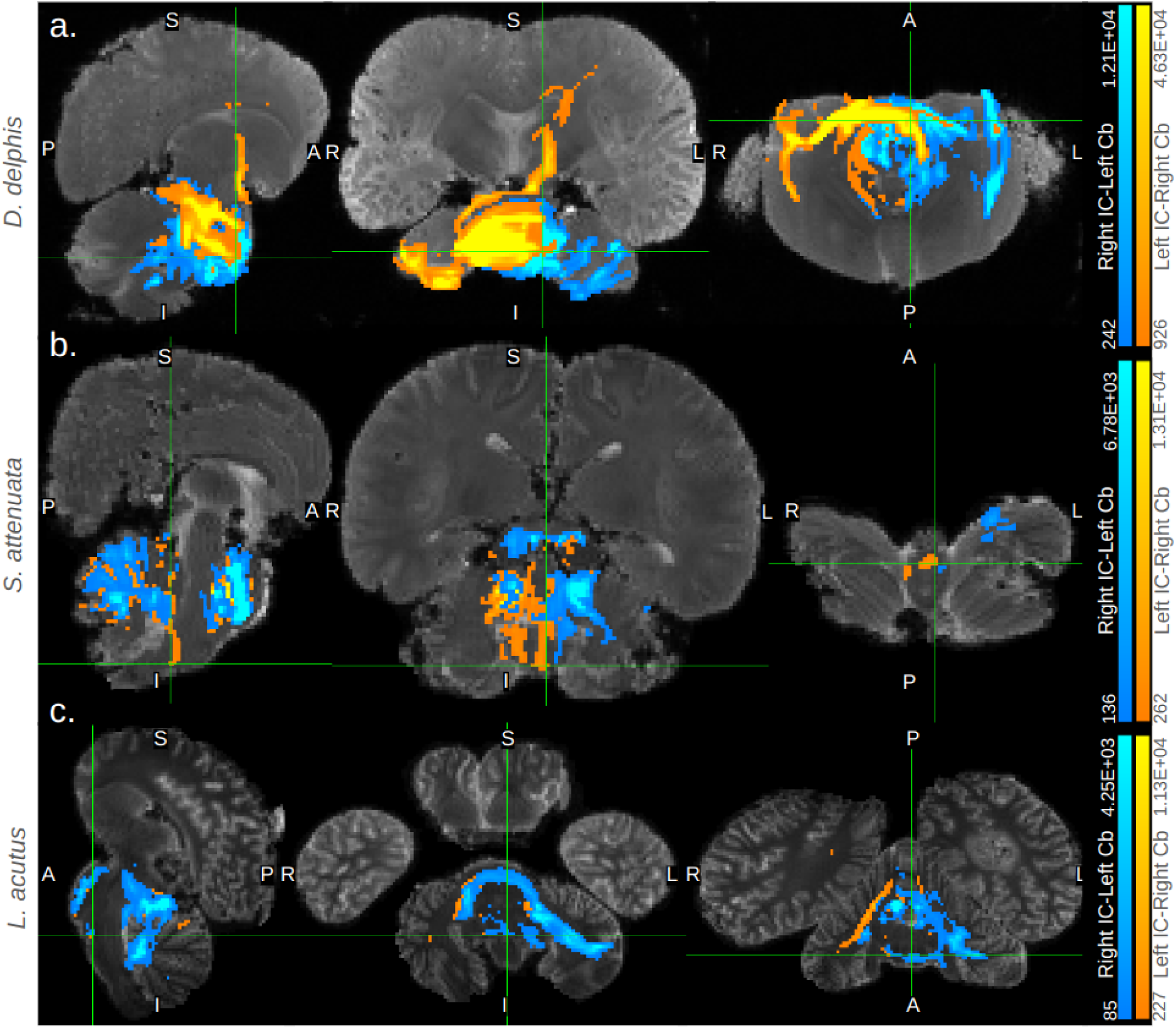
Subcortical features of IC-cerebellar traces in odontocetes. Orthographic cross-sections displaying key subcortical features of the traces between IC and cerebellum in the odontocete specimens (a) *D. delphis*, (b) *S. attenuata*, and (c) *L. acutus*. Each trace is displayed with minimum and maximum values set to 0.1% and 5% of its waytotal, respectively. Traces between right IC and left cerebellum are colored blue, while those between left IC and right cerebellum are orange. The correspondence between predicted number of tracts in a given voxel and displayed colors of tracts is given in the gradient color bars on the right edges of the panels. IC= inferior colliculi and Cb= cerebellum.

In the other dolphins, such instances of qualitative lateralization were often apparent. Strikingly, in the left IC to right cerebellar trace of *D. delphis*, the most dorsal of the pontine pathways continuously connected a dorsal-rostral portion of the right cerebellum, likely in lobule IX, with a dorsal-rostral extreme of the left cortex, passing through rostral extremes of diencephalic and mesencephalic structures such as the crus cerebri and substantia nigra in between (Fig 7a, S6 Figures A1-4). Meanwhile, in *L. acutus*, right IC to left cerebellum traces tended to produce more bilateral tracts caudal to the pons in the auditory centers of the brainstem, appearing on both right and left sides of the brainstem and cerebellar peduncles, while the left IC to right cerebellar traces remained more concentrated on the right side (Fig 7c). Compared to the dolphins, in *B. borealis*, these pontine pathways tended to be slightly more ventrally-shifted, and were more robust in right IC to left cerebellar traces as opposed to left IC to right cerebellar ones (Fig 2k-l, S6 Figure D1, S7 Figure D2).

In the cerebellum itself, a handful of subregions emerged as the most relevant targets of IC-cerebellar traces among all specimens. The most prominent regions were Crus I and lobule IX, which received projections from IC-cerebellar traces in all four specimens (S9 Table). Crus I showed consistent patterns of lateralization, with left Crus I implicated in right IC to left cerebellar traces in both the mysticete and odontocetes (S1 Text, S9 Table, Fig 6, S7 Figures). Lobule IX was bilaterally implicated in the mysticete, while in the dolphins, the right IC to left cerebellum traces more prominently targeted left lobule IX (S1 Text, S9 Table, Fig 6, S7 Figures). The next most prominent subregions were lobule VIIIa and Crus II, with each receiving projections from three of the four specimens (S9 Table). Crus II was only targeted in the odontocetes and *not* in the Mysticete, and in all odontocetes, this was observed *only* in the right IC to left cerebellar traces, i.e. in left Crus II (S1 Text, S9 Table, Fig 6, S7 Figures). Lobule VIIIa was targeted in all specimens except *S. attenuata*, and was bilaterally implicated in *B. borealis* and *L. acutus*, while *D. delphis* exhibited right lobule VIIIa projections only (S1 Text, S9 Table). Finally, lobules VI, VIIb, VIIIb, and the vermis were all observed as prominent targets in two of the four specimens (S1 Text, S9 Table). Lobule VI was implicated in the left cerebellum of the mysticete, but in the right cerebellum of one odontocete (S1 Text, S9 Table). Meanwhile, lobule VIIb received bilateral targeting in the mysticete, but only received projections in the right cerebellum of its odontocete representative, *D. delphis* (S1 Text, S9 Table). By contrast, lobule VIIIb was only targeted in odontocete specimens, and in both of these was bilaterally implicated (S1 Text, S9 Table). Likewise, the vermis only received projections from IC to cerebellar traces in odontocete specimens, though in *L. acutus* the vermis was targeted more by right IC to left cerebellar traces as opposed to bilaterally (Fig 7c, S1 Text, S7 Figures C1-2). Notably, anterior lobules I-V were not meaningfully targeted by IC-cerebellar traces in any of the specimens, and lobule X was prominently and bilaterally targeted, with both cerebella receiving projections from both IC, in *D. delphis*, but was not pronouncedly targeted in any other specimens (S1 Text, S7 Figure A1).

In all odontocete specimens, traces between IC and contralateral cerebella included some cortical projections at the more and less stringent thresholds specified above (S6 Figures A1-4, B1-4, and C1-4). In the mysticete, cortical projections were sparse at the liberal threshold and nearly absent at the more conservative one (S6 Figures D1-4). Among the odontocetes, cortical projections appeared weakest in *S. attenuata* and strongest in *D. delphis* and *L. acutus*, and were qualitatively lateralized in all, but most strikingly so in the latter specimens, with more notable projections in the left cortex for *D. delphis* and the right for *L. acutus* (S6 Figures A1-4, B1-4, and C1-4). When compared to the ascending auditory traces, IC to cerebellar traces in *D. delphis* and *L. acutus* tended to exhibit more robust paths to rostral and dorsal portions of cortex, and less prominent connections to ventral and caudal extremes (S6 Figures A1-4 and C1-4). For more details on these cortical projections, see S2 Text or S6 Figures.

## Discussion

In this post-mortem diffusion tractography study of one mysticete and three odontocete brains, we replicated and extended the findings on cetacean auditory pathways from Berns et al. (2015). Further, as suggested by Wright et al. (2018), we assessed these tracts for evidence of lateralization, and found none. In all four species, tract density of ascending cortical auditory pathways was largely symmetrical. We did, however, find evidence of strong asymmetry in descending auditory pathways in all four species. Each of the dolphins showed greater tract density between left inferior colliculus and right cerebellum than vice versa, while the sei whale showed the opposite pattern. This lateralization of descending auditory pathways is intriguing, particularly in light of the lack of lateralization of ascending cortical tracts.

Unlike the mysticetes, which have bilateral laryngeal innervation, the odontocete sound production system separately controls the right and left phonic lips. In the dolphins, increased descending tract strength from left hemisphere to right cerebellum may serve to support the greater predictive sensorimotor processing demands of echolocation as opposed to passive hearing. This accords with a cluster of behavioral findings reporting that the right phonic lips more frequently produce navigational clicks in multiple odontocete species, and may support the resulting functional laterality hypothesis, which posits that echolocation clicks are preferentially produced by the right phonic lips and communication sounds by the left (Ames et al., 2020; Madsen et al., 2013; Ridgway et al., 2015). Although ascending auditory projections in mammals are bilateral, decussating in the brain stem, motor processing is strictly lateralized. Thus the right phonic lips should be controlled by vocal motor regions in the left cortical hemisphere. These, in turn, communicate most directly with auditory regions in the left hemisphere, which communicate most directly with the left inferior colliculus. Cerebellar-cortical communication is contralateral, so descending left cortical auditory signal from the auditory cortex through the IC will target the right cerebellum (Fig 8).

**Fig 8.**
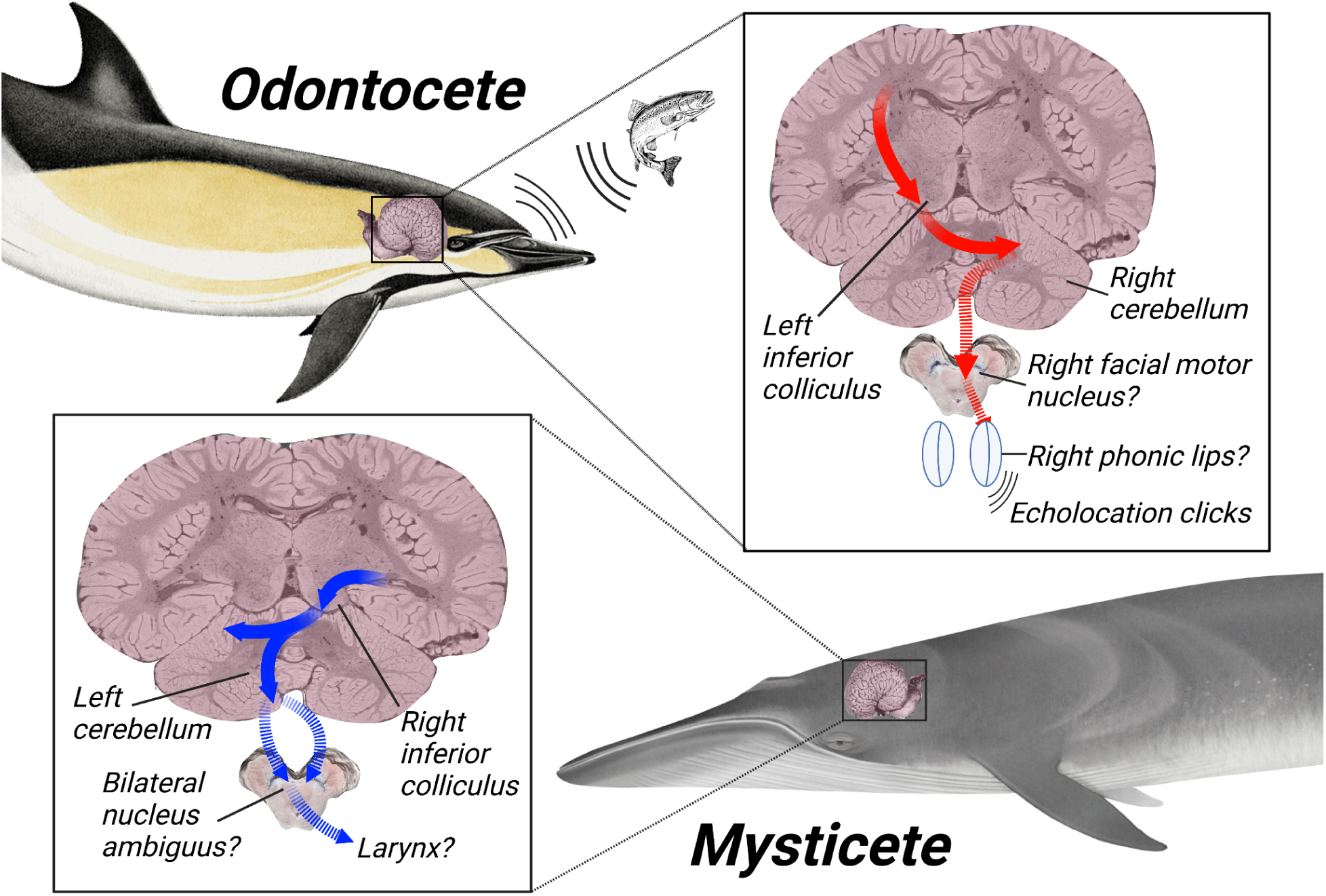
Putative model of differential auditory cortical-cerebellar connectivity related to echolocation processing in odontocetes. Red arrows color-code for right-cerebellar lateralization of these tracts in odontocetes, while blue arrows color-code for left-cerebellar lateralization of these tracts in mysticetes. Brainstem cross-sections display the hypothesized brainstem nuclei targeted by these tracts in each, and the hypothesized peripheral organs that would in turn receive motor signals from these nuclei. Coronal cortical cross-section adapted from Brain Catalogue (Toro, 2015), and brainstem cross-section adapted from Neurosurgical Atlas (Cohen-Gadol, 2024).

Beyond the behavioral findings described under the functional laterality hypothesis, anatomical studies of asymmetries in odontocete nasal musculature and skull also suggest that the right phonic lips may be differentially employed in the generation of echolocation clicks. In many dolphins, the fat-filled melon, which serves to direct and amplify echolocation clicks, is directly connected to the right-side, and not left-side, bursae within the phonic lip complex (Cranford et al., 1996; Harper et al., 2008; McKenna et al., 2011). Further, paleontological studies of cetacean skulls from fossil records dating back millions of years suggest a general tendency toward increasing nasofacial asymmetry among odontocetes (but not mysticetes) as their echolocation capabilities have been more finely honed across successive generations (Coombs et al., 2020). These findings clearly complement the evidence at hand of an asymmetrical hypertrophy in descending acousticomotor tracts that is unique to odontocetes (Table 2). Furthermore, left-side skeletal features are more likely to progressively shift in position, and nasal and melon regions responsible for sound production have shifted the most, suggesting that a progressive expansion of musculature supporting sound production on the right side has occurred at the expense of the left, leading left-side nasofacial landmarks to shift further from the median (Laeta et al., 2023). Especially when overlaid with behavioral reports of preferential use of the right phonic lips in echolocation, such paleontological reports seem to further emphasize that in our results, the robust right-side auditory-cerebellar afferents found in odontocetes may support the generation of echolocation clicks (Fig 8).

### Ascending auditory pathways

Qualitatively, we identified direct projections to temporal lobe from inferior colliculus in a new species of dolphin *(L. acutus*) and, for the first time, in a baleen whale (*B. borealis*). Contrary to results from early invasive electrophysiology (Sokolov et al., 1972; Popov et al., 1986), our findings add further support for a primary auditory field in cetacean temporal lobe, as is found in most terrestrial mammals. Further, we provide the first characterization of auditory brain projections in a large baleen species.

Despite the lack of echolocation capabilities in baleen whales, the relative density of cortical auditory tracts was greater in the sei whale compared to the three dolphin brains. Because sei whales specialize in low frequency calls (Baumgartner et al., 2008), and high frequency auditory processing is likely to be more computationally demanding (Clark et al., 2009), it is unlikely that this difference reflects enhanced auditory processing demands in the sei whale. Despite being the oldest of the specimens, and thus the most susceptible to potential age-related fractional anisotropy (FA) loss (Shepherd et al., 2008), the sei whale had the highest mean FA value (Table 1). This increased FA could be due, in part, to the greater myelination required to support signal transmission over longer distances in a larger brain (Lazari & Lipp, 2021)--though see Beaulieu (2002) for a case against a specific FA–myelin correlation. If the heightened FA found in the sei whale indeed reflects greater myelination in its brain overall, it is possible that such a global tendency could affect our measures of tract strength– though this interpretation would in turn underscore the notability of our finding that descending IC to cerebellum tract strength in the sei whale was markedly lower than in the lower-FA odontocetes (Fig 1a).

The location and organization of primary auditory cortex (A1) in odontocetes remains a mystery. Before Berns et al. (2015), the general consensus among cetacean neurobiologists was that primary auditory cortex in the dolphin was posterodorsal, running along the suprasylvian gyrus adjacent to primary visual cortex (Oelschlager, 2008, although see Revischin and Garey, 1990). While this dorsal A1 location matches the most precise odontocete electrophysiological data, it would represent a striking adaptation compared to terrestrial mammals, including the dolphin’s closest terrestrial relatives, the artiodactyls, in which A1 is situated in dorsal temporal lobe (Adrian, 1943; Andrews et al., 1990; Plogman & Kruska, 1990). The Berns et al. (2015) findings were the first to directly support a typical mammalian dorsal temporal location for dolphin A1. We have now replicated those findings with a third dolphin species. Our parallel finding of temporal projections in the mysticete provide further evidence that cetacean A1 may be conserved from the shared common terrestrial ancestor, not having diverged during the evolution of auditory mechanisms supporting echolocation following the primary odontocete split. However, it must be emphasized that our diffusion tractography results do not match the earlier electrophysiological data. There are unlikely to be new invasive electrophysiological data collected in these species, so other lines of evidence are required to address this apparent conflict. Initial work with functional near infrared spectroscopy measures suggests that, with a smaller odontocete such as a porpoise, it might be possible to access neural signal in parts of the cortex (Ruesch et al., 2023). While logistically difficult, functional MRI with dolphins is also a possibility (Ridgway et al., 2006). Histological analyses of putative auditory cortices may be more immediately instructive. In a recent paper examining the putative posterodorsal A1 location, Graic et al. (2023) identified a number of unusual cytoarchitectural characteristics that do not neatly match what is known about cellular organization of A1 in terrestrial species. Notably, the region did not have the standard koniocortical organization of mammalian sensory fields. In addition, the organization of the region was less complex than the putative dolphin primary visual region. The authors suggested this might be due to some of the general whole-brain cytoarchitectural differences between odontocetes and terrestrial mammals, such as reduced or missing layer IV in dolphins and distribution of cortical processing modules over larger less densely packed layers of cortex. However, the findings are far from conclusive. Future work should at least examine cytoarchitecture in the putative dorsal temporal A1 location as well.

One possibility, raised first in Berns et al. (2015) and echoed in Ridgway et al. (2015), is that, similar to bats, dolphins may have separate cortical regions for auditory processing, including a conserved dorsal temporal region for tonotopic response and an enhanced posterodorsal cortical region for processing spatial characteristics of sound related to echolocation, including direction and delay. In the current study, lower-threshold images revealed that IC projections to cortical regions besides the dorsal temporal lobe were indeed present in the echolocating odontocetes, though less so in the non-echolocating mysticete, lending some credence to this hypothesis (S4 Figures). However, the posterior dorsal belt of suprasylvian cortex highlighted in past literature on putative cetacean A1 was not consistently or particularly robustly implicated across the odontocete specimens. Instead, rostro-ventral projections towards the basal ganglia (Fig 4b), dorso-medial projections towards putative cingulate and somatomotor cortices (Fig 4c), and branches to a ventrocaudal extreme of cortex (Fig 4d) were more frequently observed. Future diffusion tractography studies should further examine any putative secondary auditory projections and corticocortical connectivity with dorsal temporal lobe. In Wright et al. (2018), a putative arcuate fasciculus was identified in both hemispheres with deterministic tractography. Both left and right tracts passed through the temporal lobe, but, notably, the left hemisphere fasciculus did appear to project more caudally. These tracts were extracted from whole-brain deterministic maps, however, and were not explicitly seeded in either putative A1 location.

If, as suggested by Ridgway et al. (2015), dolphins do indeed privilege social auditory processing in the right cortical hemisphere and echolocation in the left, the lack of lateralization in primary ascending auditory tracts, found in in all three dolphin species and the mysticete analyzed here, is notable. Prima facie, these findings do not appear to parallel the finding of more robust arcuate fasciculus in the dolphin’s right hemisphere (Wright et al., 2018). However, corticocortical fasciculi may be lateralized even when afferent fibers targeting the relevant cortical regions are not. In addition, lateralized ascending auditory-motor connectivity may be more relevant to long-term flexible vocal production learning, as seen in some dolphin vocal communication (Janik, 2014; Janik & Knornschild, 2021; Vernes et al., 2021), and brainstem, midbrain, and cerebellar adaptations may be more relevant to the real-time, high-speed auditory-motor integration required by echolocation. Further, although auditory motor connectivity may be enhanced in the right hemisphere to promote social communication in the dolphin, there are other potential corticocortical connections required to support echolocation and related learning (e.g., multisensory integration to link echolocation with vision). Therefore, our findings here, of no asymmetry in ascending auditory projection strength, contextualize, as opposed to support or contradict, those in Wright et al. (2018).

### Descending auditory pathways

Given the speed of processing required for echolocation, much has rightly been made of the raw size of early auditory nuclei in dolphin brainstem and midbrain (Malkemper et al., 2012; Ridgway et al., 1981; Schulmeyer et al., 2000; Zook et al., 1988). Less work has examined potential contribution of the cerebellum. While transit time from auditory cortex to cerebellum might on first consideration seem too slow to meaningfully contribute to echolocation, the cerebellum appears to operate as a feed-forward Bayesian system for sensorimotor integration and related movement planning (Ebner & Pasalar, 2008; Ohyama et al., 2003; Tanaka et al., 2020). Part of what the cerebellum probabilistically models is the organism’s own subsequent behavior based on the current sensorimotor situation, leaving open the possibility for a strong cerebellar role in both click production and body-movement based on click production and reception. In Bayesian terms, the dolphin’s current sensorimotor situation, comprising body location and movement, vocal motor echolocatory production, and echoic feedback could be integrated with prior knowledge of successful and unsuccessful movements and click productions in similar situations, with the potential to output a feedforward motor plan judged most likely to be successful for the immediate motor goal. This is very much in line with evidence and interpretation from Beedholm et al. (2023), which showed that evoked auditory responses to echolocation clicks happen *after* subsequent echolocation clicks are produced. A cerebellar model of sensorimotor prediction has also been proposed for general mammalian hearing, wherein auditory stimuli are expected as a byproduct of an upcoming motor act (Knolle et al., 2012).

Although data on the functional role of cerebellum in dolphin echolocation are limited, there is strong evidence of cerebellar hypertrophy in the odontocete line (Hanson et al., 2013; Marino et al., 2000). In the present study, the sei whale cerebellum had similar relative brain volume as the Atlantic white-sided dolphin, but the common dolphin and pantropical spotted dolphin cerebella were notably larger (Table 1). Prior evidence indicates that baleen whales and also sperm whales have smaller relative cerebella in comparison to most toothed whales (Ridgway & Hanson, 2014; Ridgway et al., 2017). It appears that the cerebellum does not scale allometrically with brain size in cetaceans. Given the apparent enlargement of auditory regions in the cerebella of echolocating odontocetes, Oelschlager (2008) has suggested there may be a link between cerebellar hypertrophy and echolocation. This would parallel extensive data showing A. relatively larger cerebella in bats compared to non-echolocating bats and terrestrial rodents (Paulin, 1993), and B. the specific physiological involvement of the bat cerebellum in echolocatory processing (Covey, 2005; Jen & Schlegel, 1980; Kamada & Jen, 1990; Sun et al., 1983). Specifically, auditory receptive regions in bat cerebellum have been found to code for range and direction of moving targets, which is likely necessary for the rapid, subcortical feedforward motor decisions required to get where a target is going, as opposed to where it was.

Given the functional laterality hypothesis and its prediction of lateralized neural processing of production and reception of communication vs. echolocation sounds, the fact that primary ascending cortical auditory tracts in the dolphins were not as heavily lateralized as the descending tracts tentatively suggests that, at least in terms of post-IC neural processing, the cerebellum may play a larger role in managing echolocation-specific computation than the literature has previously suggested. In baleen whales, which do not appear to have lateralized sound production mechanisms (Elemans et al., 2024), it remains to be seen to what extent the functional laterality hypothesis applies, if at all. In the current study, the lateralization pattern of auditory cerebellar pathways was reversed in the sei whale in comparison to the dolphins, with far denser tracts entering left cerebellum than right. However, the fact that *left* tract density was roughly the same in all four species suggests that hypertrophy of the right tract may drive the asymmetry seen in the dolphins, *not* a reduction of left cerebellar auditory processing. Why, then, is the sei whale’s right-to-left descending auditory tract so much stronger than its right? Baleen whales, just like odontocetes, show right side bias for social and feeding behavior, which has been suggested to be rooted in left-hemisphere sensory biases (Canning et al., 2011; Karenina et al., 2016), so the sei whale’s left-cerebellar lateralization is unlikely to be due to primary sensorimotor processing. The cerebellum is a multipurpose structure, and serves many divergent and integrative functions. One possible explanation is that, if, as in the dolphins, the right cortical hemisphere is specialized for communication in baleen whales, the left cerebellum may play an important role in production of vocal communication signals. There are bird data showing the importance of the cerebellum to vocal learning (Pidoux et al., 2018), and there is evidence in humans that the cerebellum contributes meaningfully to language processing (Murdoch, 2010; Nakatani et al., 2022). It is plausible that a right cortical-left cerebellar circuit for social vocal learning is present in both toothed and baleen whales. One interpretation of our current findings is that while all whales, toothed and baleen, have complex social vocal behavior that is right-hemisphere lateralized with robust support from descending auditory pathways to the left cerebellum, toothed whales have also added a hypertrophic descending pathway from the left cortical hemisphere to the right cerebellum to manage the processing-intensive, highly time-constrained demands of echolocation. The question remains whether baleen whales do show right-lateralized vocal communication processing at the cortical level. A number of terrestrial species have been found to have left-lateralized vocalization mechanisms, but preliminary findings in hoofed mammals do not clearly support this pattern (Leliveld, 2019), leaving open the possibility that right-lateralized social vocal mechanisms are conserved in cetaceans from a terrestrial ancestor.

### Specific cerebellar targets

Existing data on the function of specific cerebellar lobules in other mammals can be tentatively extrapolated to support our conjectures regarding the strength and laterality of ascending and descending auditory pathways in cetaceans. Extensive physiological and anatomical evidence suggests that the posterior and lateral lobules of the cerebellum have evolved to support highly skilled behavior and domain-general cognitive abilities in humans and multiple distinct mammalian lineages (Schmahmann, 2005; Smaers et al., 2018). Meanwhile, medial and anterior cerebellar regions are hypothesized to subserve more traditionally-noted cerebellar functions like sensorimotor representation and limb coordination (Klein et al., 2016; Schmahmann, 2005). In humans, prefrontal (PFC) and parietal associative areas tend to coactivate with posterior and lateral areas, *particularly* Crus I and II, with Crus I being especially linked to PFC, executive function, and working memory tasks, and Crus II being especially linked to PFC *and* parietal association areas, social cognition, and theory of mind (Metoki et al., 2022; O’Reilly et al., 2009; Van Overwalle et al., 2020). Meanwhile, motor cortex tends to coactivate with anterior lobules plus lobule VIII (Metoki et al., 2022; O’Reilly et al., 2009), and the vermis appears to be most active in tasks involving socio-emotional processing (Klein et al., 2016). Interestingly, language processing appears to involve Crus I and II (Metoki et al., 2022), with Crus I being more linked to syntactic and Crus II to semantic processing (Nakatani et al., 2022), and generally, comparative evidence supports the idea that vocal learning and lateral cerebellar expansion are linked in mammalian lineages (Smaers et al., 2018). The apparent targeting of *left*-lobule Crus I and II by right IC-left cerebellar traces in the odontocetes seems to accord with the putative role of the *left* phonic lips in communication and social sound production in odontocetes (Ames et al., 2020; Madsen et al., 2013; Ridgway et al., 2015). This interpretation is further backed by the targeting of the vermis, also linked to socio-emotional processing (Klein et al., 2016), by the right IC-left cerebellar traces in two out of the three odontocetes.

Notably, compared to humans, Crus I and II are fairly diminutive in cetaceans (Cozzi et al., 2018), while lobule IX, VIIIa and b, VIIb, and VI are greatly expanded (Hanson et al, 2013). Existing mammalian findings help make sense of the pathways observed in these lobes in the current study. Suga and Horikawa (1986) found that lobules IX and VIIIa/VIIIb are active in bats during production of echolocation sounds, and additional studies in humans, felines, and other mammals suggest lobules VIIIa/VIIIb are related to audition, auditory motion processing, audiovisual integration, and sensorimotor representation (Baumann et al., 2014; Metoki et al., 2022). The targeting of *right* VIIIa in odontocetes, but bilateral VIIIa in the mysticete, is suggestive of this study’s core line of interpretation: namely, that left IC to right cerebellar pathways have proliferated to subserve echolocation and acousticomotor integration in odontocetes, while *non-social* acousticomotor tracts have remained bilateralized and are less developed in the mysticete. Furthermore, IC to lobule VI pathways were more right-dominant in the odontocetes, but were biased toward left cerebellum in the mysticete. Lobule VI has been linked to audition as well as somatosensory and motor representation in face, neck, head, tongue, and lips (Baumann et al., 2014; Cozzi et al., 2018; Hanson et al., 2013). In odontocetes, the left IC to right cerebellar tracts’ presence in lobule VI seems to align well with the proposed role of these tracts in echolocation, given that echolocation involves tight integrative looping of somatosensory information from the head with precise motor coordination of their musculature (Cook et al., 2024; Cozzi et al., 2018). It should be noted that Hanson et al. (2013) found that lobule VI was larger on the left than the right in tursiops. The size disparity might be driven by non-auditory processing related to sound production by left phonic structures (i.e., whistling) and patterns of non-IC connectivity from these lobules should be examined in toothed and baleen whales.

In sum, the observation of qualitatively pronounced tracts between right IC and left Crus I and II in odontocetes corroborates the hypothesized specialization of their right cortex and left cerebellum for social and communicative functions. Importantly, because Crus I and II are quite small in cetaceans (Cozzi et al., 2018; Hanson et al., 2013), tracts to these regions may not be as quantitatively robust as those to other lobules, and this helps make sense of the comparatively weaker tract strength observed in odontocete right IC to left cerebellar traces. Meanwhile, IC projections to lobules involved in sensorimotor representation, and especially in audition or head, neck, and face somatomotor activity, appear to be more pronounced between the left IC and right cerebellum in odontocetes, and this is the side that is *quantitatively* more robust in the odontocetes too. As mentioned previously, dolphin trigeminal and facial nerves heavily innervate the phonic lips and associated sound-production mechanisms. The qualitative-quantitative accordances here strongly support the hypothesis that right cortical to left cerebellum tracts support social and communicative functions across all cetaceans, but that in odontocetes, additional tracts have asymmetrically proliferated to connect the left cortex and right cerebellum to support the rapid acousticomotor integration demands of echolocation.

On the other hand, our cerebellar projection findings contain numerous remaining mysteries. Lobule IX is the largest in cetaceans but fairly small in humans, though it is unclear precisely what role it plays in cetaceans (Cozzi et al., 2018; Hanson et al., 2013). In rodents, this lobule seems to be involved in auditory *and* visual processing (Azizi et al., 1985; Burne & Woodward, 1983; Vogler et al., 2016); in bats, it is again linked to production of echolocation sounds (Suga & Horikawa, 1986); and in humans, it may play a role similar to Crus I/II in language and social cognition (Metoki et al., 2022), or may be linked to auditory processing as well (Mennink et al., 2020). Lobule IX’s expansion in cetacea therefore could be linked to an expansion of sensorimotor feedback and audio-visual integration capabilities for echolocation, but it may well serve as a substrate for expanded social and communicative behavior, too. The left-side prominence of IC to cerebellar tracts to this lobule in odontocetes, in light of the other findings at hand, tentatively suggests the latter. Finally, lobule VIIb, while fairly large in cetaceans (Hanson et al., 2013), is acutely understudied, rendering it difficult to postulate its functional significance in cetacea. The minimal literature that does exist suggests it might be involved in visuospatial working memory in humans (Brissenden et al., 2018). Some older evidence in cats indicates its potential role in audition and sensorimotor limb representation (Snide & Stowell, 1944). Though interpretations are limited by the sparseness of existing evidence, is possible that in cetaceans, this lobule may be involved in sensorimotor representation of the body stem and fins (Cozzi et al., 2018), and perhaps even in the continuous integration of spatial, motor, and proprioceptive information about the body and its motion within the environment.

### Contralateral ascending pathways

Our data pose a further mystery. While direct ipsilateral ascending tracts from IC to cortex were generally not lateralized, contralateral projections (from right IC to left cortex and from left IC to right cortex) *were* lateralized, with denser tracts going from left IC to right cortex. The exception to this was *D. delphis*, in which ipsilateral ascending tracts were more robust between right IC and right cortex than left IC and left cortex, and contralateral projections between IC and cortex were non-lateralized. The cerebella were necessarily excluded for all specimens in both ipsilateral and contralateral cortical traces used for the quantification of ascending auditory tract strength so that streamline counts did not include IC to cerebellar tracts. In follow-up traces of ipsilateral IC connectivity *without* cerebellar exclusion conditions in *D. delphis*, a less robust lateralization effect was observed, suggesting the seemingly robust right-lateralization initially discovered may in fact partially reflect a *relatively greater loss* of left-side IC tracts and a *relative lack of loss* of right-side connectivity in *D. delphis* upon the exclusion of contralateral and cerebellar connections (<u>S8 Table 1</u>). Similarly, follow-up traces in *L. acutus* revealed that removing cerebellar exclusion conditions in the contralateral IC-cortical traces attenuated the right-lateralization effect, suggesting that the right-lateralization effect may be partially mediated by a relative *lack of loss* of right-side IC connectivity due to the exclusion of the cerebella (<u>S8 Table 3</u>). These findings tentatively reinforce the notion that quantitatively robust cerebellar connectivity is more relevant to the left IC and cortex than the right in the odontocetes. It may be that some significant portion of auditory cortical projections to the left hemisphere in dolphins are reciprocally connected with the cerebellum. Such pronounced integration between left auditory cortex and right cerebellum, in light of the functional laterality hypothesis, further suggests this circuit’s role in the production and reception of echolocation sounds.

A separate interpretation of the finding of right-cortical bias for contralateral IC-cortical connectivity is that auditory information from both ears is preferentially sent to the hemisphere more responsible for processing communication, as opposed to echolocation. One possible explanation is that this represents segregation of echolocation sound processing to one hemisphere to prevent interference from communicative signals, as has been suggested in bats (Kanwal, 2012), or, more broadly, could reflect the specialization of different cortical regions for different types of neural processing. This may be useful given that dolphins employ high frequency pulses, produced by the left phonic lips, in communicative contexts, signals that have some spectral overlap with echolocation vocalizations (Ames et al., 2020; Herzing & Johnson, 2015). Further, communication and echolocation signals can be produced simultaneously (Ridgway et al., 2015). However, there may be other factors at play, such as asymmetry in sound receptivity of external hearing pathways on each side of the dolphin head (Ryabov, 2023). Lateralization of sound production and hearing in baleen whales is understudied, but if they are right-hemisphere lateralized for communicative sound production, it may be more efficient to get auditory information to this hemisphere as opposed to the left hemisphere for this species. There is a tendency toward right-side behavioral lateralization in cetaceans as well, with related perceptual adaptations (Karenina et al, 2016). Regardless of laterality, in terms of overall tract strength, interhemispheric connectivity in cetaceans is generally low (Wright et al., 2018), and our findings reflect this as well, as in all specimens, the strength of contralateral IC-cortical tracts were fairly weak compared to ipsilateral cortical projections.

### Limitations

In the current study, our findings were limited predominantly by sample size– it has been particularly difficult to acquire usable, preserved whole baleen whales for tractography, due to both regulatory concerns and the logistical demands of acquiring, fixing, and imaging such large brains. Further collaboration between scientists and veterinary and stranding response personnel may provide more opportunities for acquiring these valuable specimens and data.

As with all diffusion tractography studies, there are known limitations to the accuracy of the tensor fit and pathway construction (Schilling et al., 2019). However, our data were acquired with a specialized SSFP imaging sequence optimized for post-mortem tissue (McNab et al., 2009; Miller et al., 2012). This approach allows for high-resolution (sub mm isotropic), high signal-to-noise ratio imaging, with a large number of sampled angles (52 in our case), and makes use of the FSL crossing fibers model (Jbabdi et al., 2010), which, together, limits some of the concerns regarding anatomical precision. Further, the preprocessing involves specifically modeling and accounting for divergent T1 and T2 tissue values, which helps control for accumulating changes to tract integrity and diffusion parameters with increased time post-mortem. The primary tracts we assessed in the current study were very robust, even with heavy thresholding.

### Future Work

The current findings would be complemented by a parallel study with a sperm whale brain. Sperm whales have the smallest relative cerebellum size among the odontocetes (Ridgway & Hanson, 2014). In addition, they have only *one* pair of phonic lips, and these are situated on the right side of their head, where they are presumably employed for both social and echolocation sound production (Thornton et al., 2015). It is of great interest whether sperm whales also show right cerebellum lateralization of auditory processing.

The hypotheses put forth above also assume integration of auditory and vocal motor processing in cetaceans. However, there have been essentially no studies of vocal motor pathways in the cetacean brain. Their apparent capacity for vocal production learning and the relevance of vocal production to auditory processing in echolocation makes this a tempting target (Janik & Slater, 1997; Janik, 2014; Tyack & Sayigh, 1997). Although vocal motor regions of the cetacean *cortex* have not been identified, relevant brainstem nuclei have been mapped via prior histological work and could be used to seed tractography (Cozzi et al., 2018).

## Conclusion

In summary, here we have extended the diffusion tractography evidence on the odontocete auditory system, and have presented the first diffusion MRI data in a baleen species to wit, allowing a landmark comparison of auditory and acousticomotor tracts in echolocating versus non-echolocating whales. Each brain showed, as in Berns et al. (2015), a clear pathway from IC to temporal lobe via the thalamus, further challenging the account that primary auditory processing has been completely dorsally-shifted in odontocetes. The most striking difference between the species was in the density and lateralization of descending IC to cerebellum tracts, with notable hypertrophy in right-cerebellar tracts in the odontocetes, potentially corresponding to production of echolocation sounds by the right phonic lips and contralateral left-cortical processing. Meanwhile, density and lateralization of ascending auditory tracts did not differ significantly between the echolocating and non-echolocating species. We suggest that the specialization of auditory-cerebellar processing for echolocation in odontocetes may represent parallel evolution to that seen in echolocating bats. This heretofore underexplored neural component of the cetacean echolocation system should be more closely examined in future studies, including diffusion tractography, histology, behavioral, and noninvasive electrophysiological methods.

## Materials and Methods

### Specimens

In total, brains from four different cetaceans were used in this study, including three odontocetes and one mysticete. The first specimen was a post-mortem brain extracted from a pregnant adult female common dolphin (*Delphinus delphis*) that stranded dead in Buxton, North Carolina (Field #PTM135) in February 2001 (Berns et al., 2015). The second specimen was a post-mortem brain extracted from an adult female pantropical spotted dolphin (*Stenella attenuata*) that stranded dead at Camp Lejeune, North Carolina (WAM576). Both carcasses were in fresh condition (Smithsonian Condition Code 2 (Geraci & Lounsbury, 2005)) with no apparent damage, and the brains were extracted while still fresh. The total body length and body weight of the *D. delphis* and *S. attenuata* were 203 cm and 191 cm, and 83 kg and 57 kg, respectively. The brain of *D. delphis* had a fresh weight of 981 g, an anterior-posterior length of 132 mm, a bitemporal width of 155 mm, and a height of 96 mm. Up until the scanning of the *D. delphis* brain in Marino et al. (2002) and the scanning of both brains in Berns et al. (2015), both specimens were stored at Emory University, where they were kept in 10% neutral buffered formalin that was changed regularly. The images procured during the scanning session for Berns et al. (2015) are used in the current study. Immediately prior to the Berns et al. (2015) scanning session, the specimens were set in 2% agarose (Phenix Research Products Low EEO Molecular Biology Grade Agarose) doped with an insoluble mixture of 2 mM gadolinium (III) oxide (Acros Organics, Fisher Scientific).

The third specimen was a post-mortem brain extracted from an adult female Atlantic white sided dolphin (*Lagenorhyncus acutus;* specimen code CCSN03-164La) that was found dead in Cape Cod, Massachusetts in 2003. A necropsy was performed the day following the animal’s discovery, and the extracted brain was fixed in 10% phosphate-buffered formalin and stored at Woods Hole Oceanographic Institute until the scanning session described below. Histological analysis determined the animal had an extensive fungal infection with degeneration, necrosis, and inflammation. The total body length of the animal was 213.5 cm.

The fourth specimen was a post-mortem brain extracted from an adult sei whale (*Balaenoptera borealis*), sex and body length unknown. The whale was killed for its meat by Icelandic hunters sometime in the mid-1980s. Its brain was extracted and stored in formalin for decades, and was in remarkably good condition prior to its scanning in the 3 Tesla Siemens Trio MRI at University of California, Berkeley, as will be further described below.

### Imaging

As described in Berns et al. (2015), the brains of *D. delphis* and *S. attenuata* were scanned in a 3 T Siemens Trio using standard gradients (40 mT/m maximum) and a 32-channel head receive coil over the course of approximately 8 hours. Following Miller et al. (2012), DTI was carried out using a diffusion-weighted steady state free procession (DW-SSFP) sequence, as this acquisition method is preferable for maximizing the signal-to-noise ratio in diffusion-weighted images of post-mortem brains. For each brain, a set of DW-SSFP images were collected, weighted along 52 directions (FOV= 166 mm, voxel size= 1.3 mm isotropic, TR= 31 ms, TE= 24 ms, flip angle= 29°, bandwidth= 159 Hz pixel^−1,^ *q*= 255 cm^−1^, G_max_= 38 mT m^−1^, gradient duration= 15.76 ms (Berns et al. (2015)). Additionally, six images were collected to serve as a signal reference, and their acquisition followed the above parameters, except with *q*=10 cm^-1^ solely applied in one direction. Because the progressive relaxation of post-mortem brain tissue leads its T1 and T2 values to differ significantly from those observed in in vivo tissue, we calculated these values from a series of T1- and T2-weighted images to ensure proper modeling of the DW-SSFP signal. The T1-weighted images were acquired with a TIR sequence in which TR= 1000 ms, TE= 12 ms, and TI= 30, 120, and 900 ms, while T2-weighted images were acquired with a TSE sequence in which TR= 1000 ms, TE= 14, 29, and 43 ms. Structural image acquisition was performed using a balanced SSFP sequence (TR= 7.03 ms, TE= 3.52 ms, and flip angle= 37°), and balanced SSFP images were collected in pairs with the RF phase incrementing 0° and 180°, which were later averaged to minimize banding artifacts (Miller et al., 2012). The resulting structural images had a high resolution of 0.6 x 0.6 x 0.5 mm.

The *L. acutus* and *B. borealis* brains were scanned at the Brain Imaging Center of the University of California, Berkeley. The brains were again scanned in a 3 T Siemens Trio with standard gradients, a 32-channel head receive coil, and the DW-SSFP scanning protocol adapted from Miller et al. (2012). Because of its size, the *B. borealis* brain was held only in plastic Ziploc bags to prevent drying, and was inserted intact-hemisphere-first into the 32-channel coil. Meanwhile, the *L. acutus* brain was held in a plastic Ziploc bag filled with perfluoropolyether (PFPE) fluid, and was inserted rostral-first into the scanner with the cerebellum to the rear. Trying to hold the buoyant specimen fully under the fluid proved difficult, resulting in about 2/3 coverage. As a result, some fixative floating on the top of the PFPE is seen as a rim around the *q*= 20 images. As above, images were collected along 52 diffusion weighting directions, though this time a maximum of 8 diffusion weighting directions per block was established due to file size limitations on the acquisition computer. Accordingly, acquisition occurred in blocks of (3,8) = 11, (5,7) = 12, (3,5,6) = 14, and (7,8) = 15 directions. Extra *q*= 20 scans were included between every block to account for the long acquisition times. For *B. borealis*, FOV= 230 mm x 230 mm, voxel size= 1.1 mm^3^, TR= 34 ms, TE= 25 ms, flip angle= 44°, bandwidth= 90 Hz pixel^−1^, *q*= 270 cm^-1^, G_max_= 38 mT m^−1^, and gradient duration= 16.7 ms. Each individual DW-SSFP scan took 15 minutes and 14 seconds, for a total acquisition time of 16 hours. For *L. acutus*, FOV= 112 mm x 190 mm, voxel size= 1 mm^3^, TR= 34 ms, TE= 25ms, flip angle= 34°, bandwidth= 90 Hz pixel^−1^, *q*= 269 cm^-1^, G_max_= 38 mT m^−1^, and gradient duration= 16.7 ms. Each individual DW-SSFP scan took 9 minutes and 10 seconds, for a total acquisition time of 9 hours. Directed bore ventilation was used to maintain the specimens at a uniform temperature.

### Processing

Following image acquisition but prior to tractographical analysis, images were processed by adapting FSL tools to suit the DW-SSFP signal model. The processing method described in Berns et al. (2015) for the *D. delphis* and *S. attenuata* specimens was repeated here with the new specimens of *L. acutus* and *B. borealis*. To approximate the T1 and T2 values of the brains, we assessed the image intensities of several T1 and T2 images with different TR and TE values. The T1 and T2 values of all brains were estimated to be 350 and 50 ms, respectively, yielding an approximate *b*_eff_ value of 3500 s mm^-2^ in the diffusion-weighted scans (McNab et al., 2009). As before, all diffusion images were registered to a common q = 10 cm^−1^ reference image, and all reference images were averaged to create a mean reference image. The diffusion tensor model for describing anisotropic diffusion was fit to each voxel using a Metropolis Hastings algorithm that imposes a positivity constraint on tensor eigenvalues. This procedure produced estimates for the fractional anisotropy, mean diffusivity, and three eigenvectors of the diffusion tensors. Finally, a modified BedpostX model with DW-SSFP signal equations was applied with default settings, save for the specification of 2 crossing fibers per voxel (Buxton, 1993; McNab et al., 2009; Miller et al., 2012).

### ROI selection and tractography

Following Berns et al. (2015), the inferior colliculi (IC) were selected as the key region of interest (ROI) for the auditory pathway in all specimens. The IC were selected as the start point because nearly all ascending auditory information passes through them (Cozzi et al., 2018), and also because they are large, prominent, and easy to identify. Additionally, diffusion-weighted imaging has been used to plausibly trace IC tracts in humans (Sitek et al., 2022). For reference, in the following, all referenced sagittal planes span between the lateral-most points of each hemisphere; all coronal planes span from the rostral-most point of the cerebral cortex to the caudal-most point of the cerebral cortex; and all axial planes span from the dorsal-most vertex of the cortex to the ventral-most point of the brainstem. In other words, analyzed brains are presented in orientations akin to those employed for human brains, rather than the more naturalistic 90-degree-rotated orientation that reflects the actual positioning of the brain in the skull of cetaceans in vivo. In all specimens, the IC were identified as round protrusions on the caudal surface of the midbrain, apparent in the sagittal view dorso-rostral to the cerebellum and ventro-caudal to the superior colliculi and thalamus; apparent in the coronal view dorsal to the lateral lemniscus and ventral to the thalamus; and apparent in the axial view rostral to the ventral extremity of the cortical hemispheres, and caudal to the dorsal extremity of the pons. For the reasons described previously, the cerebellar hemispheres were deemed instrumental for the study of descending acousticomotor pathways, and were designated as the second ROI. The cerebellar hemispheres were also easily identified by their large size and unmistakable gross anatomical structure, protruding from the caudal surface of the midbrain and brainstem, just ventro-caudal to the inferior colliculi.

ROIs used to seed tractography were created as masks over high-resolution T2-weighted structural images in FSLeyes. In the present study, all voxels in each inferior colliculus and cerebellar hemisphere (left and right) were selected in accordance with the above-listed anatomical guidelines (Fig 9). Additionally, single-voxel-wide whole-plane masks were drawn at the medial-most sagittal plane between left and right hemispheres in order to examine and quantify tracts that were strictly contralateral and strictly ipsilateral. A single-voxel-height mask of an axial slice of the pons and upper brainstem was also created to allow specific analysis of tracts that ascended beyond this axial plane to the diencephalon and telencephalon, and to exclude tracts that did not pass to a level more dorsal than the metencephalon and myelencephalon. The masks were then loaded with the BedpostX file into the FSL tract-tracing tool, ProbtrackX, in order to perform the probabilistic tracing. Tracking was performed with the default settings of 5000 samples per voxel, 0.2 curvature threshold, and 0.5mm step length. The sole change made to ProbtrackX default settings was the enabling of the distance correction setting, given the large size of the brains being analyzed. To allow specific and isolated analysis of the potential transmission of auditory information via each IC to ipsilateral and/or contralateral cortices and cerebella, we conducted a systematic battery of traces in which traversal of at least one voxel in a cerebellar hemisphere and/or the contralateral side of the brain was either necessary for inclusion of a tract (i.e., mask files were added as ProbtrackX waypoints) or grounds for exclusion of a tract (i.e., mask files were added as ProbtrackX exclusion masks). The full battery of traces, performed with every possible permutation of inclusion/exclusion criteria, can be viewed in S8 Tables.

**Fig 9:**
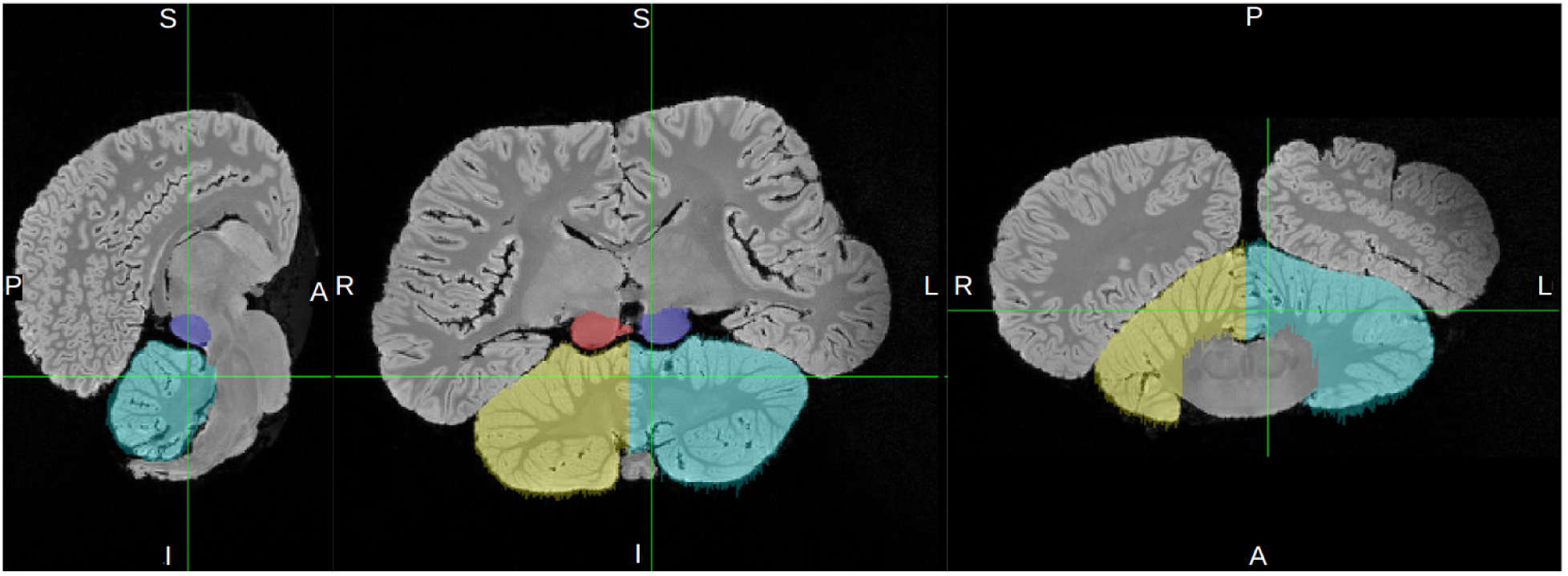
Inferior Colliculi and Cerebellum ROIs. Sagittal, coronal, and axial cross-sections showing the masks made for the cerebella (turquoise and yellow for left and right, respectively) and inferior colliculi (blue and red for left and right, respectively) in *B. borealis*. Refer to S3 Figures to view these masked ROIs in the other specimens.

Following tractography, we quantified the output using *waytotal* values: the count of streamlines that satisfy the tractography algorithm by originating in the seed ROI and satisfying waypoint and/or exclusion criteria (Thaploo et al., 2022). Additionally, volumetrics for each animal and each ROI were calculated by dividing the volume of each masked ROI by its specimen’s whole-brain volume. Notably, the volumetric values for each whole brain and region may not precisely reflect the true volumes in live animals, as formalin fixation can lead to marked tissue shrinkage (Ridgway & Hanson, 2014)--though Cook et al. (2018) and Montie et al. (2010) present evidence to the contrary. Additionally, though differences in fixation times may cause variations in degrees of shrinkage *between* specimens, to wit, there is no substantial evidence suggesting disproportionate shrinkage occurs between different regions, suggesting that calculating the *relative* volumes of regions within formalin-fixed brains remains valid. For each diffusion tractography output, i.e.,“trace,” these whole-brain volume measurements were used to calculate tract *strength* by dividing the trace’s waytotal by the volume of the whole respective brain. Tract *lateralization* was calculated first by simply dividing the larger waytotal by the smaller one (ex. right IC trace waytotal divided by left IC trace waytotal) to provide the *factor* by which each trace was left- or right-lateralized. Additionally, the lateralization index (LI) for each trace was calculated by applying the LI formula used in Vernooji et al. (2007) and Wright et al. (2018), (left - right) / (left + right), to the left and right waytotal values for each trace. Finally, waytotal values also provided image thresholding guidelines for qualitative analysis, as discussed below.

### Thresholding

All tract images in this paper are presented at one of two thresholds: a more conservative one optimized for the FSLeyes orthographic view and a more liberal one optimized for the FSLeyes three-dimensional (3D) view. For the orthographic images, traces were thresholded with a minimum value set to 1% of their waytotal and a maximum value set to 30% of their waytotal. Because the 3D view in FSLeyes tends to render tractograms more weakly than in the orthographic view, in the interest of conferring useful and detailed visual information about the sites of tract projection, we selected a more liberal threshold for tracts in the 3D view. For these images, traces were thresholded with a minimum value set to 0.10% of their waytotal, and a maximum value set to 5% of their waytotal. See the highlighted rows in S8 Tables for the raw waytotal values that were used to calculate these thresholds.

## Supporting information

3D rotation of common dolphin IC-contralateral cortex tracts

3D rotation of pantropical spotted dolphin IC-contralateral cortex tracts

3D rotation of atlantic white sided dolphin IC-contralateral cortex tracts

3D rotation of sei whale IC-contralateral cortex tracts

3D rotation of pantropical spotted dolphin IC-cerebellum tracts

3D rotation of common dolphin IC-cerebellum tracts

3D rotation of atlantic whited dolphin IC-cerebellum tracts

3D rotation of sei whale IC-cerebellum tracts

3D rotation of pantropical spotted dolphin ascending auditory tracts

3D rotation of common dolphin ascending auditory tracts

3D rotation of atlantic white sided dolphin ascending auditory tracts

3D rotation of sei whale ascending auditory tracts

## Acknowledgements

Thank you to Lori Marino, Ann Pabst, and William McLellan for specimens, and thank you to Karla Miller’s colleagues at Oxford University for supporting our use of their SSFP imaging protocols.

## Ethics Declaration

The authors affirm that this work involved no financial or commercial incentives that could create potential conflicts of interest. Additionally, we affirm that no animals were harmed in this research, as all tissue was obtained from opportunistically-collected post-mortem specimens.

## Supporting Information

### S1 Text. Detailed cerebellar and subcortical projection sites in IC-cerebellar traces

In left IC to right cerebellar traces in *D. delphis*, the strongest cerebellar projections were at the rostral extreme of the cerebellum, particularly in its connections with brainstem nuclei and the pons (S6 Figure A1-2). The most rostral and dorsal of these tracts passed from a rostro-dorsal region of left cortex to the right cerebellum via the substantia nigra and crus cerebri, just dorsal to the pons (S6 Figure A1-2). From here, one branch of the trace ran ventrally to join with a larger clump of tracts traversing the pons, while another remained dorso-rostral, connecting with a dorsal-rostral extreme of the cerebellum, likely a rostral extreme of lobule IX (S6 Figure A1-2). Toward its ventral extreme, the robust pontine fiber bundle branched off into three directions. One branch extended contralaterally and caudally to the left-side trapezoid body, cerebellar peduncle, and lateral lemniscus (S6 Figure A2); a second branch remained rostral but moved laterally and ventrally via a cerebellar peduncle to putative lobule IX (S7 Figure A1); a third branch forked off from the second after passing through the cerebellar peduncle, projecting more caudally to putative lobule VIIb, a projection mirrored in the right IC-left cerebellar trace (S7 Figures A1-2). Near the rostral-caudal fork between the latter two paths, a small projection to putative lobule VIIIa can be seen (S7 Figures A1-2). Lobule X, also fairly small in cetacea, also appears to receive bilateral projections from both left and right IC in both cerebella (S7 Figure A1).

In contrast to the left IC to right cerebellar traces, the right IC to left cerebellar trace in *D. delphis* did not include a clear path from cortex to cerebellum through the rostral midbrain. There were robust tracts in the ventral pons, and similarly to the left IC to right cerebellar traces, these forked into multiple pathways, including one that passed to right-side auditory structures like the trapezoid body and lateral lemniscus, and another to a rostral portion of left-cerebellar lobule IX via the peduncles (S7 Figures A1-2). The caudal-reaching fork in this trace had a projection that reached a more caudal-lateral extreme than was observed in the left IC-right cerebellar trace, likely near Crus I and II (S7 Figure A2). Additionally, the right IC to left cerebellar trace also exhibited a more robust path to ventral lobule IX than any observed in left IC to right cerebellar traces (S7 Figure A2). Rostral to these sites, there was a rostral and medial projection to putative lobule VIIb, similar to the projection to this lobule observed in the left IC to right cerebellar trace (S7 Figures A1-2). Notably, the right IC to left cerebellar tracts were more bilaterally robust in the brainstem auditory regions like the lateral lemniscus and trapezoid body, and additionally made it further down the brainstem, than left IC to right cerebellar tracts (S7 Figures A1-2).

Similarly to *D. delphis,* the left IC-right cerebellar traces in *S. attenuata* exhibited multiple distinct pathways at the rostral extreme of the pons, with three apparently transverse bundles at its dorsal extreme, its ventral extreme, and a more medial dorsal level, possibly representing pathways through the superior, inferior, and medial cerebellar peduncles, respectively (S7 Figure B1). In contrast to *D. delphis*, all three of these, including the dorsal-most of the bundles, was also observed in the right IC to left cerebellar traces, and direct cortical connections through rostral diencephalon and mesencephalon were not observed in either (S7 Figures B1-2), though more caudal ones from IC to putative MGN were (S6 Figures B1-2). As in the other dolphins, the majority of the pontine projections for both traces appeared to target putative lobules VIIIb and IX (S7 Figures B1-2). Also, as observed in both other dolphin specimens, the right IC-left cerebellar traces in *S. attenuata* contained a far-lateral caudal projection, likely to putative left Crus I and II, not mirrored in the right cerebellum and left IC-right cerebellar traces (S7 Figures B1-2). In contrast to the other dolphins, both traces displayed robust pathways throughout the vermis (Fig 7b). Furthermore, compared to other specimens, the tracts in the brainstem regions between pons and cerebellum in *S. attenuata* tended to be more *bilaterally* robust for both left IC to right cerebellar and right IC to left cerebellar traces (S6 Figures B1-2). Finally, more clearly than in any other specimens, both IC to cerebellar traces in *S. attenuata* displayed prominent dorsal-ventral projections through the mylencephalon, possibly traversing the inferior cerebellar peduncle, that proceeded ventrally and laterally from the floor of ventricle IV on the caudal brainstem, potentially contacting the facial colliculi situated here, to the caudal border of the inferior olives and eventually the rostral-most portion of the spinal cord (Fig 7b). Interestingly, the left IC to right cerebellum trace, but *not* the right IC to left cerebellum trace, produced a second, similarly continuous and robust projection between the floor of ventricle IV and the caudal edge of the inferior olives, medial and slightly rostral to the bilateral parallel projections described above, closer to the facial motor nucleus (Fig 7b).

In *L. acutus*, both IC to cerebellar traces had their most concentrated paths in the transverse fibers of the pons most rostrally, and in all three of the cerebellar peduncles most caudally (S7 Figures C1-2). In left IC to right cerebellar traces, the most prominent of these peduncular paths appeared to target the juncture of putative lobules VIIIa and VIIIb, a pattern mirrored in the right IC to left cerebellar traces of *L acutus* (S7 Figures C1-2). We also observed a unique feature of left IC to right cerebellar traces that was *not* observed in their contralateral counterparts: a second projection more medial, caudal, and dorsal than the first, fairly close to vermis, likely targeting putative lobule VI (S7 Figure C1), a projection that was only observed here and in the left cerebellum of *B. borealis* (S6 Figure D1).

The right IC to left cerebellar traces in *L. acutus* included a unique feature also observed in those of *D. delphis*: a more caudal-lateral projection than any observed in the left IC to right cerebellar traces, likely to putative Crus I or Crus II (S7 Figure C2). Further rostral to this, the most pronounced projection to left cerebellum was found in the putative conjunction of lobules VIIIa and VIIIb, connected to the auditory brainstem regions caudal to the ventral pons via the peduncles as was observed in the left IC to right cerebellar traces of *L. acutus* (S7 Figures C1-2). Notably, right IC to left cerebellar tracts were more bilaterally robust in these brainstem regions caudal to the ventral pons than left IC to right cerebellum tracts, contacting the right cerebellar peduncles in addition to the left ones (S7 Figure C2). Additionally, the right IC to left cerebellar traces contained a secondary projection not mirrored in the left IC to right cerebellar traces. This projection was almost as robust as the one to lobules VIIIa and VIIIb, and was found more rostrally in the coronal plane as the most robust transverse pontine fiber crossover, likely targeting lobule IX (S7 Figure C2). Finally, right IC to left cerebellar traces in *L. acutus* included a rather robust set of tracts through the vermis, particularly vermal lobules IV, VI, and VIIat, that were not observed in the left IC to right cerebellum traces of this specimen (Fig 7c).

In *B. borealis*, right IC to left cerebellar traces exhibited four main projections: one to a far-rostral point in putative lobule IX (S6 Figure D1); a second that forked from the far-rostral projection to a lateral extreme that was slightly more caudal but still rostral of the midpoint of the rostral-caudal axis, putatively identified as a lobule VIIb projection (S6 Figure D1); a third that projected caudally, ventrally, and laterally relative to the first two towards the border of lobules VIIb and VIIIa (S6 Figures D1-2); and finally, a fourth far-caudal and far-dorsal projection, directly lateral to the vermis, that forked into two projections that approached putative lobule VI and crus I (S7 Figure D2). Finally, right IC to left cerebellum traces contained more paths through the pons than left IC to right cerebellum traces (S7 Figure D2).

By contrast, left IC to right cerebellar traces in *B. borealis* featured only two pronounced projections, and these mirrored the first and third above-listed ones observed in right IC to left cerebellar traces: one to a lateral, far-rostral, and mid-ventral location, putatively identified as lobule IX (S6 Figures D3-4), and the other to a lateral far-caudal and ventral site around the junctures of lobules VIIb and VIIIa (S7 Figure D1, S6 Figure D2). The left IC to right cerebellum trace also contained a minor projection to *left* cerebellum’s putative lobule VIIb, directly overlapping with the right IC to left cerebellar trace’s projection to this site (S6 Figure D1).

### S2 Text. Detailed cortical projections in IC-cerebellar traces

In *D. delphis*, the left IC to right cerebellar trace contained a visibly robust cortical projection to a medial-rostro-dorsal extreme in the left and right hemispheres alike, potentially in or near the anterior cingulate cortex (S6 Figures A1-4). These medial-rosto-dorsal cortical projections from the left IC to right cerebellum trace reached both hemispheres, crossing over visibly via the corpus callosum, and were accompanied by an additional rostro-dorsal projection site that was more lateral, potentially in or near putative motor and premotor regions, and was much more pronounced in the left cortex (S6 Figure A2). Notably, the cortical projections of the quantitatively weaker right IC to left cerebellum traces were primarily located in the temporal lobe of the left cortical hemisphere, with some sparse tracts appearing alongside the aforementioned left IC to right cerebellum projections in the rostro-dorsal regions of the left cortex, and with very few tracts visible in the right cortex (S6 Figures A1-4). On the whole, in *D. delphis*, these traces differed from ascending auditory ones in two key ways: first, temporal lobe projections were considerably weaker in left cortex and absent in right cortex, and those observed in left cortex were primarily supplied by the contralateral right IC; second, the more rostro-dorsal projections in left cortex were primarily supplied by the ipsilateral left IC and contralateral right cerebellum, whereas in the ascending auditory traces, the contralateral right IC appeared to innervate this rostro-dorsal region of left cortex more prominently (see S4 Figures A1-4 versus S6 Figures A1-4).

In contrast to *D. delphis*, in *L. acutus*, the cortical projections observed in the IC to cerebellar traces were most prominent in the right rather than left hemisphere; specifically, right IC to left cerebellar tracts were more likely to be observed in the right cortical hemisphere than left IC to right cerebellar tracts in the left cortical hemisphere (S6 Figures C1-4). Interestingly, right IC to left cerebellar traces provided the vast majority of cortical projections in the left hemisphere; left IC to right cerebellar traces showed very few left-hemispheric cortical projection sites (S6 Figures C1-4). Both right IC to left cerebellar and left IC to right cerebellar traces projected to similar cortical areas as those implicated in ascending auditory traces in *L. acutus,* though the overlap was most *minimal* in the caudal-most projection sites reached by ascending auditory traces (compare S4 Figure C2 with S6 Figure C2). IC to cerebellar traces in *L. acutus* differed from ascending auditory traces most notably in the extent of their dorsal-rostral projections; in the right hemisphere and left hemispheres, respectively, left IC to right cerebellar and right IC to left cerebellar traces were observed reaching a dorso-rostral extreme that was not reached by plain ascending auditory traces (most apparent in S6 Figure C4).

Out of all of the dolphins, *S. attenuata* exhibited the weakest cortical projections in IC to cerebellar traces. As in the case of *L. acutus*, cortical projections for these traces overlapped the *least* with those in ascending auditory traces in the caudal and ventral cortical extremes, and were more likely to appear in dorsal extremes than their ascending auditory counterparts (compare S4 Figure B2 with S6 Figure B2). On the other hand, in *S. attenuata*, a more unique dorso-caudal left-cortical projection site was observed in both right IC to left cerebellar traces and left IC to right cerebellar ones (S6 Figure B2). Mirroring the case of *D. delphis*, both right IC to left cerebellar traces and left IC to right cerebellar traces were more likely to innervate the left cortex than the right, though in *S. attenuata* this tendency was less pronounced, as *some* right IC-left cerebellar projections to right cortex appeared at the more liberal threshold (S6 Figure B2).

As stated in the main text, cortical projections from IC to cerebellar traces were incredibly sparse in *B. borealis* (S6 Figures D3-4). Some weak tracts from left IC to right cerebellar traces were observed in the left thalamus at the more conservative threshold (S6 Figure D1), and the more liberal threshold revealed minor right IC-left cerebellar tracts that reached right dorso-rostral cortex (S6 Figure D2). No other cortical projections were robust enough to remain visible after thresholds were applied.

### S3 Figures. Masked regions of interest in FSLeyes

**Figure S3A.**
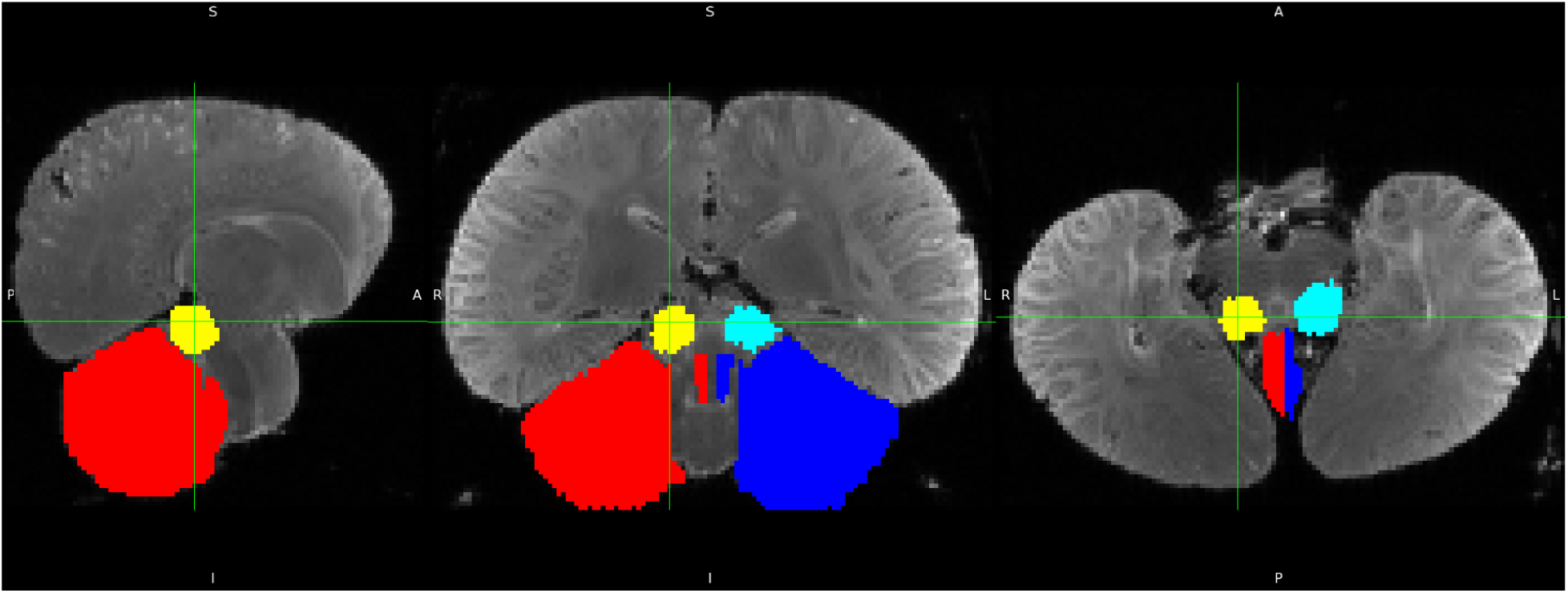
*D. delphis*. Red= right, blue=left for cerebella, yellow=right and turquoise=left for inferior colliculi.

**Figure S3B.**
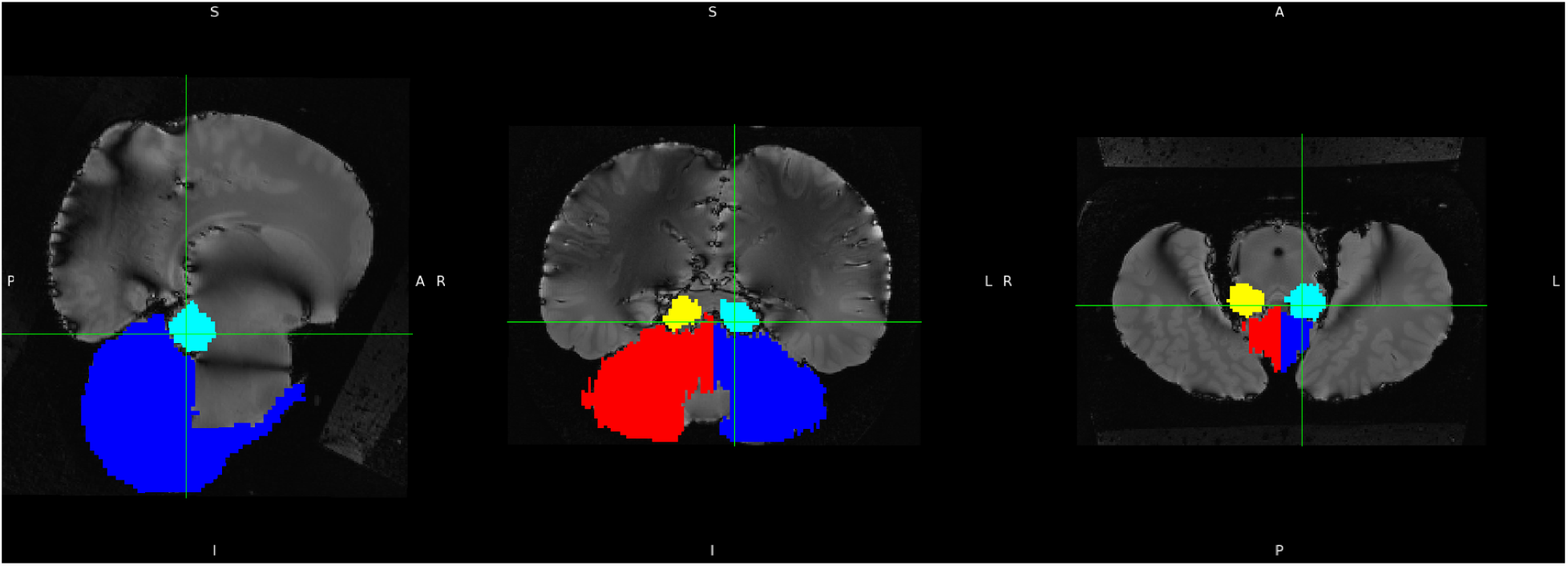
*S. Attenuata*. Red= right, blue=left for cerebella, yellow=right and turquoise=left for inferior colliculi.

**Figure S3C.**
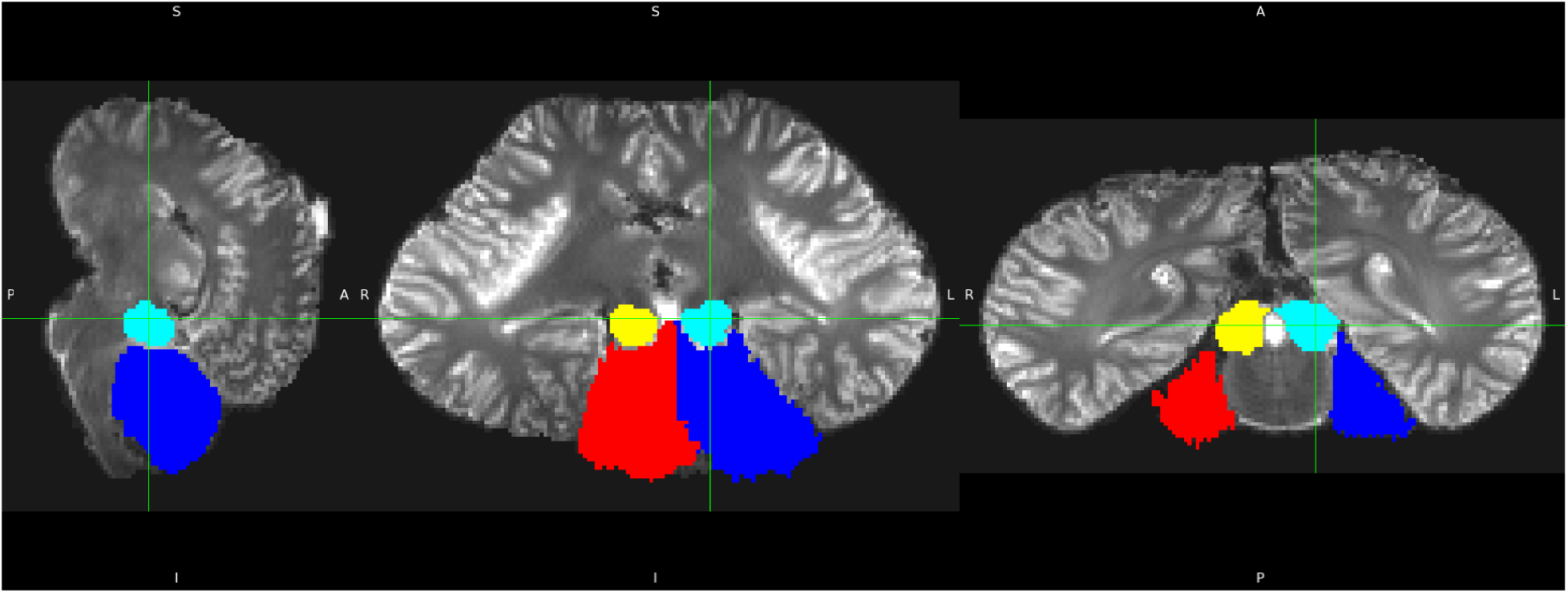
*L. acutus*. Red= right, blue=left for cerebella, yellow=right and turquoise=left for inferior colliculi.

**Figure S3D.**
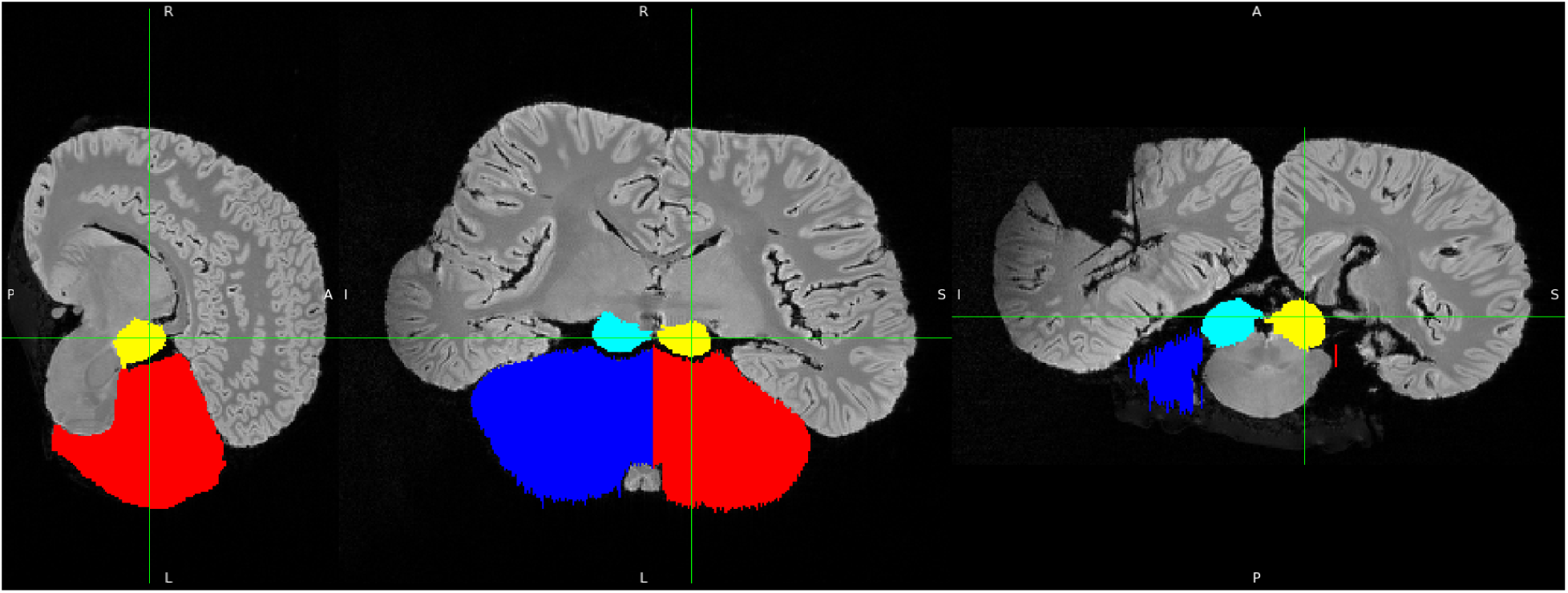
*B. borealis.* Red= right, blue=left for cerebella, yellow=right and turquoise=left for inferior colliculi.

### S4 Figures. Ascending auditory tractograms

**Figure S4A1:**
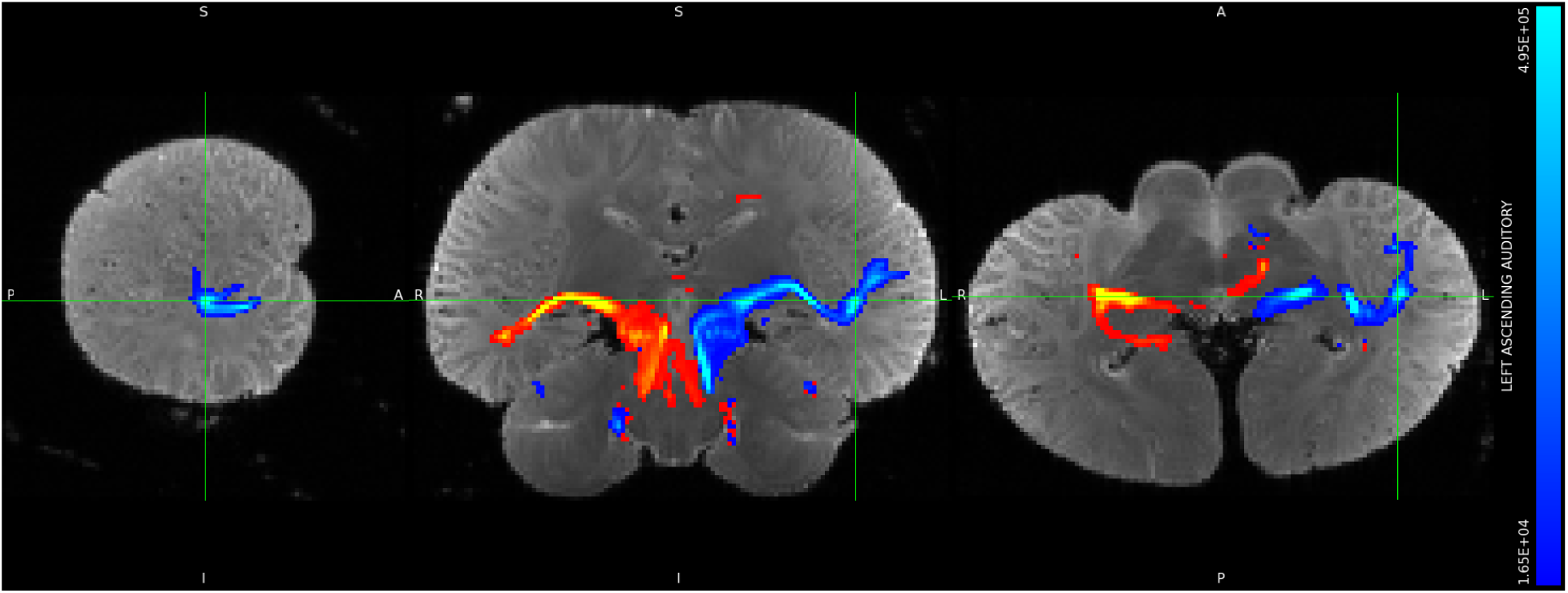
*D. delphis,* left IC tracts shown in blue, right IC tracts shown in red, minimum threshold set to 1% and maximum threshold set to 30% of waytotals. Orthographic view.

**Figure S4A2:**
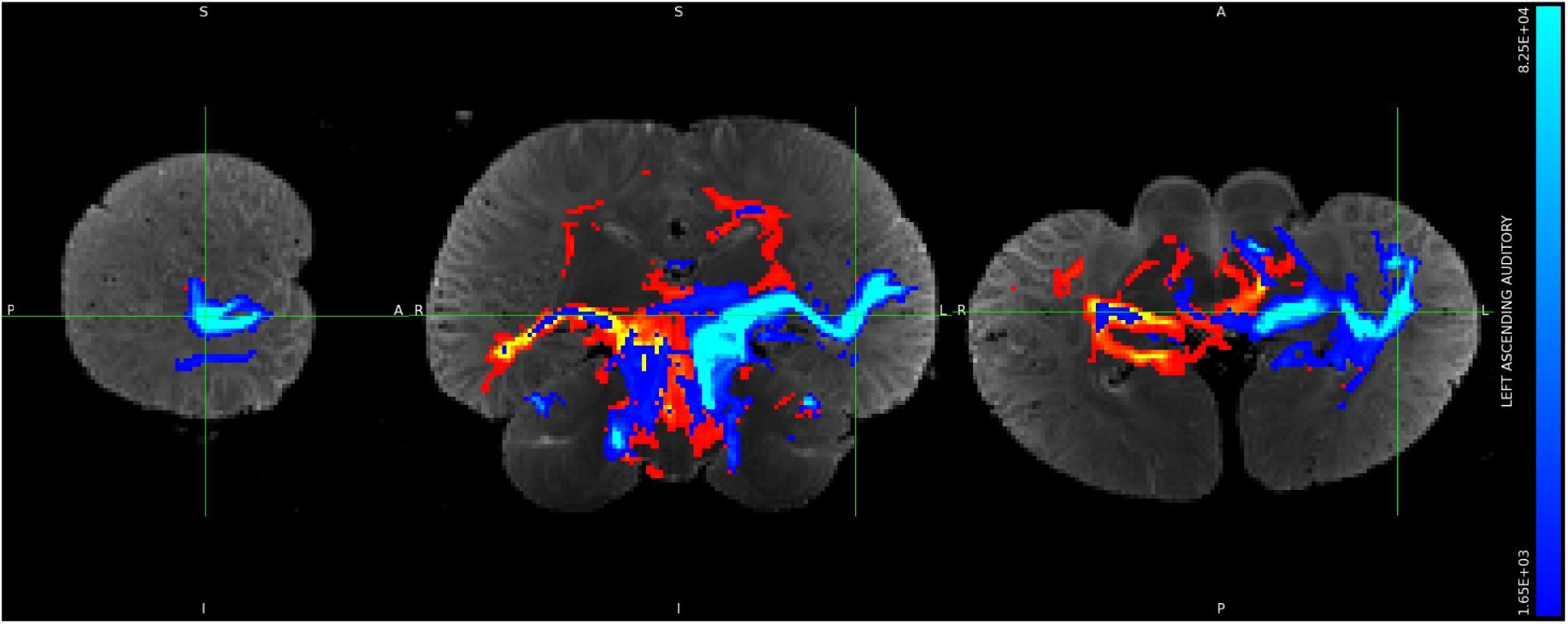
*D. delphis,* left IC tracts shown in blue, right IC tracts shown in red, set to a more liberal threshold of minimum 0.1% and maximum 5% of waytotals. Orthographic view.

**Figure S4A3:**
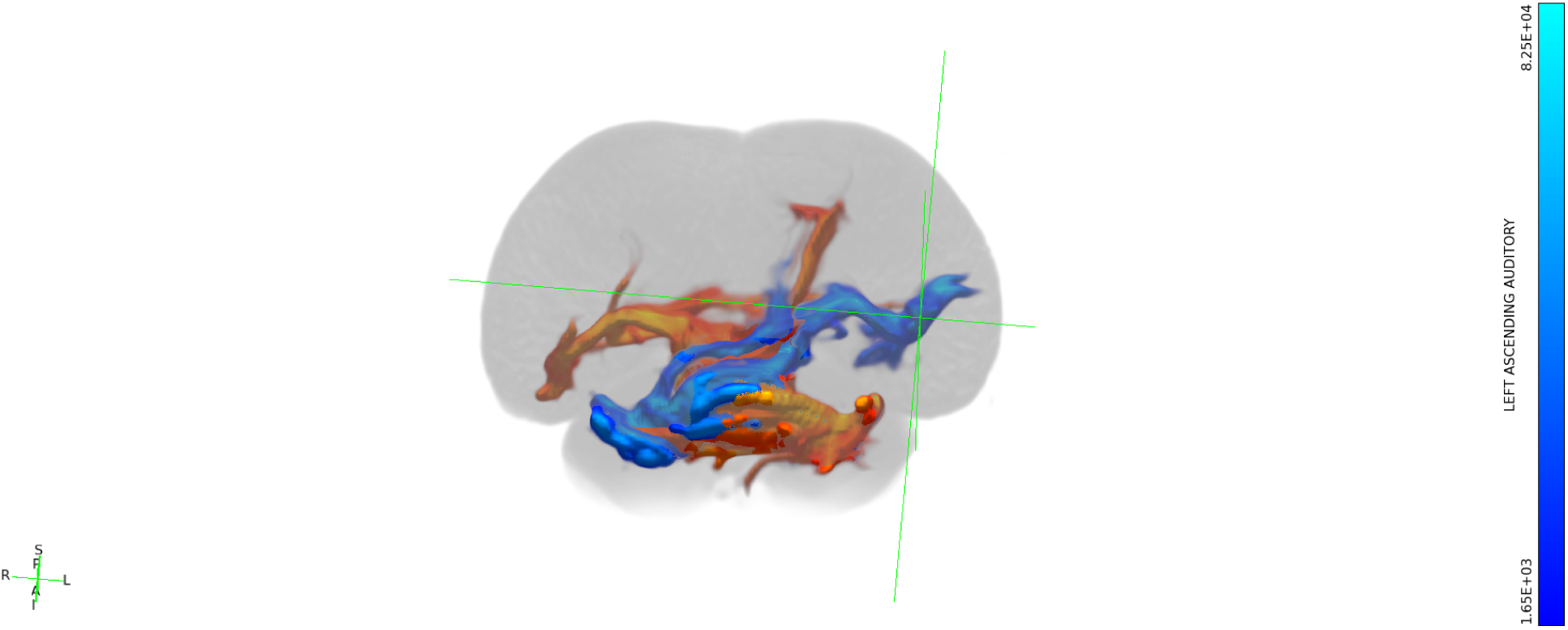
*D. delphis,* left IC tracts shown in blue, right IC tracts shown in red, set to a more liberal threshold of minimum 0.1% and maximum 5% of waytotals. Still 3-dimensional view.

**Figure S4A4:**
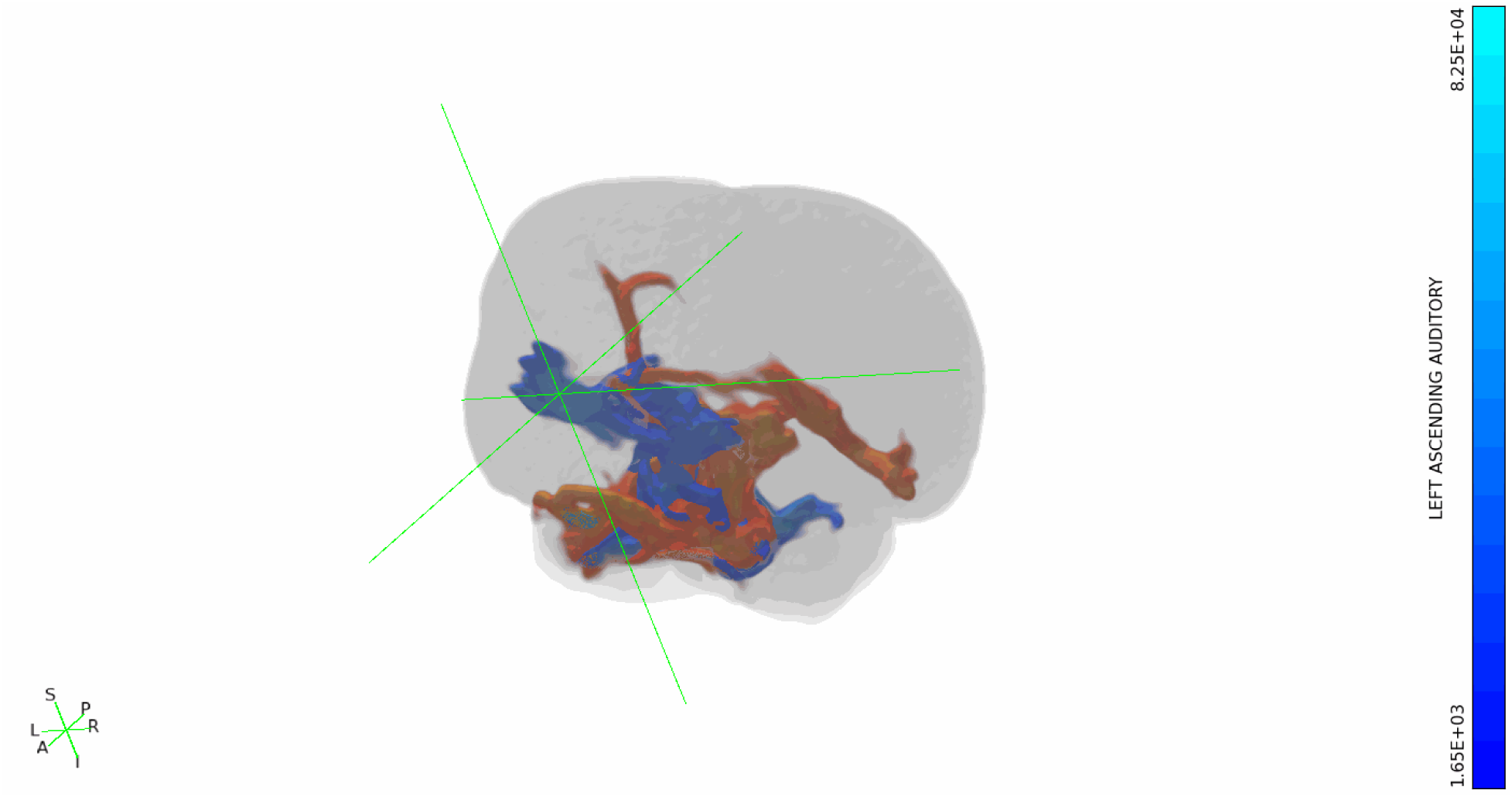
*D. delphis,* left IC tracts shown in blue, right IC tracts shown in red, set to a more liberal threshold of minimum 0.1% and maximum 5% of waytotals. Rotating 3-dimensional view.

**Figure S4B1:**
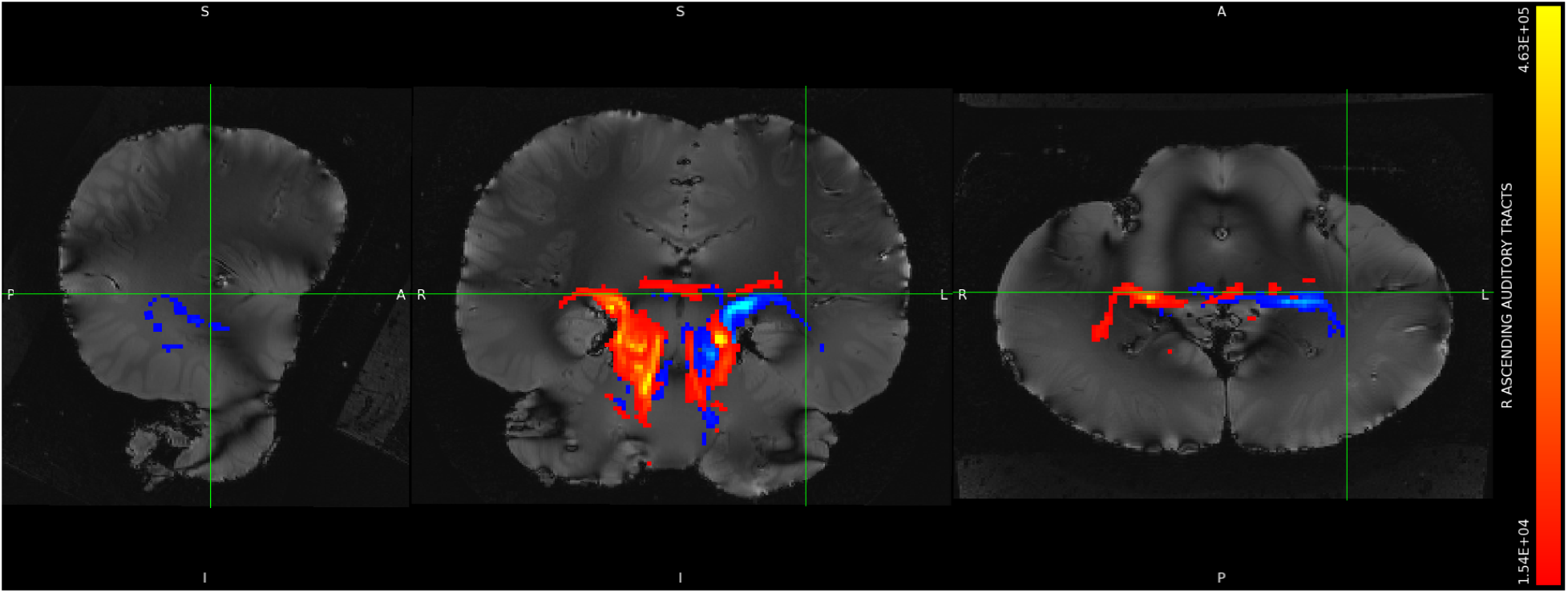
*S. attenuata,* left IC tracts shown in blue, right IC tracts shown in red, minimum threshold set to 1% and maximum threshold set to 30% of waytotals. Orthographic view.

**Figure S4B2:**
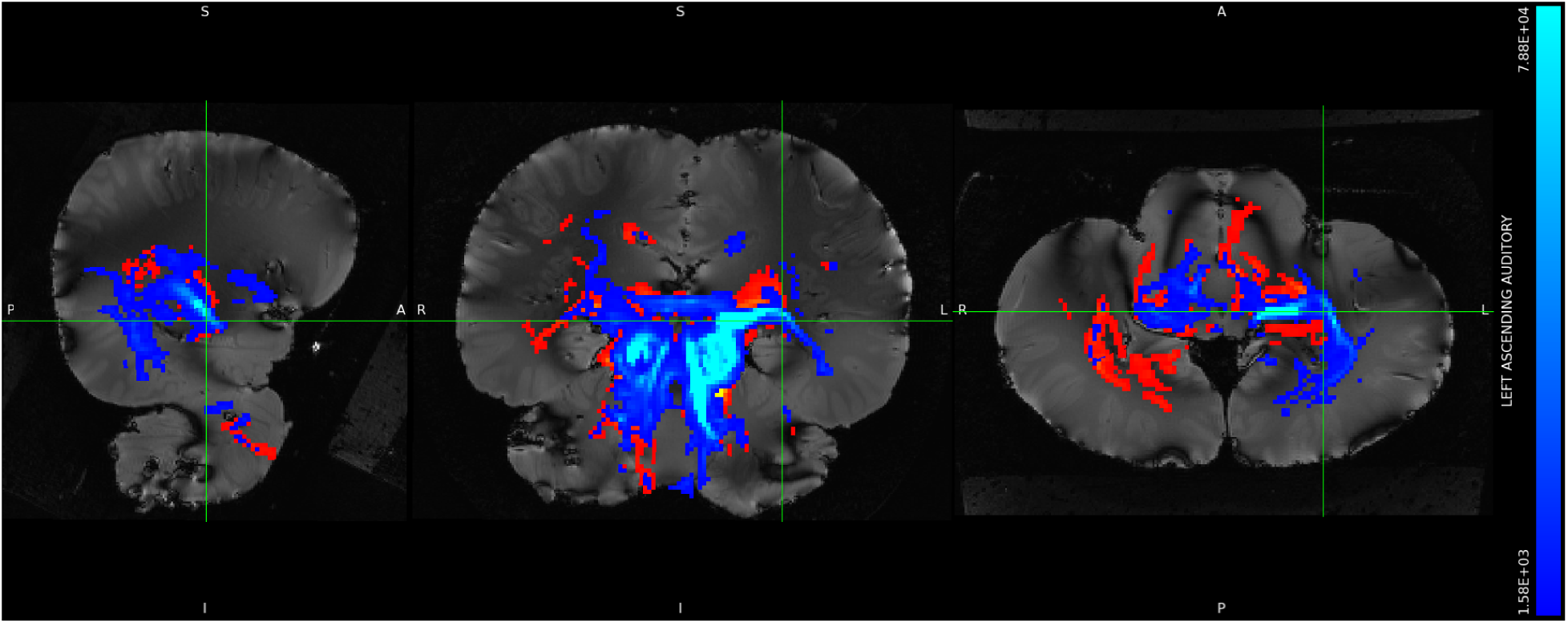
*S. attenuata,* left IC tracts shown in blue, right IC tracts shown in red, set to a more liberal threshold of minimum 0.1% and maximum 5% of waytotals. Orthographic view.

**Figure S4B3:**
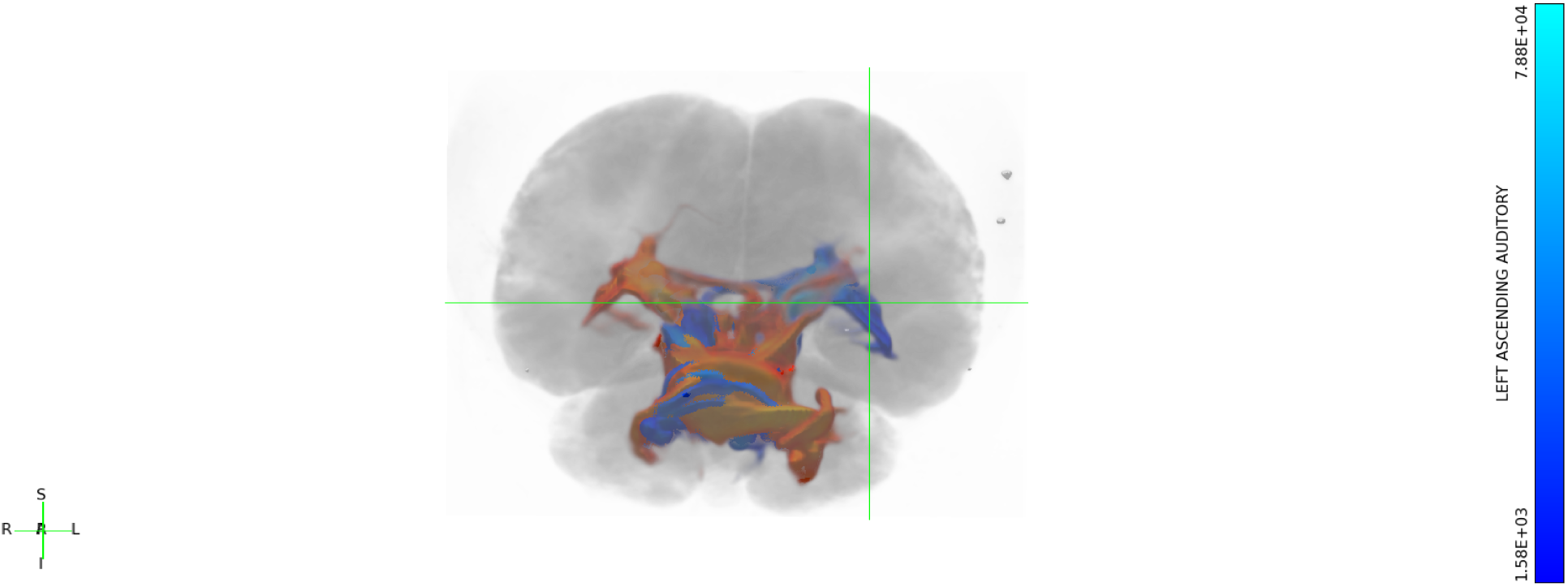
*S. attenuata,* left IC tracts shown in blue, right IC tracts shown in red, set to a more liberal threshold of minimum 0.1% and maximum 5% of waytotals. Still 3-dimensional view.

**Figure S4B4:**
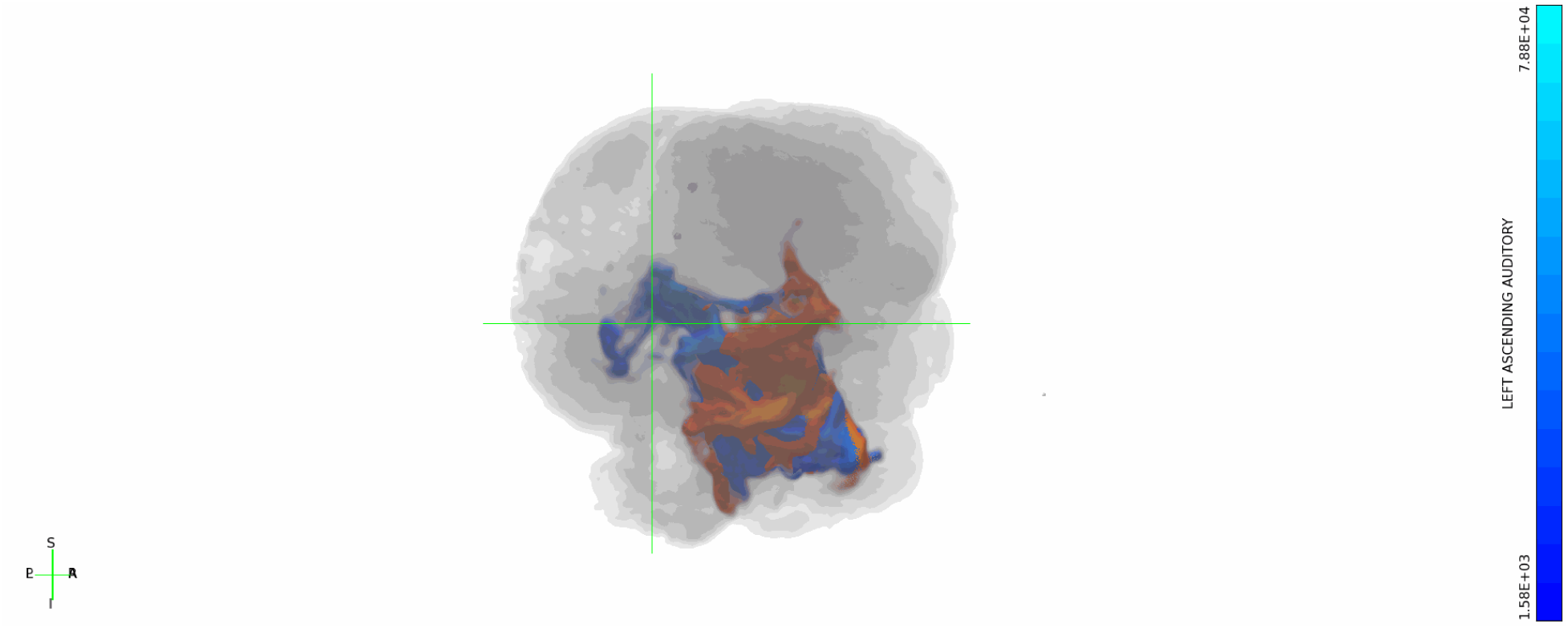
*S. attenuata,* left IC tracts shown in blue, right IC tracts shown in red, set to a more liberal threshold of minimum 0.1% and maximum 5% of waytotals. Rotating 3-dimensional view.

**Figure S4C1:**
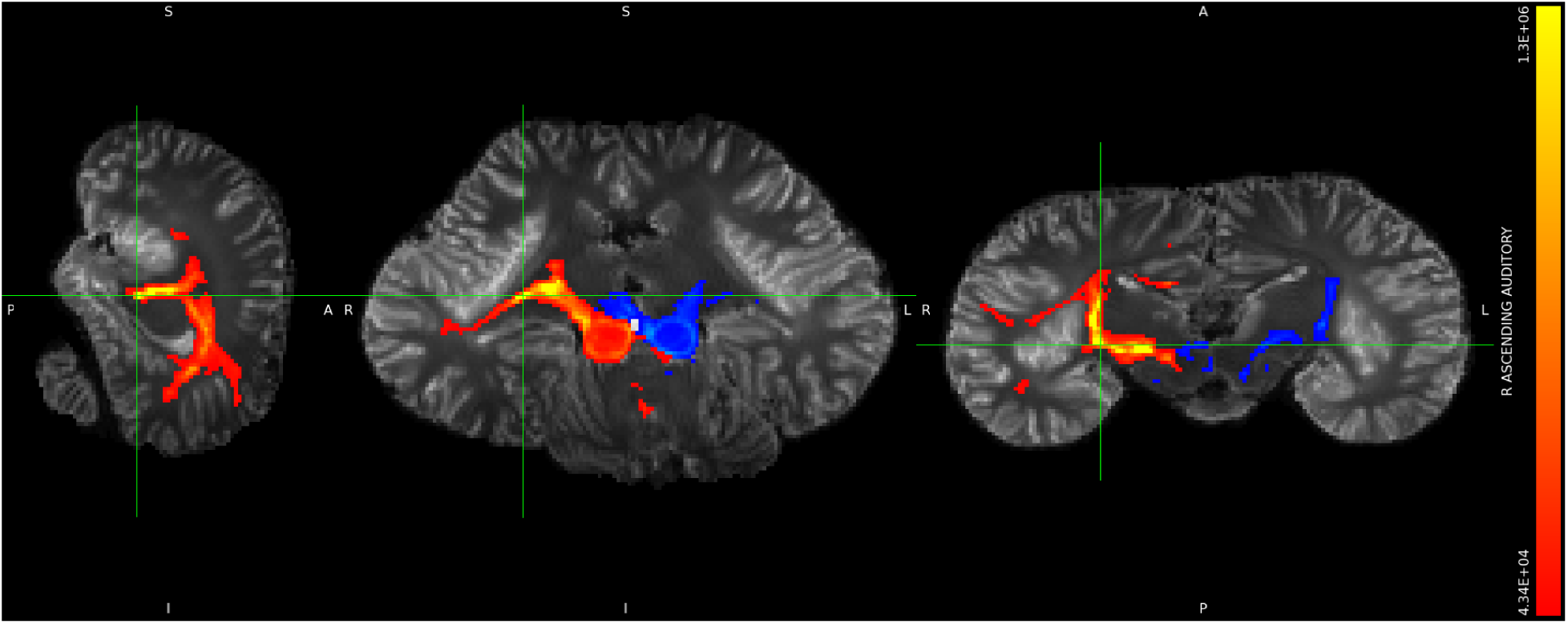
*L. acutus,* left IC tracts shown in blue, right IC tracts shown in red, minimum threshold set to 1% and maximum threshold set to 30% of waytotals. Orthographic view.

**Figure S4C2:**
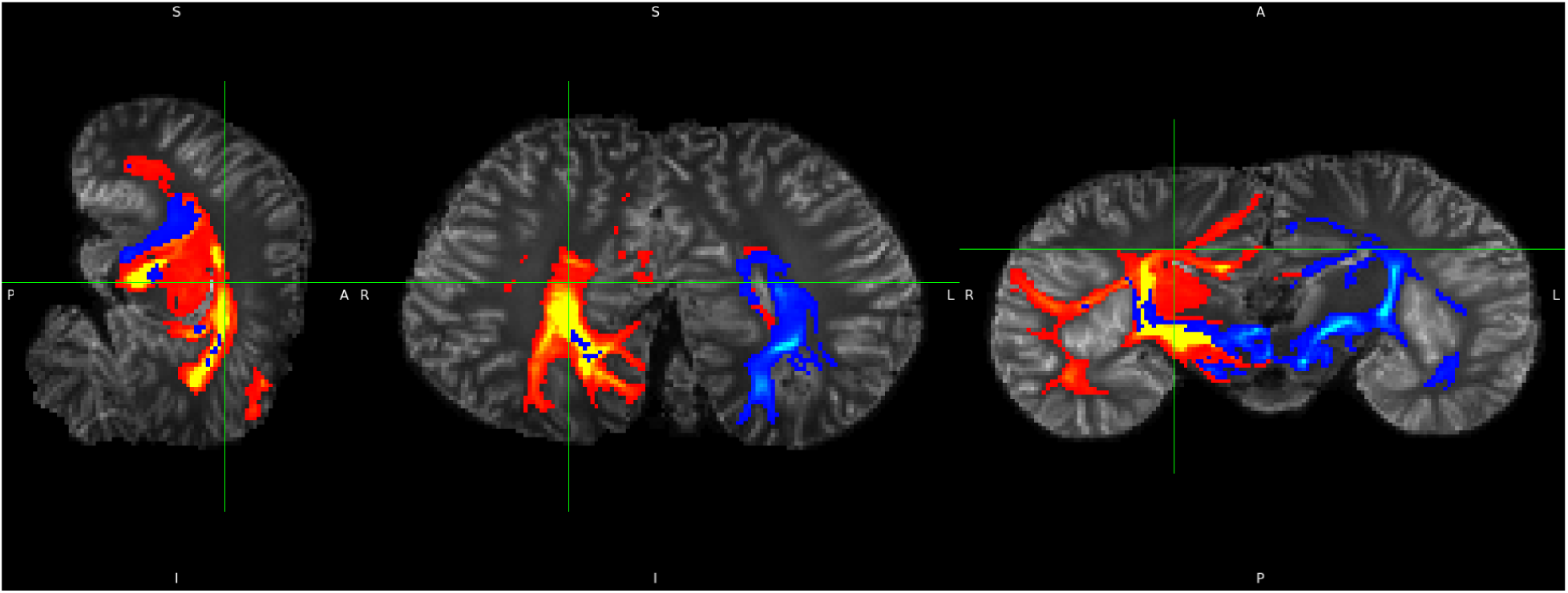
*L. acutus,* left IC tracts shown in blue, right IC tracts shown in red, set to a more liberal threshold of minimum 0.1% and maximum 5% of waytotals. Orthographic view.

**Figure S4C3:**
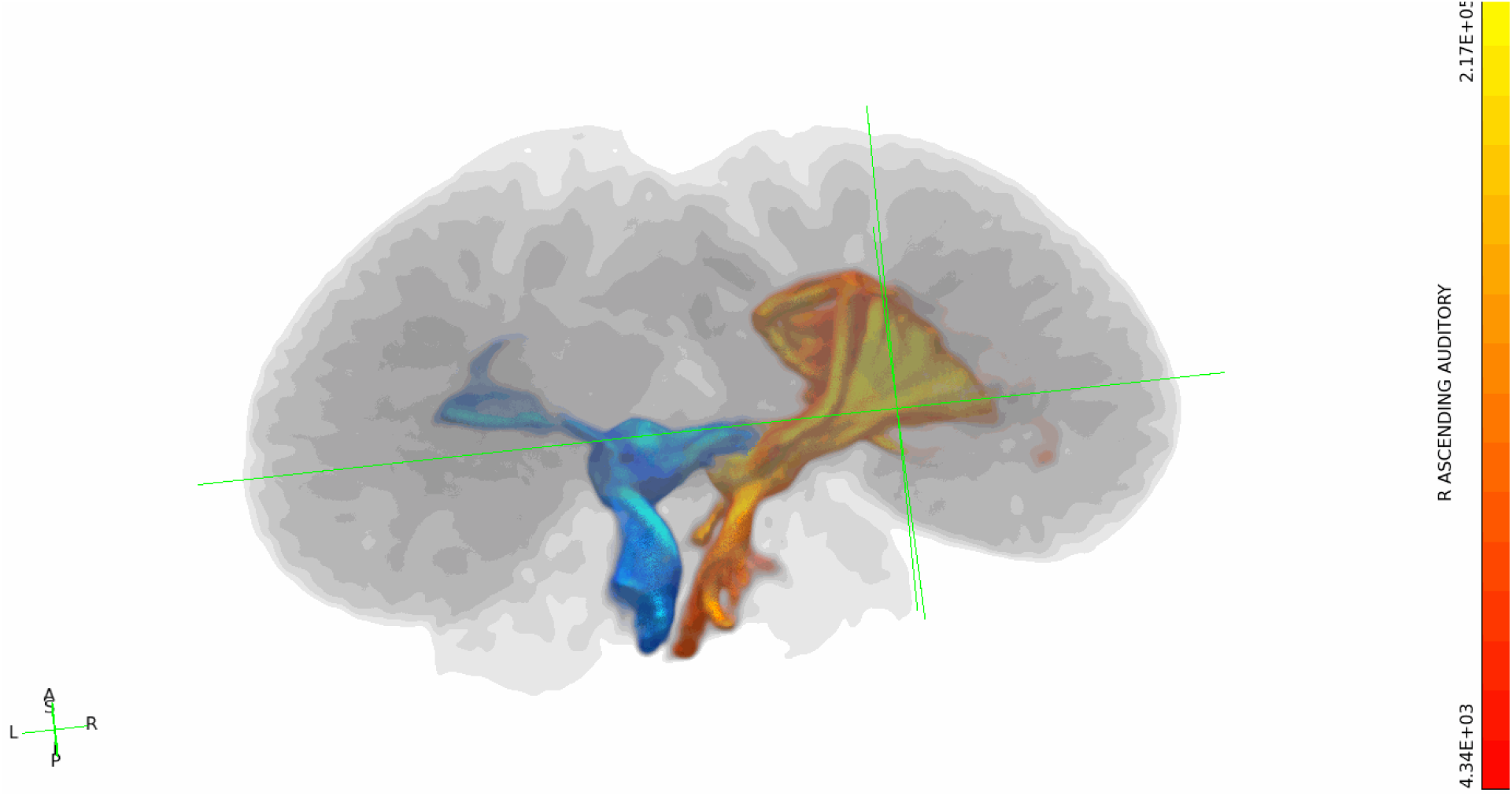
*L. acutus,* left IC tracts shown in blue, right IC tracts shown in red, set to a more liberal threshold of minimum 0.1% and maximum 5% of waytotals. Still 3-dimensional view.

**Figure S4C4:**
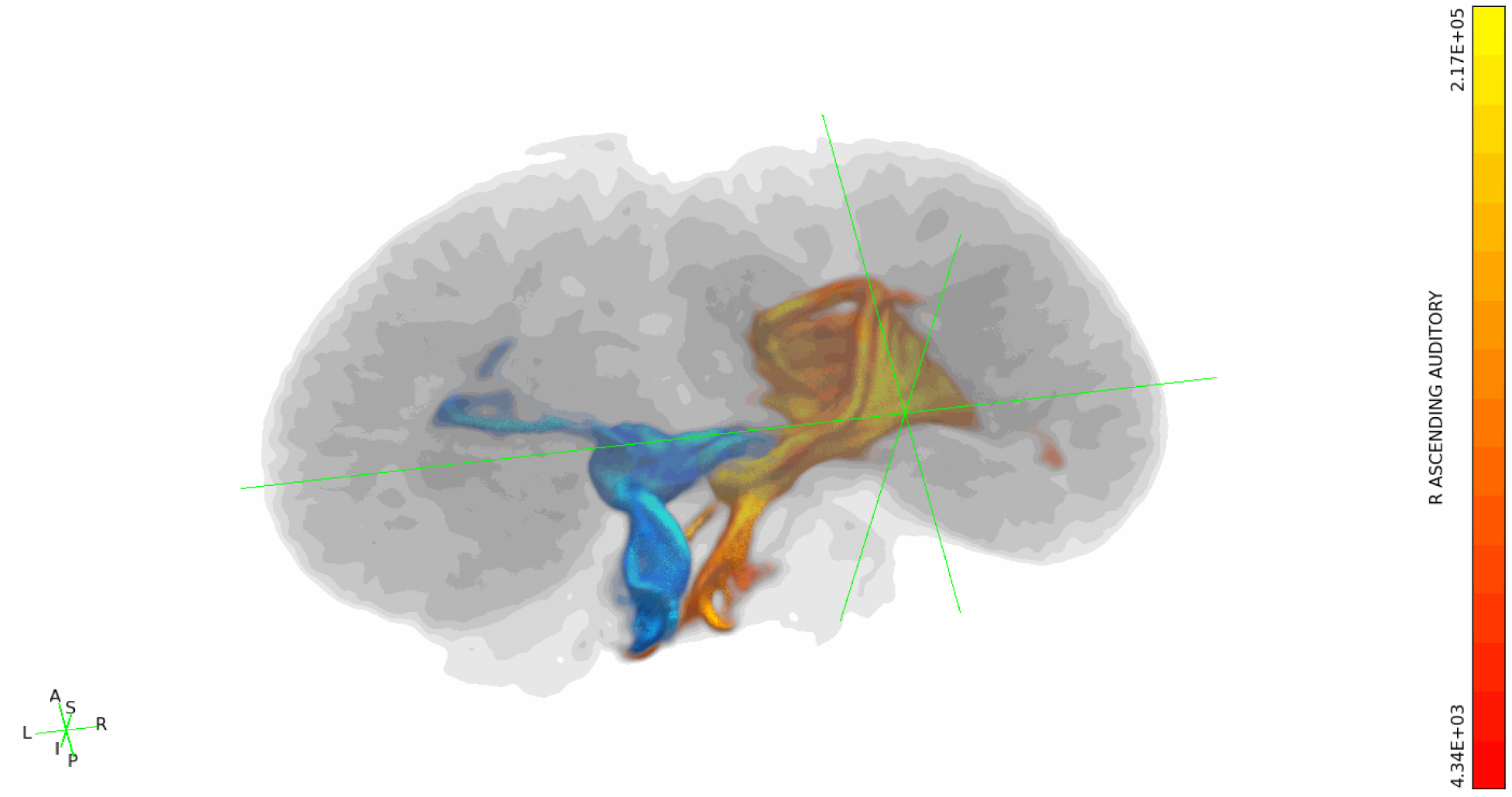
*L. acutus,* left IC tracts shown in blue, right IC tracts shown in red, set to a more liberal threshold of minimum 0.1% and maximum 5% of waytotals. Rotating 3-dimensional view.

**Figure S4D1:**
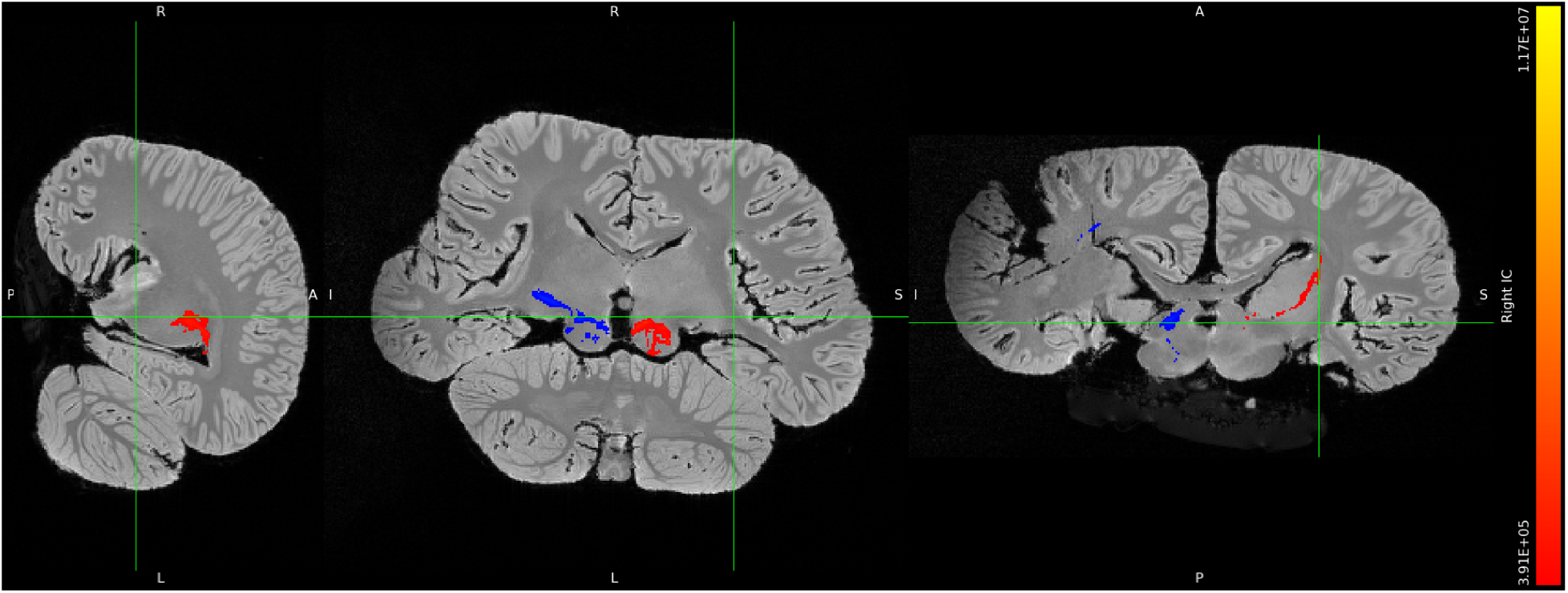
*B. borealis,* left IC tracts shown in blue, right IC tracts shown in red, minimum threshold set to 1% and maximum threshold set to 30% of waytotals. Orthographic view.

**Figure S4D2:**
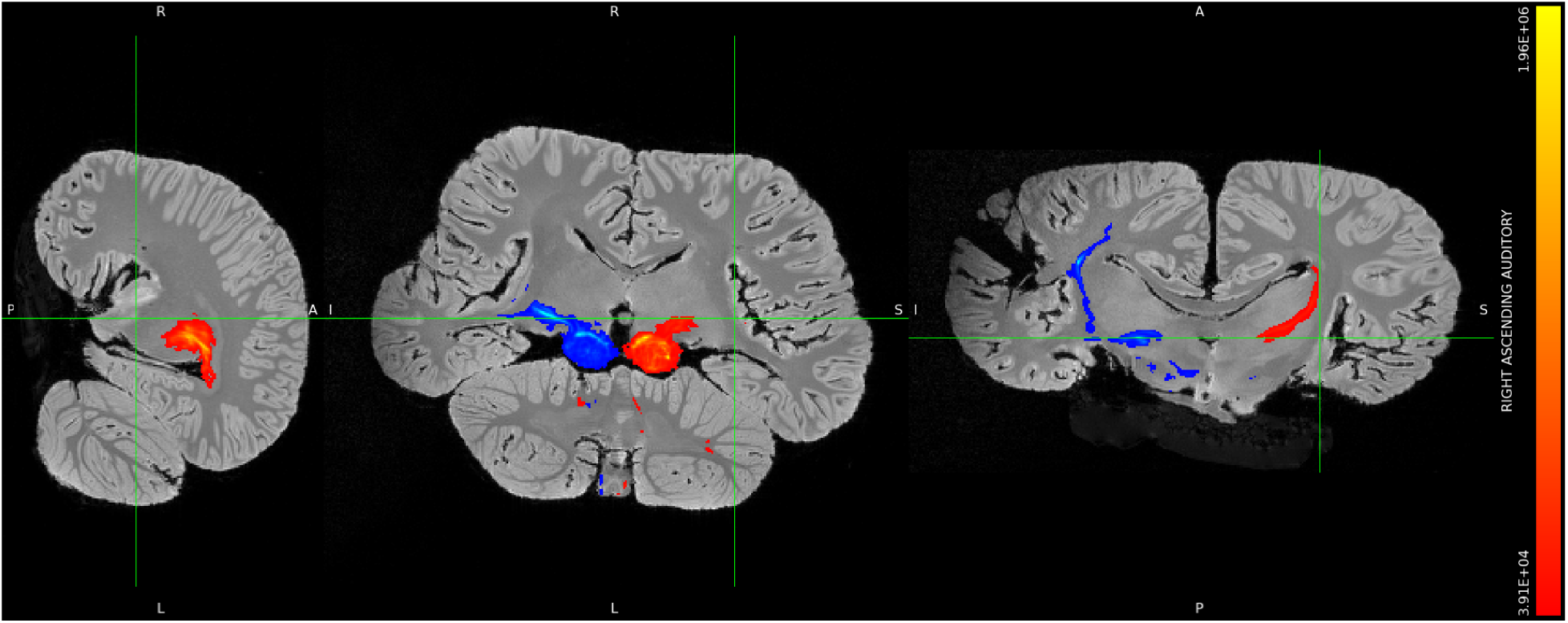
*B. borealis,* left IC tracts shown in blue, right IC tracts shown in red, set to a more liberal threshold of minimum 0.1% and maximum 5% of waytotals. Orthographic view.

**Figure S4D3:**
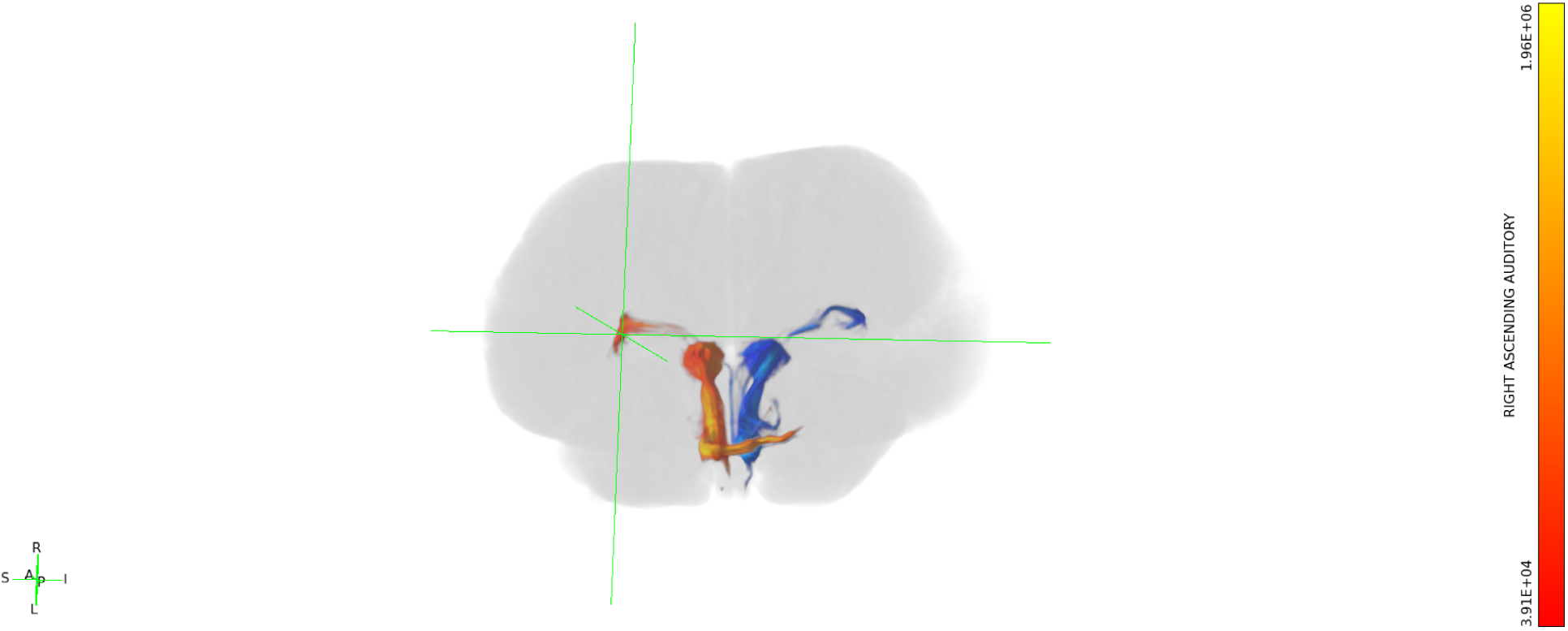
*B. borealis,* left IC tracts shown in blue, right IC tracts shown in red, set to a more liberal threshold of minimum 0.1% and maximum 5% of waytotals. Still 3-dimensional view.

**Figure S4D4:**
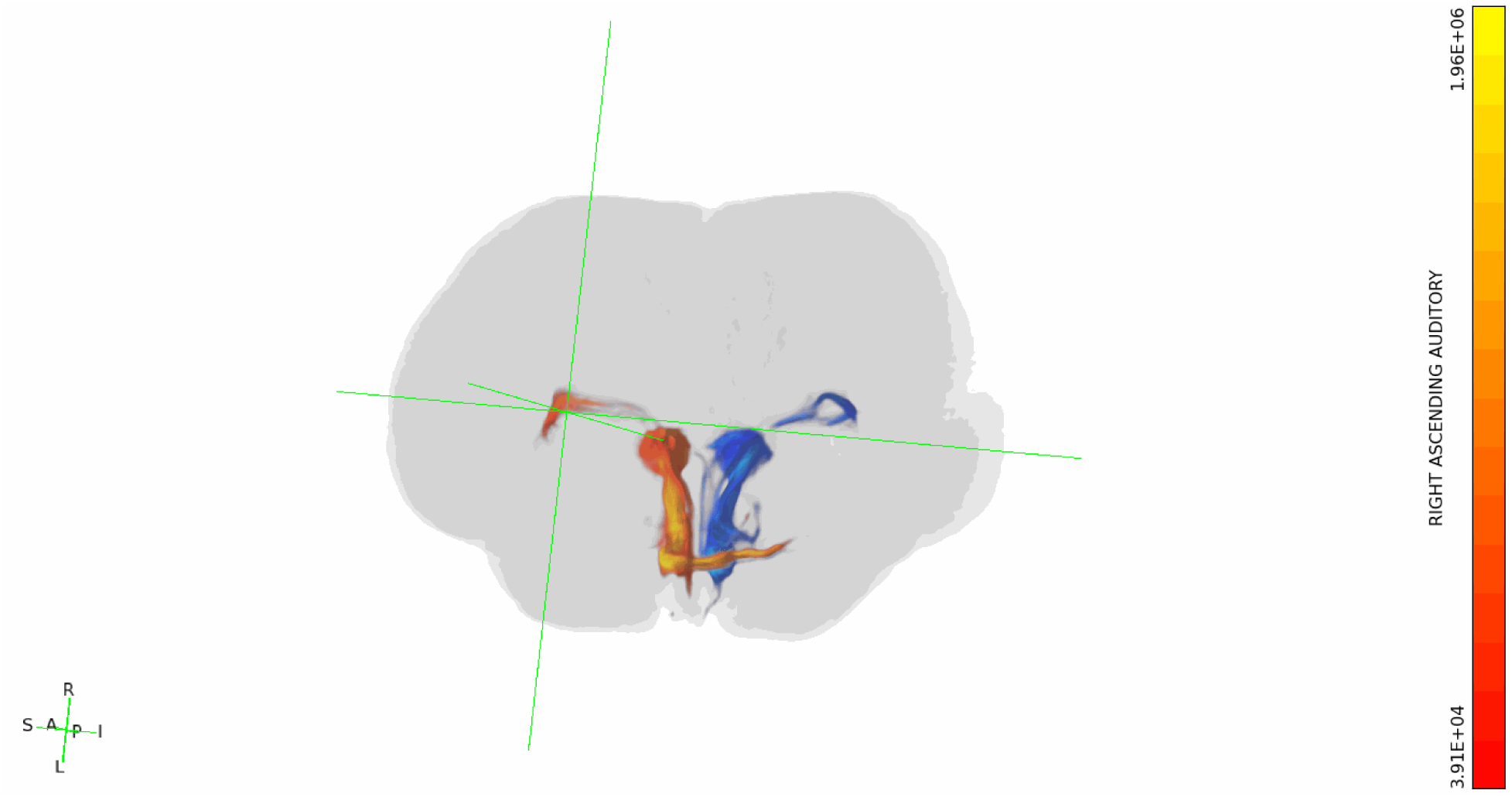
*B. borealis,* left IC tracts shown in blue, right IC tracts shown in red, set to a more liberal threshold of minimum 0.1% and maximum 5% of waytotals. Rotating 3-dimensional view.

### S5 Figures. Contralateral collicular-cortical tractograms

**Figure S5A1:**
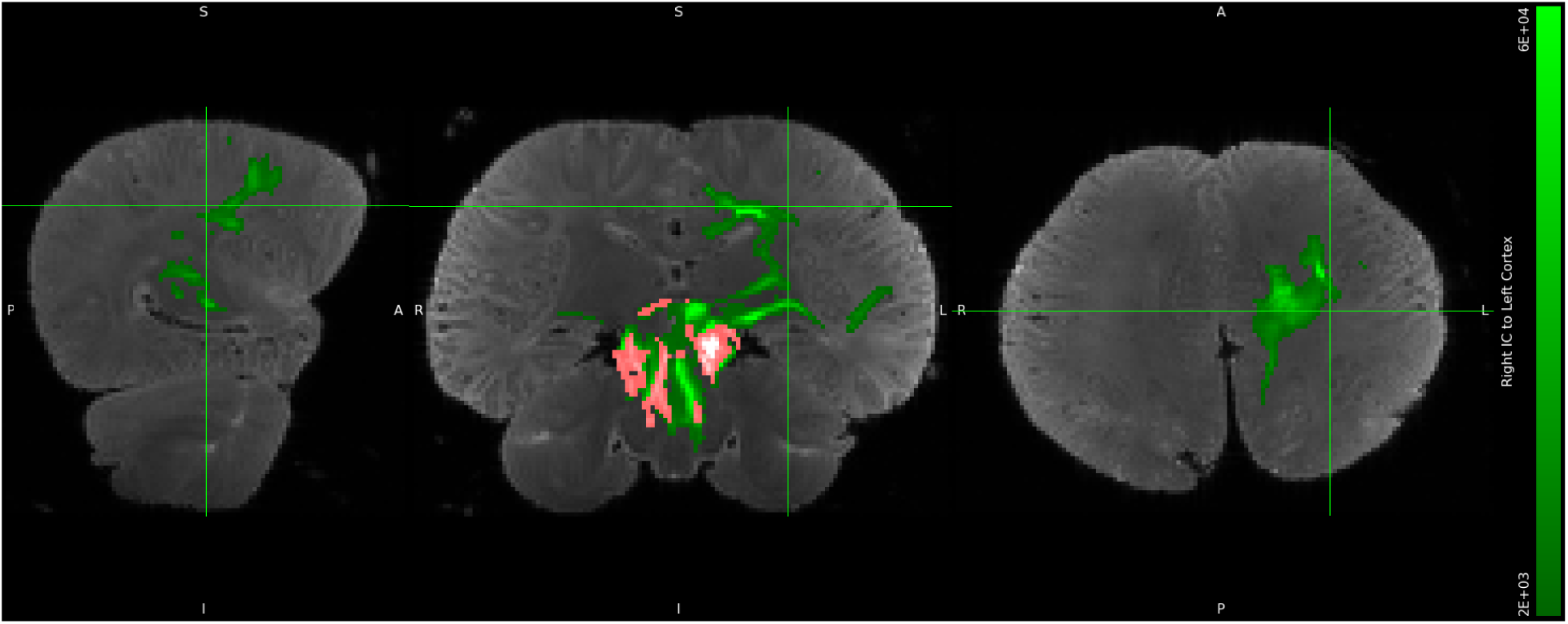
*D. delphis,* left IC to right cortex tracts shown in green, right IC to left cortex tracts shown in pink, minimum threshold set to 1% and maximum threshold set to 30% of waytotals. Both cerebella excluded. Orthographic view.

**Figure S5A2:**
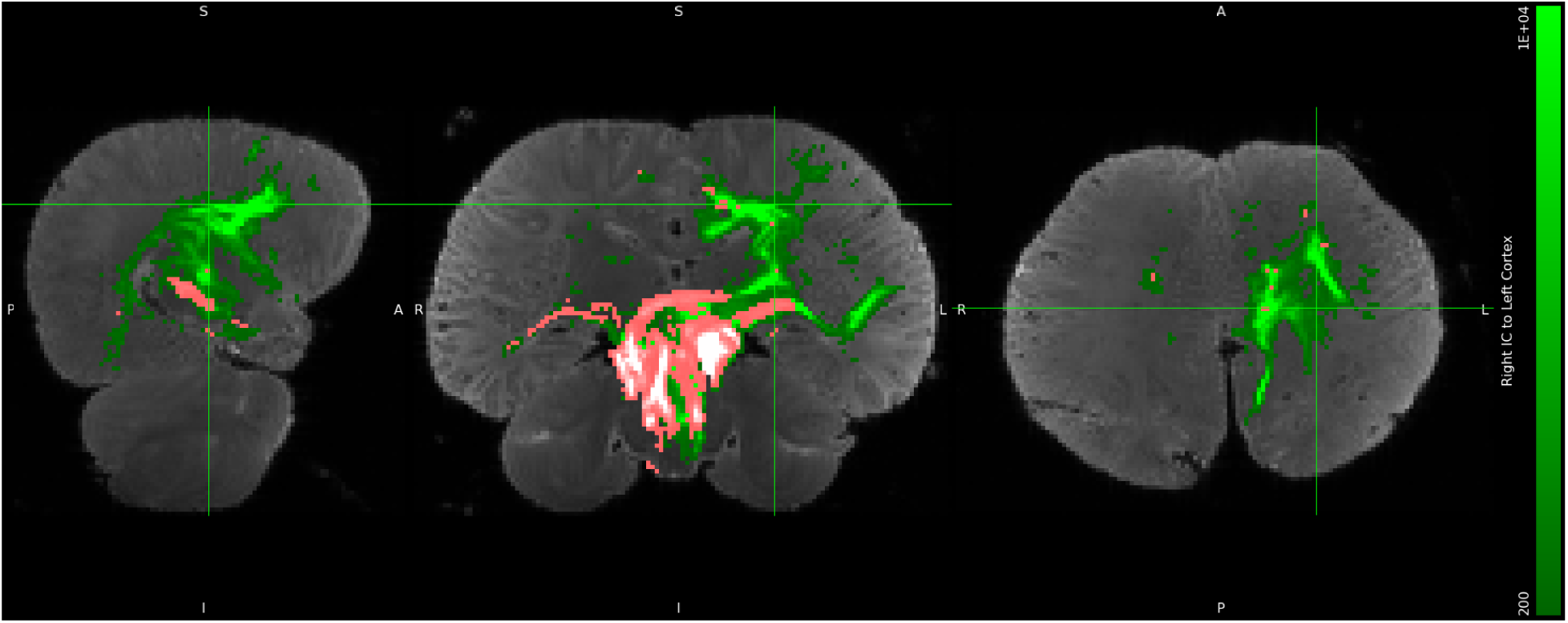
*D. delphis,* left IC to right cortex tracts shown in green, right IC to left cortex tracts shown in pink, set to a more liberal threshold of minimum 0.1% and maximum 5% of waytotals. Both cerebella excluded. Orthographic view.

**Figure S5A3:**
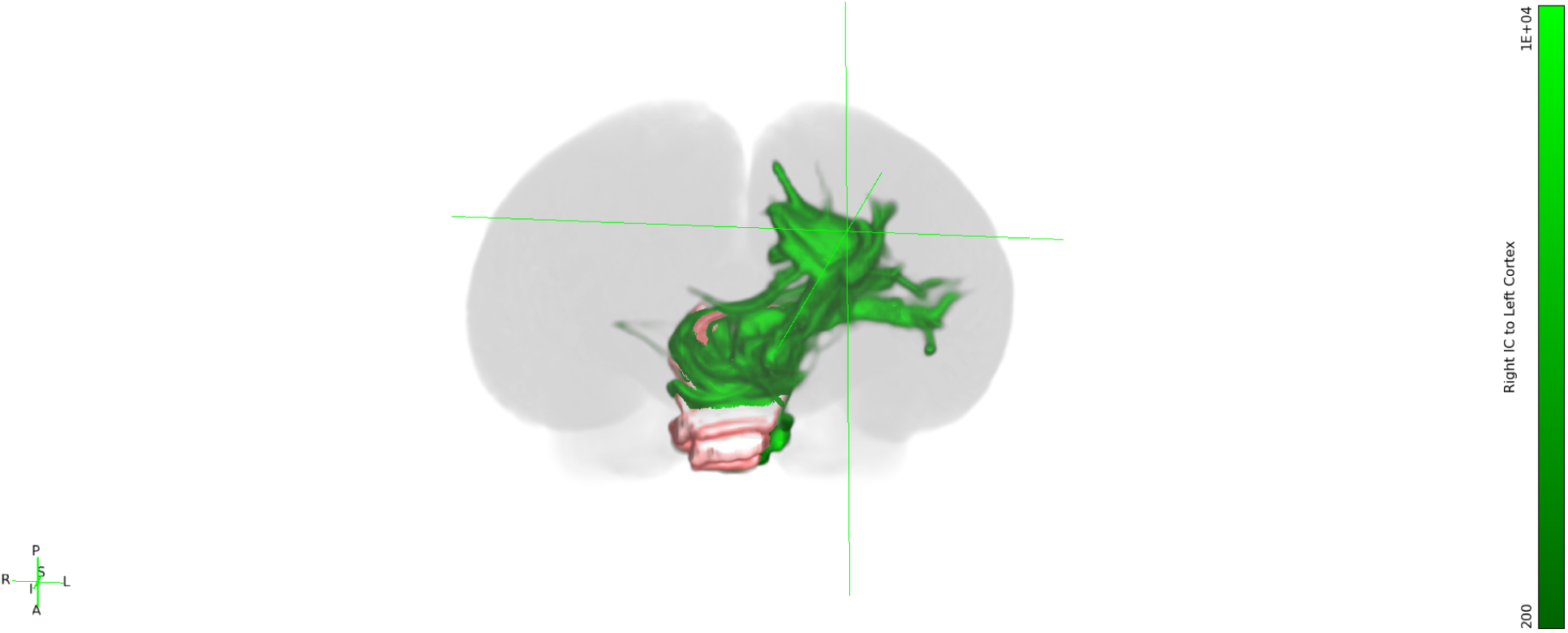
*D. delphis,* left IC to right cortex tracts shown in green, right IC to left cortex tracts shown in pink, set to a more liberal threshold of minimum 0.1% and maximum 5% of waytotals. Both cerebella excluded. Still 3-dimensional view.

**Figure S5A4:**
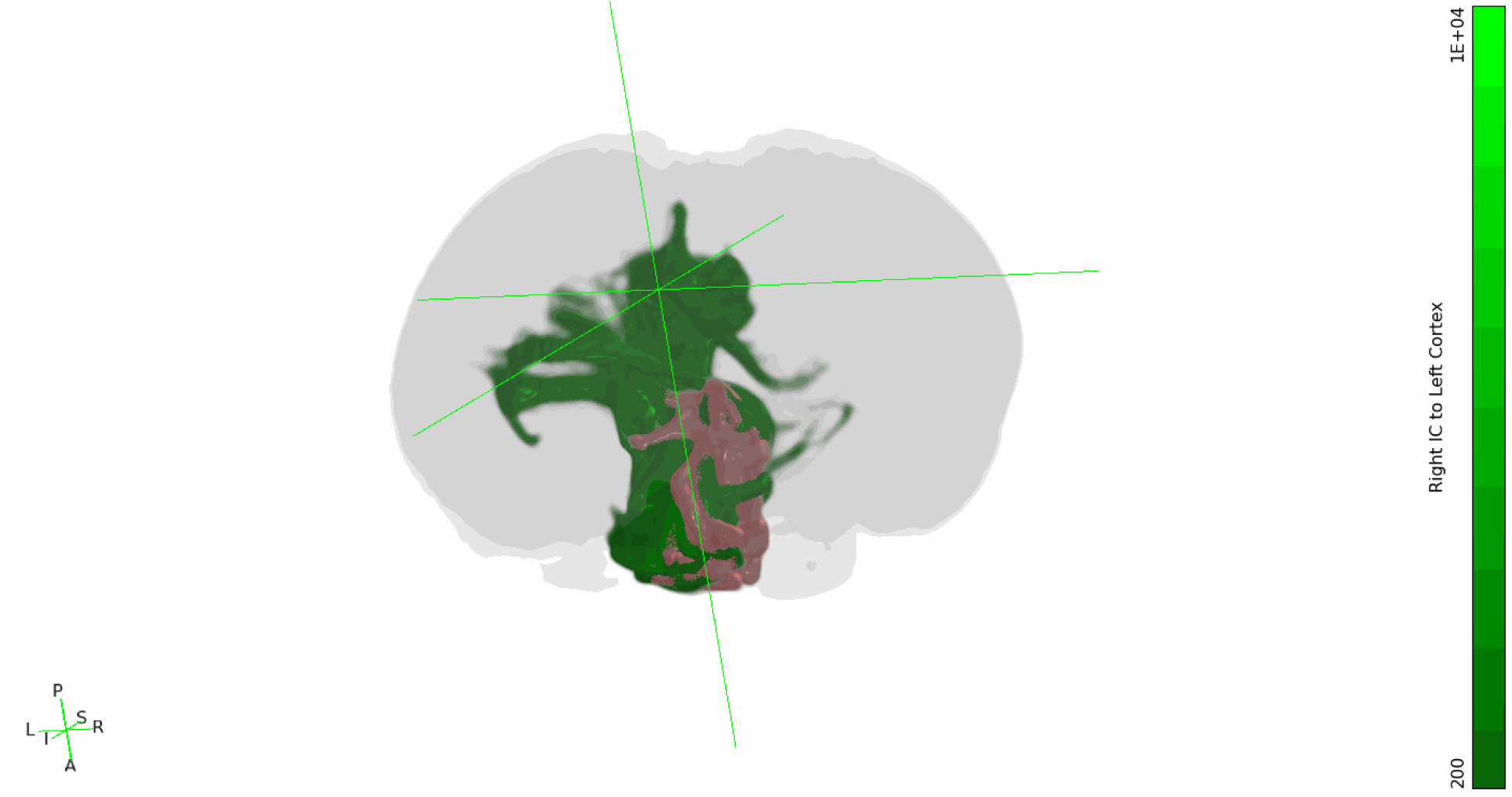
*D. delphis,* left IC to right cortex tracts shown in green, right IC to left cortex tracts shown in pink, set to a more liberal threshold of minimum 0.1% and maximum 5% of waytotals. Both cerebella excluded. Rotating 3-dimensional view.

**Figure S5B1:**
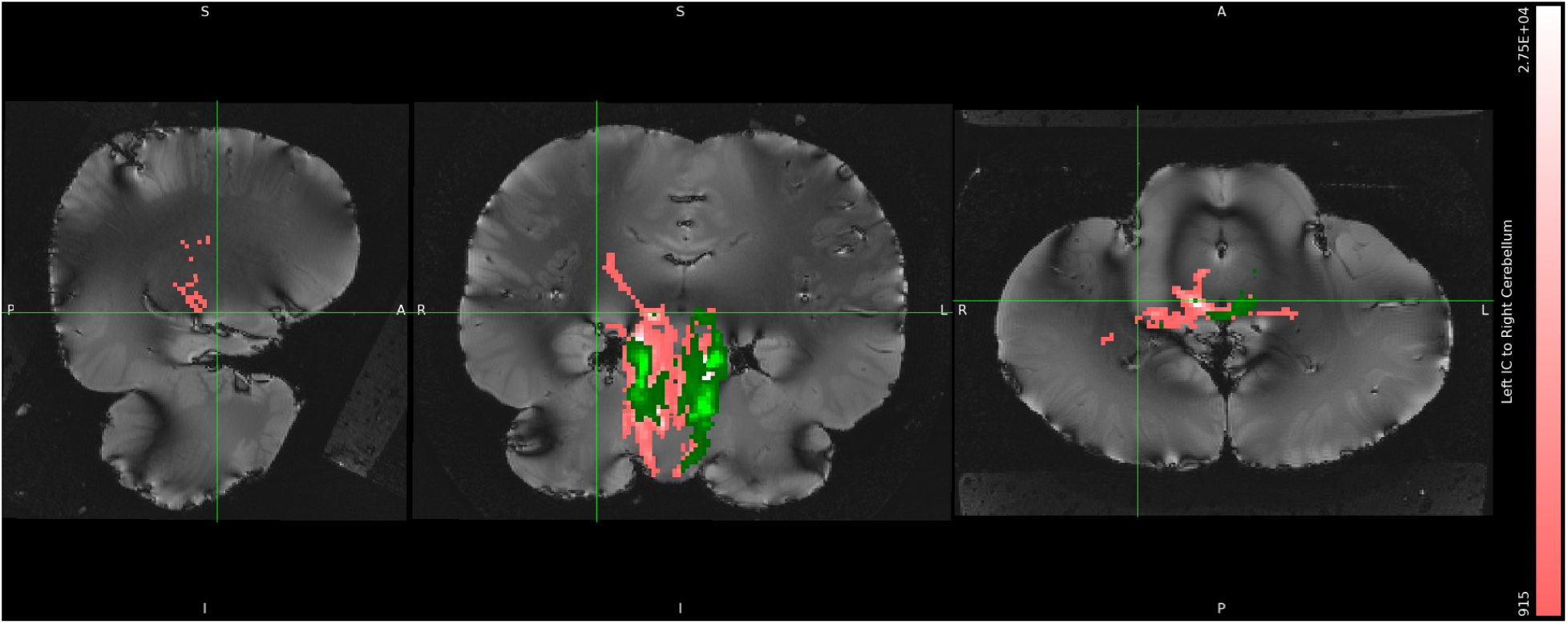
*S. attenuata,* left IC to right cortex tracts shown in green, right IC to left cortex tracts shown in pink, minimum threshold set to 1% and maximum threshold set to 30% of waytotals. Both cerebella excluded. Orthographic view.

**Figure S5B2:**
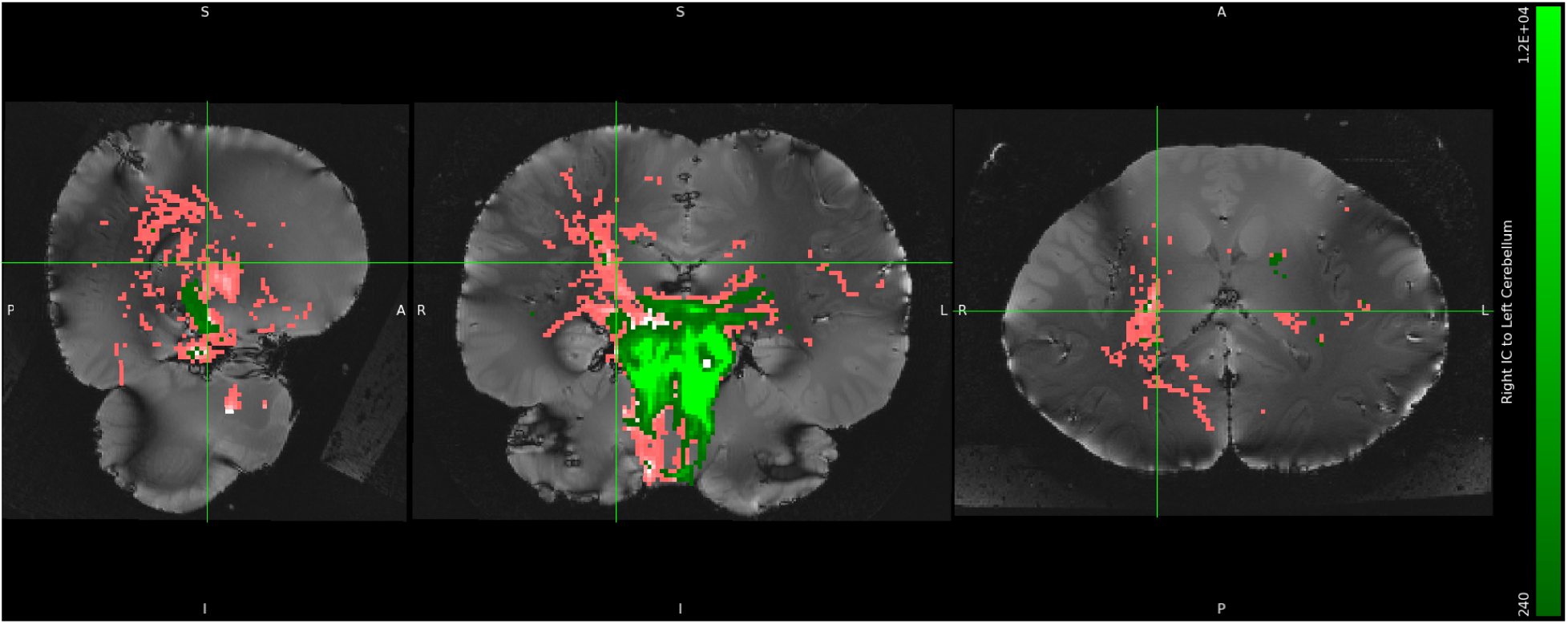
*S. attenuata,* left IC to right cortex tracts shown in green, right IC to left cortex tracts shown in pink, set to a more liberal threshold of minimum 0.1% and maximum 5% of waytotals. Both cerebella excluded. Orthographic view.

**Figure S5B3:**
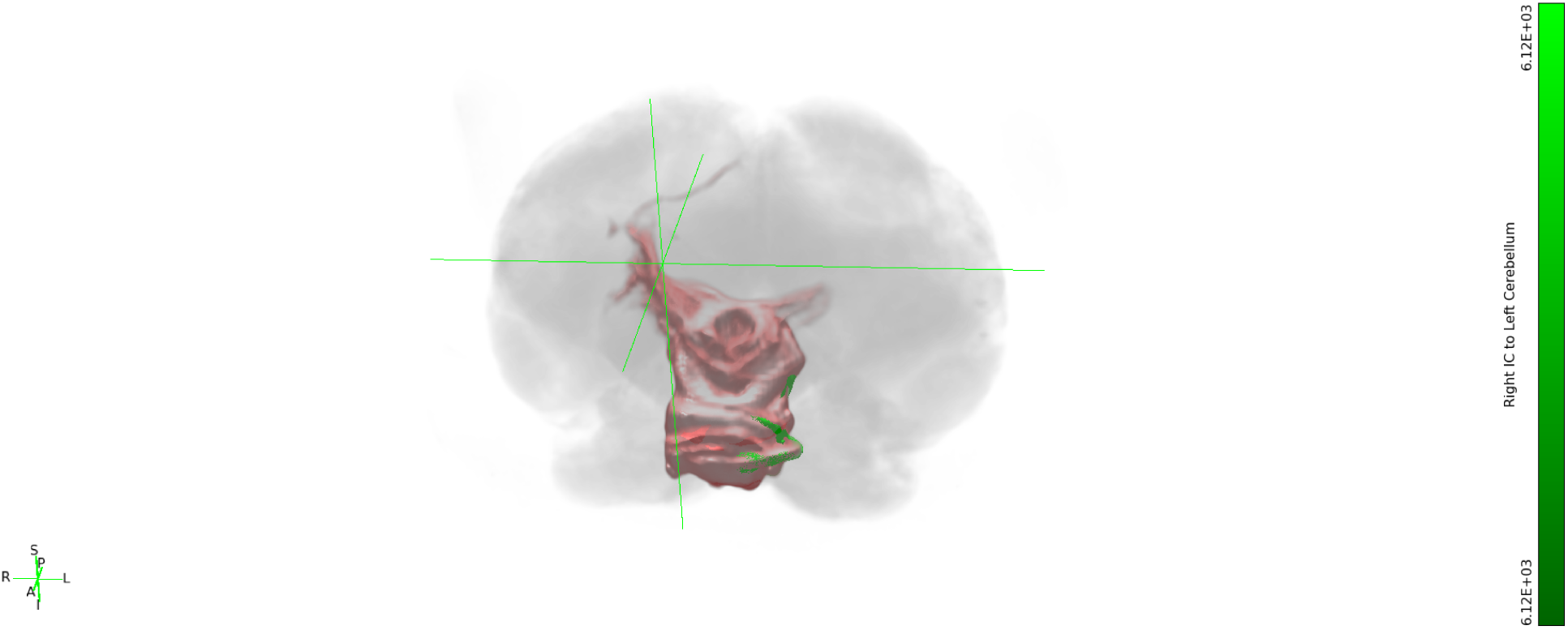
*S. attenuata,* left IC to right cortex tracts shown in green, right IC to left cortex tracts shown in pink, set to a more liberal threshold of minimum 0.1% and maximum 5% of waytotals. Both cerebella excluded. Still 3-dimensional view.

**Figure S5B4:**
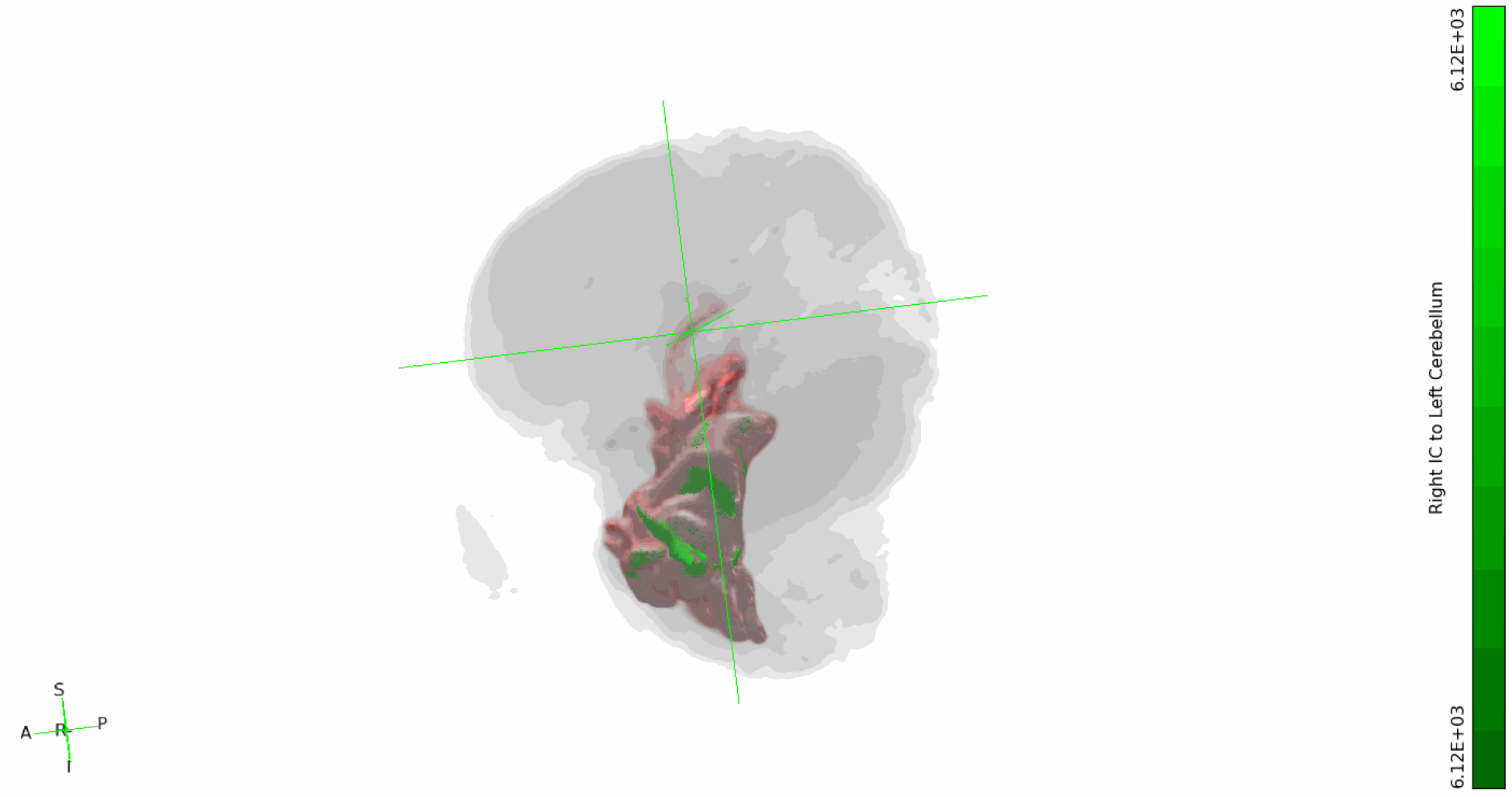
*S. attenuata,* left IC to right cortex tracts shown in green, right IC to left cortex tracts shown in pink, set to a more liberal threshold of minimum 0.1% and maximum 5% of waytotals. Both cerebella excluded. Rotating 3-dimensional view.

**Figure S5C1:**
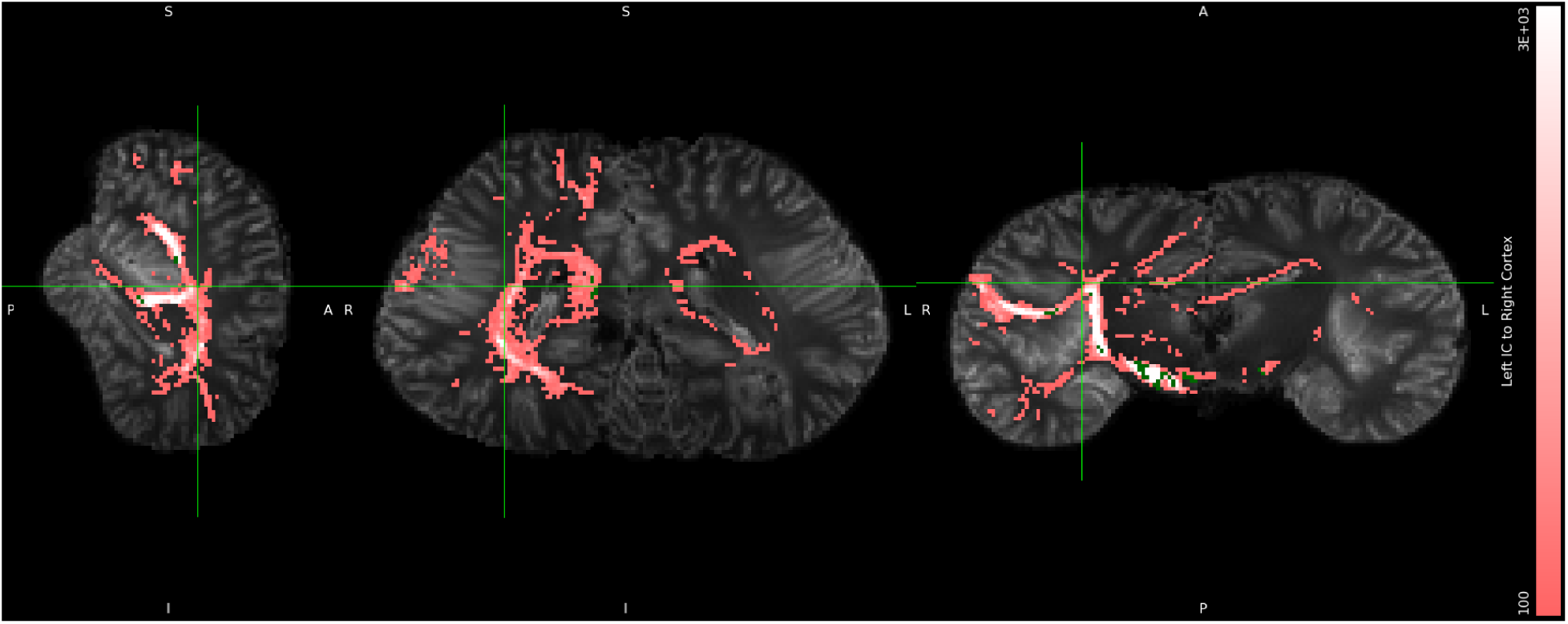
*L. acutus,* left IC to right cortex tracts shown in green, right IC to left cortex tracts shown in pink, minimum threshold set to 1% and maximum threshold set to 30% of waytotals. Both cerebella excluded. Orthographic view.

**Figure S5C2i:**
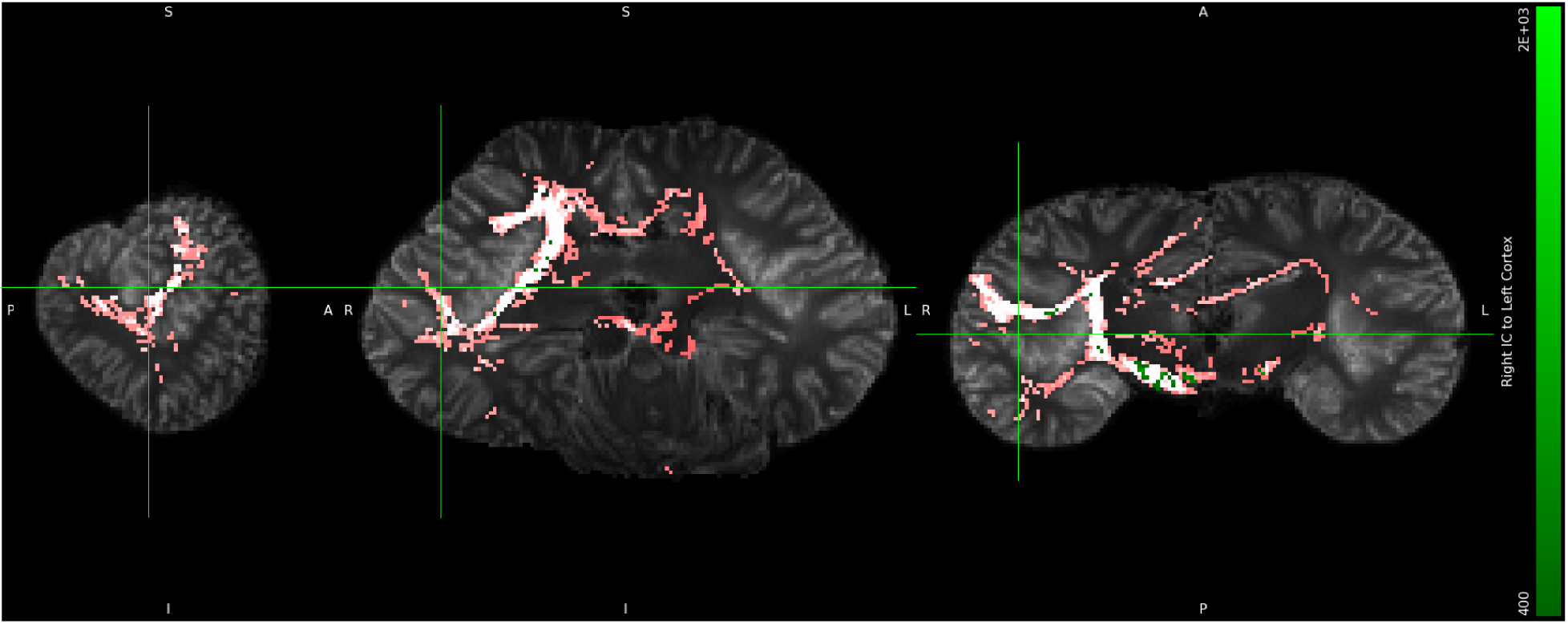
*L. acutus,* left IC to right cortex tracts shown in green, right IC to left cortex tracts shown in pink, set to a more liberal threshold of minimum 0.1% and maximum 5% of waytotals. Both cerebella excluded. Orthographic view.

**Figure S5C3:**
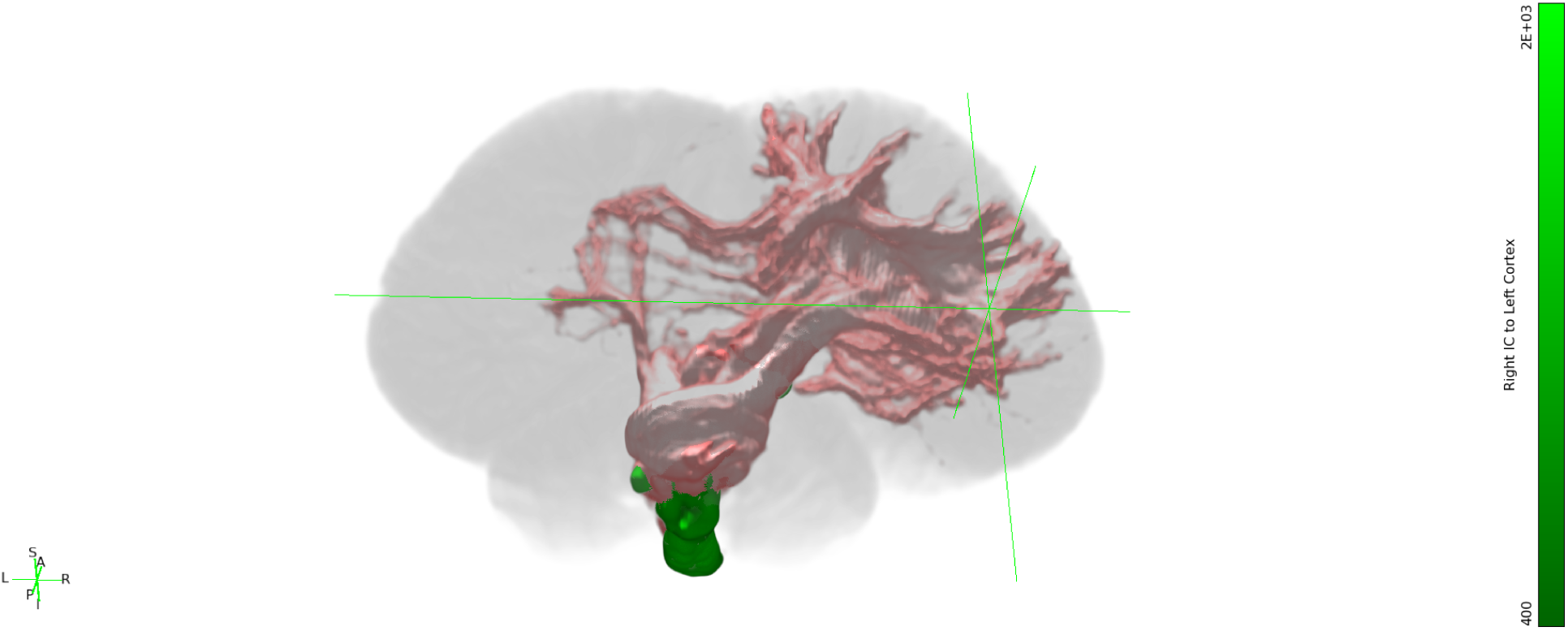
*L. acutus,* left IC to right cortex tracts shown in green, right IC to left cortex tracts shown in pink, set to a more liberal threshold of minimum 0.1% and maximum 5% of waytotals. Both cerebella excluded. Still 3-dimensional view.

**Figure S5C4:**
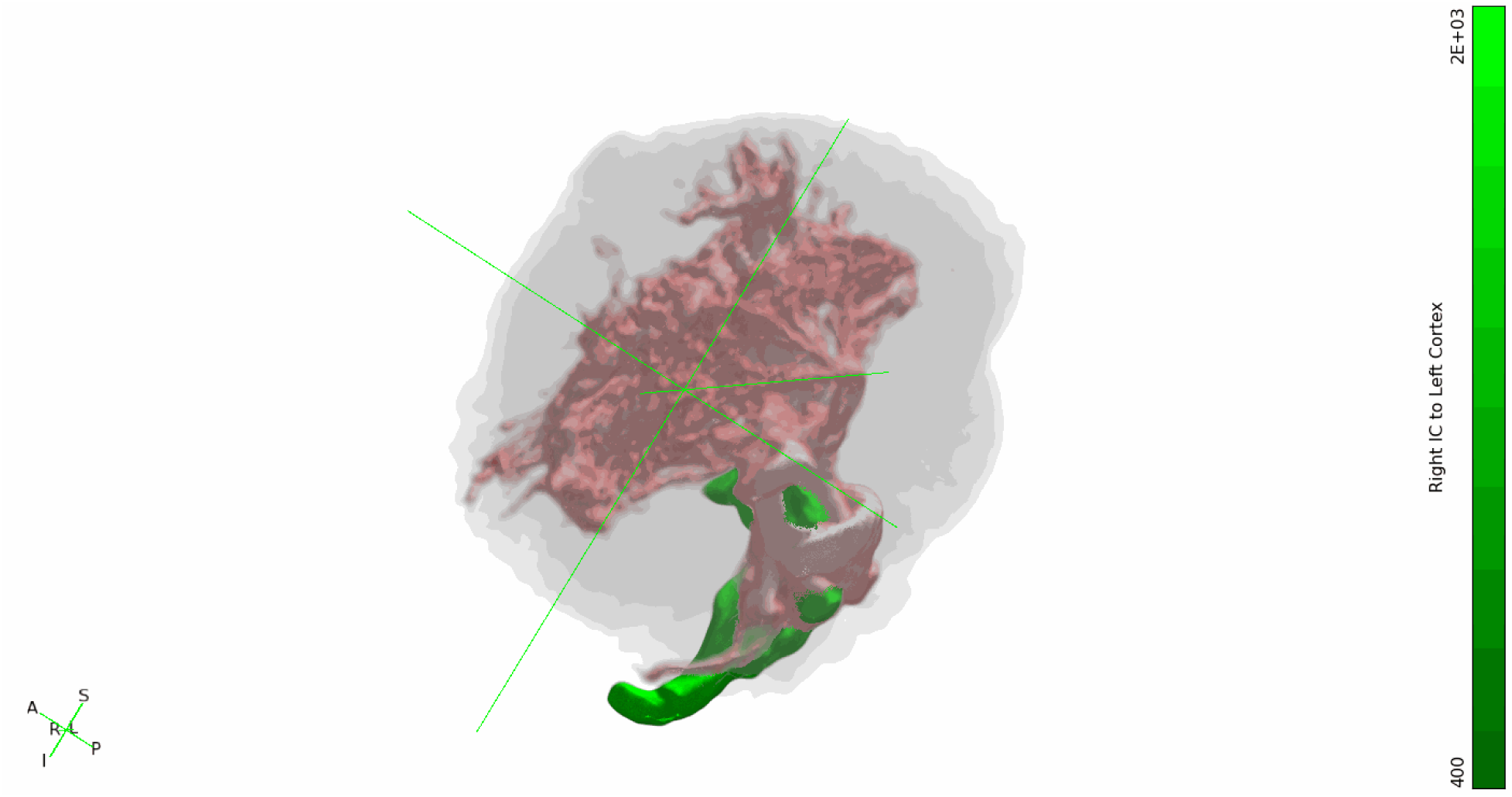
*L. acutus,* left IC to right cortex tracts shown in green, right IC to left cortex tracts shown in pink, set to a more liberal threshold of minimum 0.1% and maximum 5% of waytotals. Both cerebella excluded. Rotating 3-dimensional view.

**Figure S5D1:**
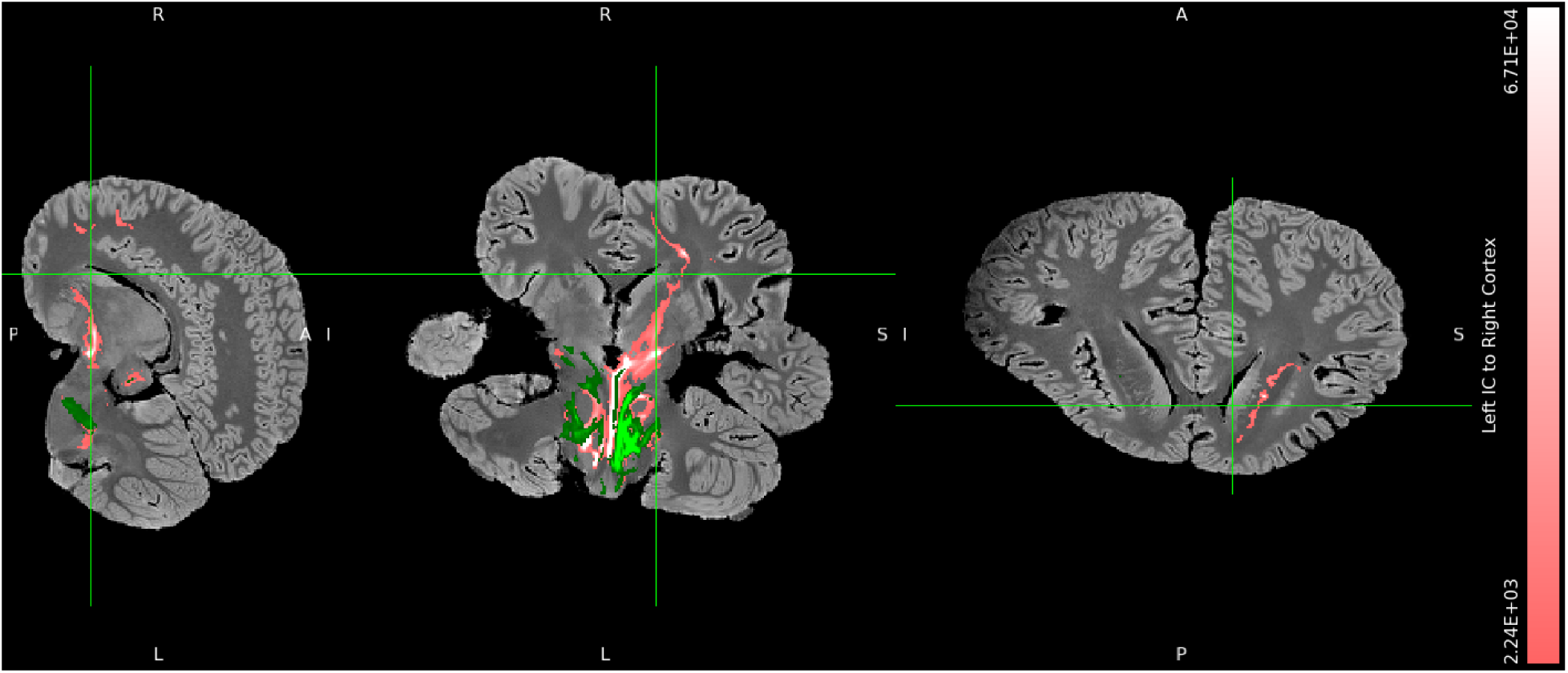
*B. borealis,* left IC to right cortex tracts shown in green, right IC to left cortex tracts shown in pink, minimum threshold set to 1% and maximum threshold set to 30% of waytotals. Both cerebella excluded. Orthographic view.

**Figure S5D2:**
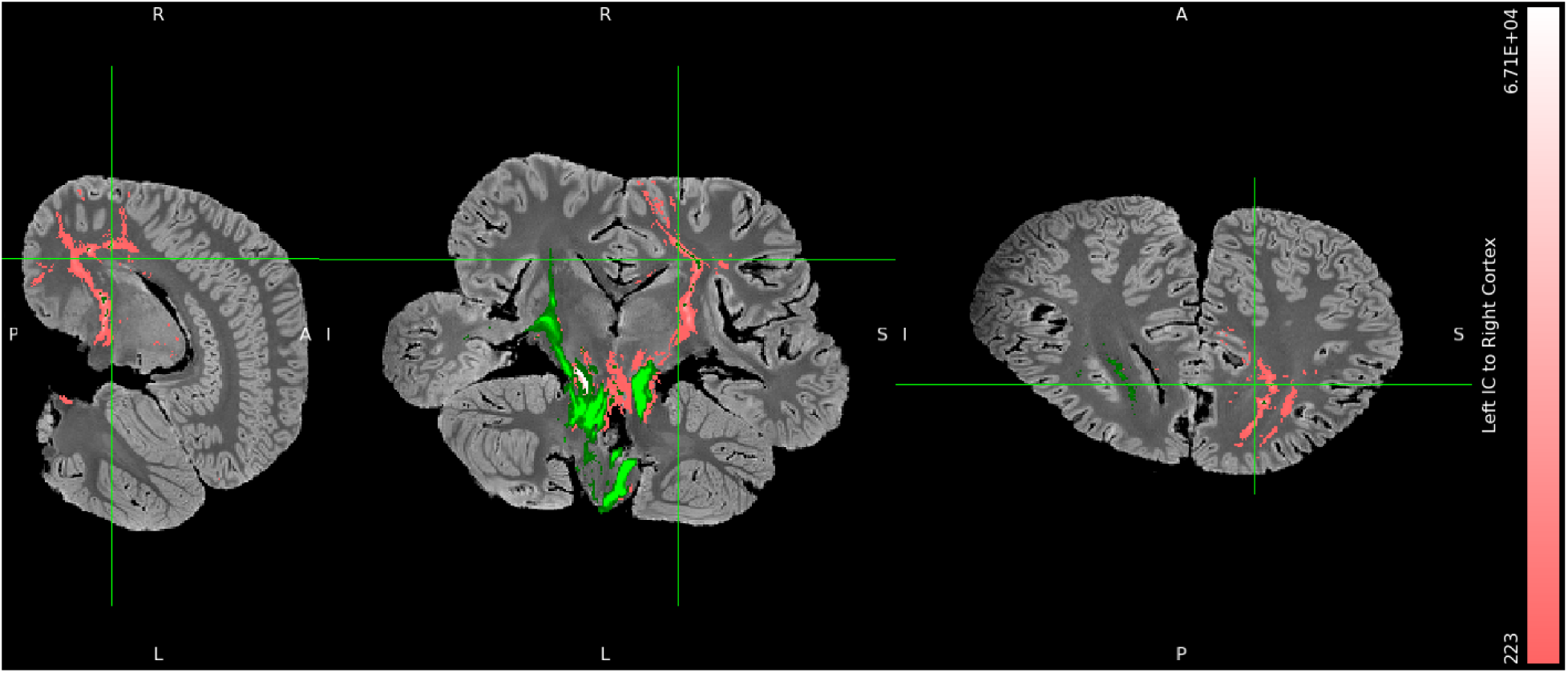
*B. borealis,* left IC to right cortex tracts shown in green, right IC to left cortex tracts shown in pink, set to a more liberal threshold of minimum 0.1% and maximum 5% of waytotals. Both cerebella excluded. Orthographic view.

**Figure S5D3:**
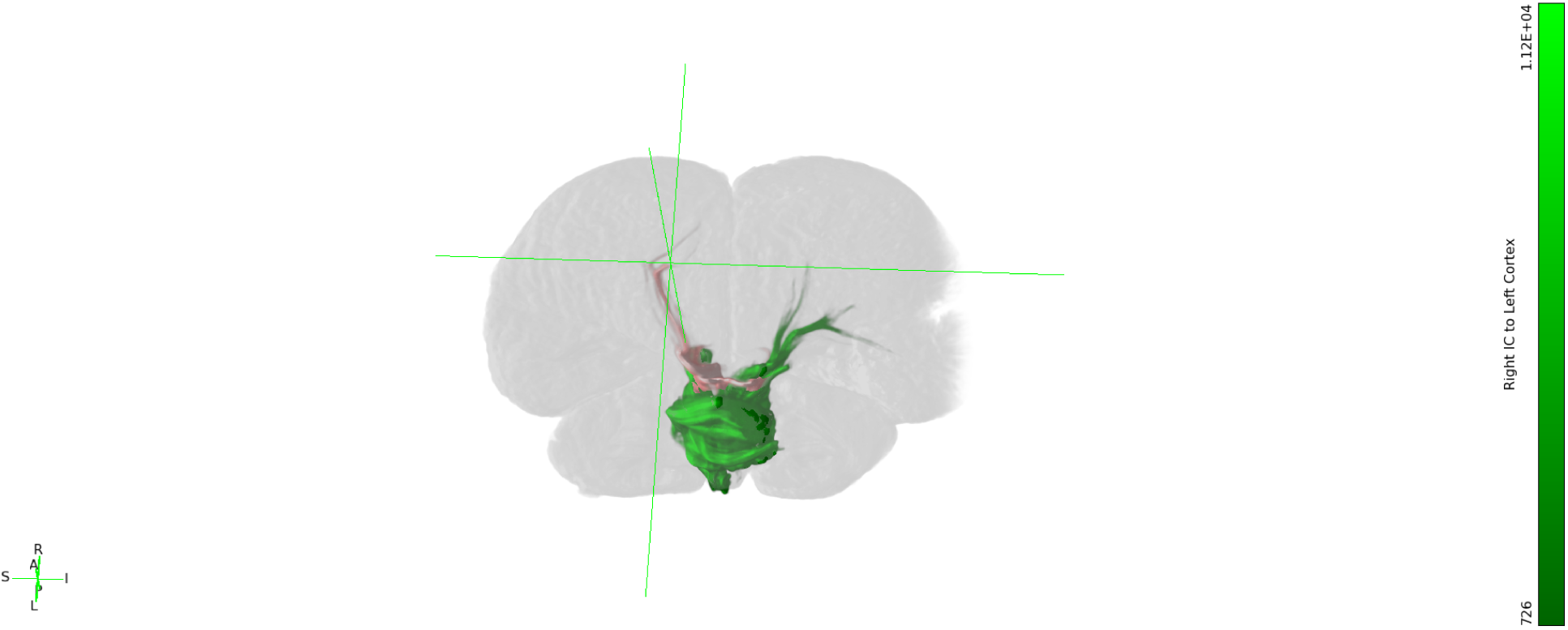
*B. borealis,* left IC to right cortex tracts shown in green, right IC to left cortex tracts shown in pink, set to a more liberal threshold of minimum 0.1% and maximum 5% of waytotals. Both cerebella excluded. Still 3-dimensional view.

**Figure S5D4:**
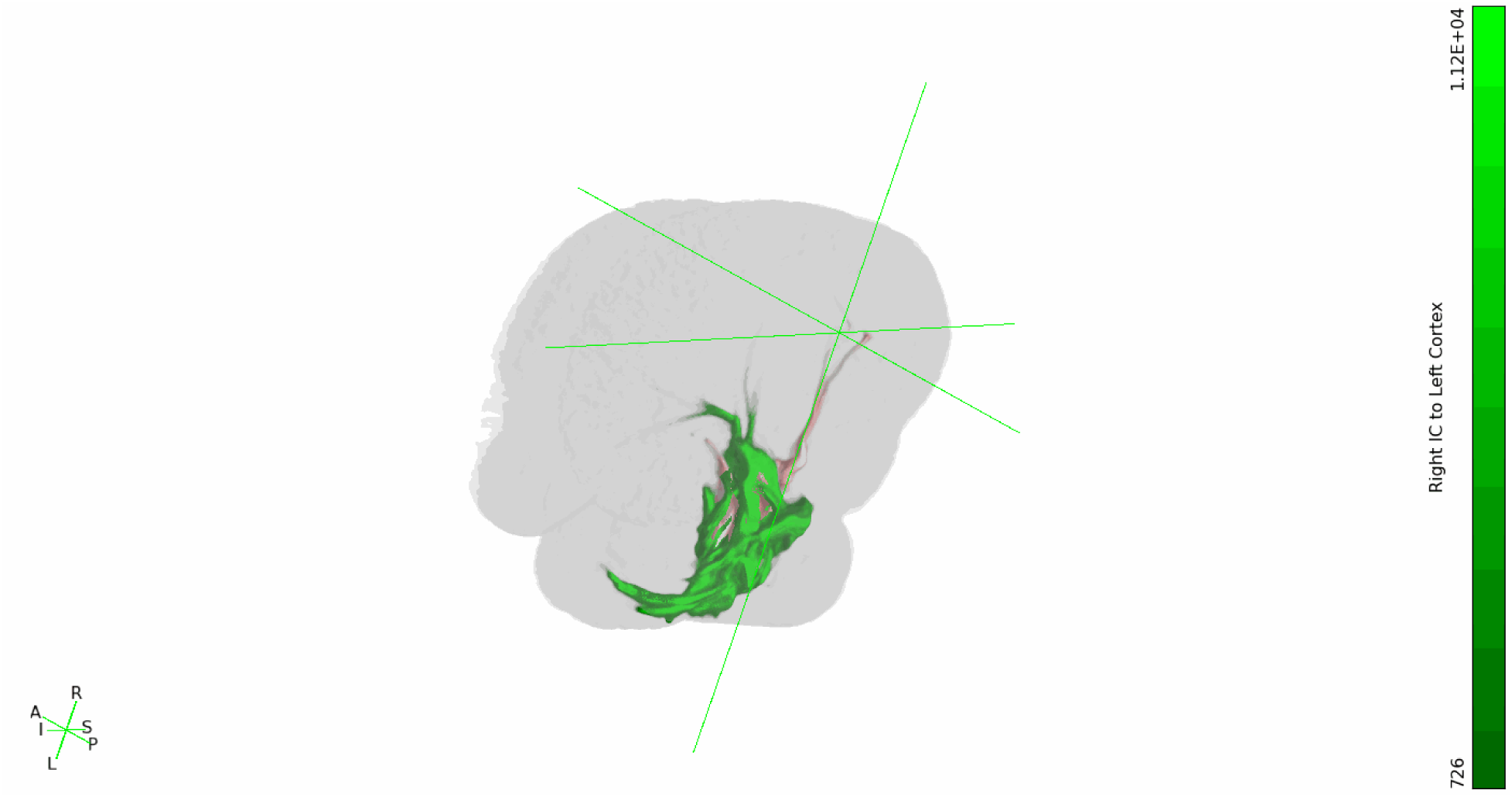
*B. borealis,* left IC to right cortex tracts shown in green, right IC to left cortex tracts shown in pink, set to a more liberal threshold of minimum 0.1% and maximum 5% of waytotals. Both cerebella excluded. Rotating 3-dimensional view.

### S6 Figures. Contralateral collicular-cerebellar tractograms

**Figure S6A1.**
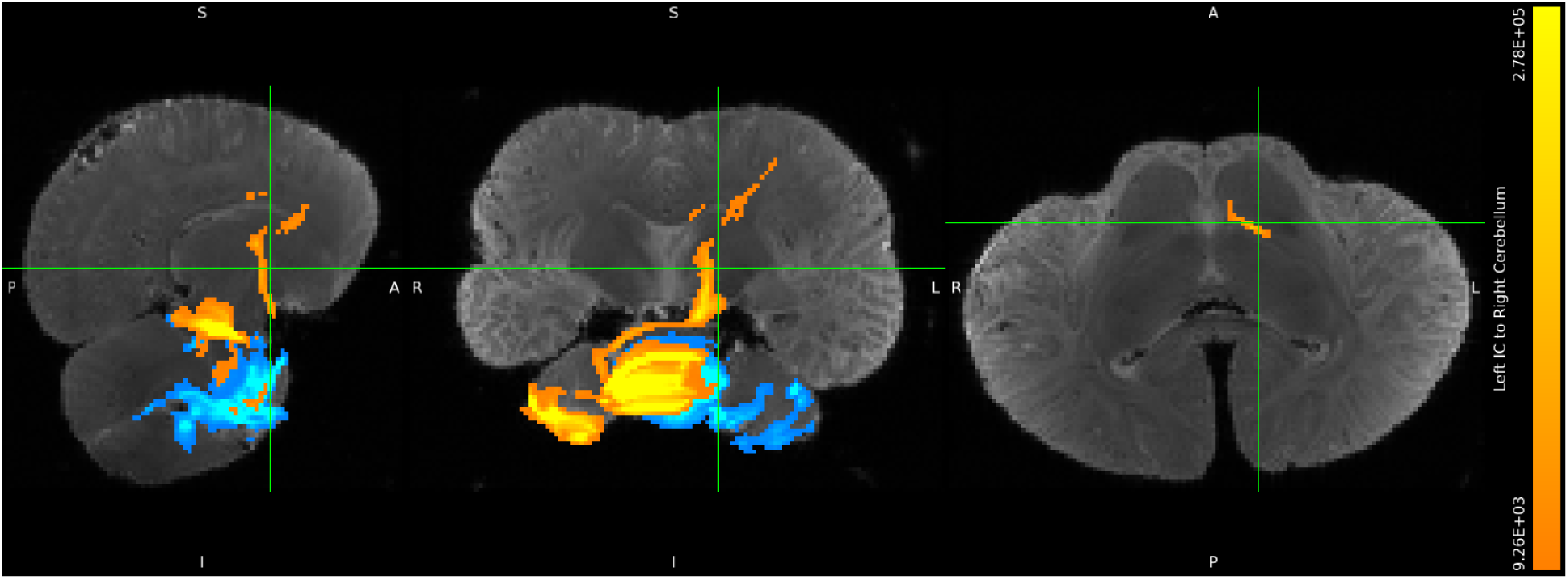
*D. delphis,* left IC to right cerebellum tracts shown in orange, right IC to left cerebellum tracts shown in turquoise, minimum threshold set to 1% and maximum threshold set to 30% of waytotals. Orthographic view.

**Figure S6A2.**
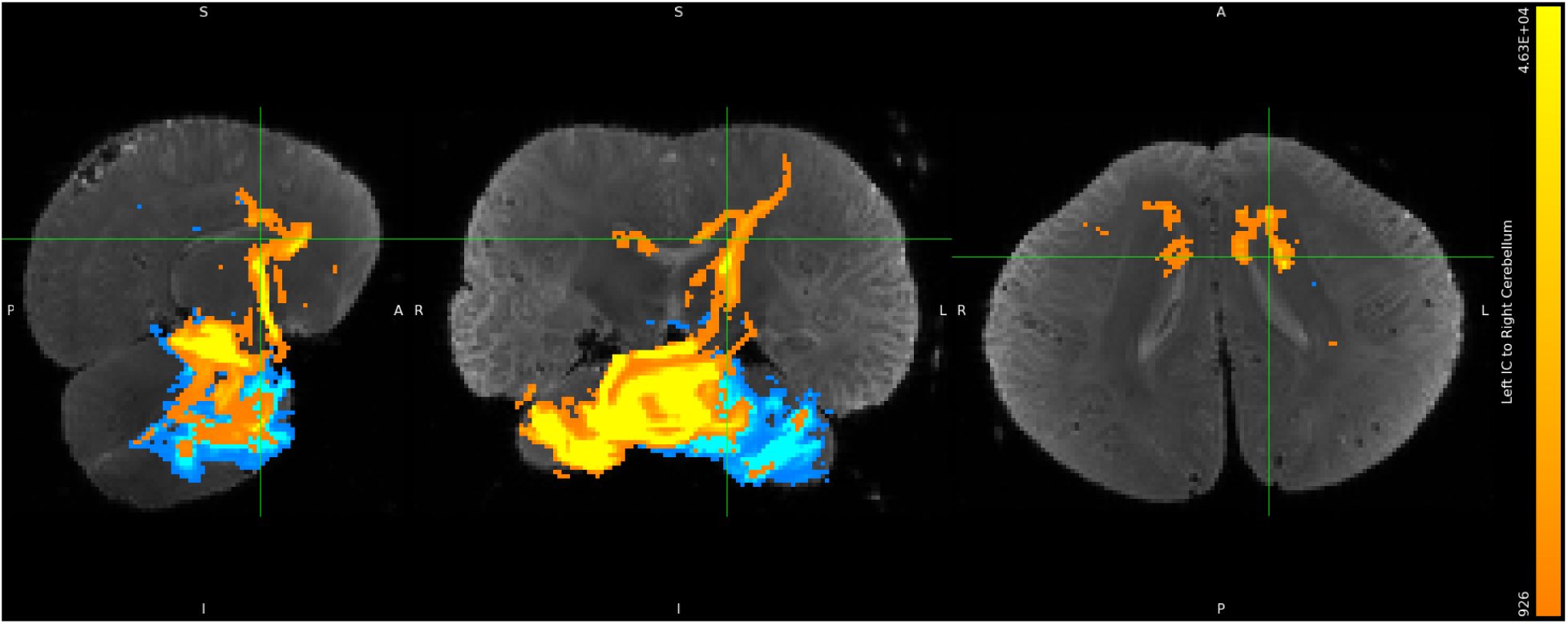
*D. delphis,* left IC to right cerebellum tracts shown in orange, right IC to left cerebellum tracts shown in turquoise, set to a more liberal threshold of minimum 0.1% and maximum 5% of waytotals. Orthographic view.

**Figure S6A3.**
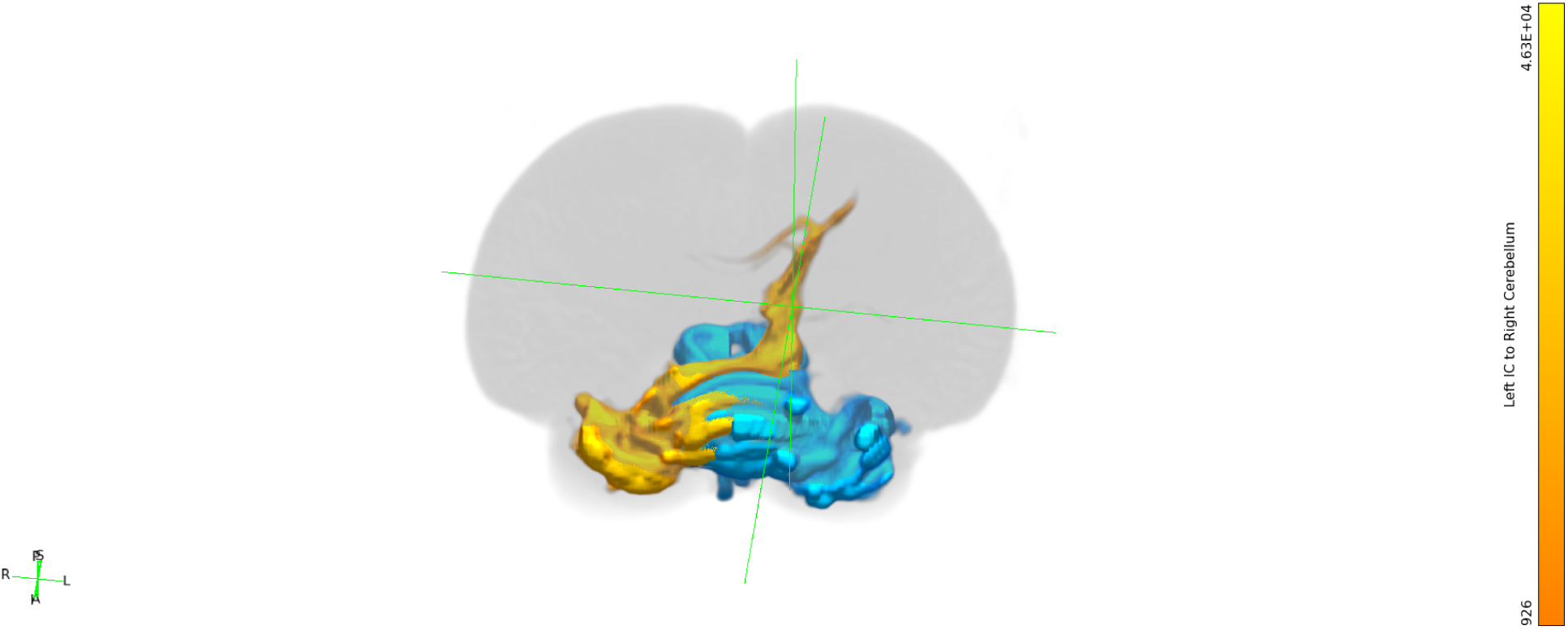
*D. delphis,* left IC to right cerebellum tracts shown in orange, right IC to left cerebellum tracts shown in turquoise, set to a more liberal threshold of minimum 0.1% and maximum 5% of waytotals. Still 3-dimensional view.

**Figure S6A4.**
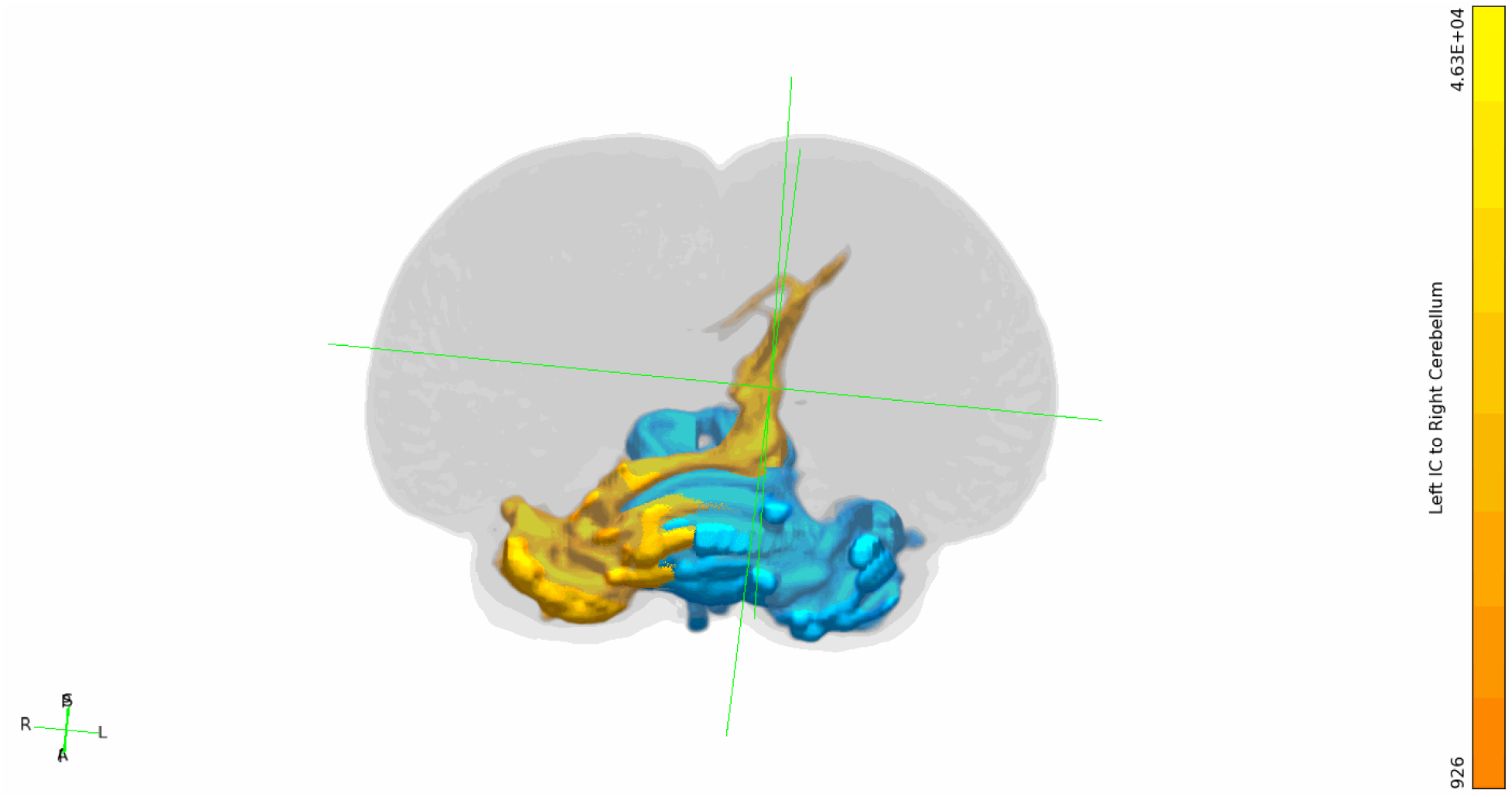
*D. delphis,* left IC to right cerebellum tracts shown in orange, right IC to left cerebellum tracts shown in turquoise, set to a more liberal threshold of minimum 0.1% and maximum 5% of waytotals. Rotating 3-dimensional view.

**Figure S6B1.**
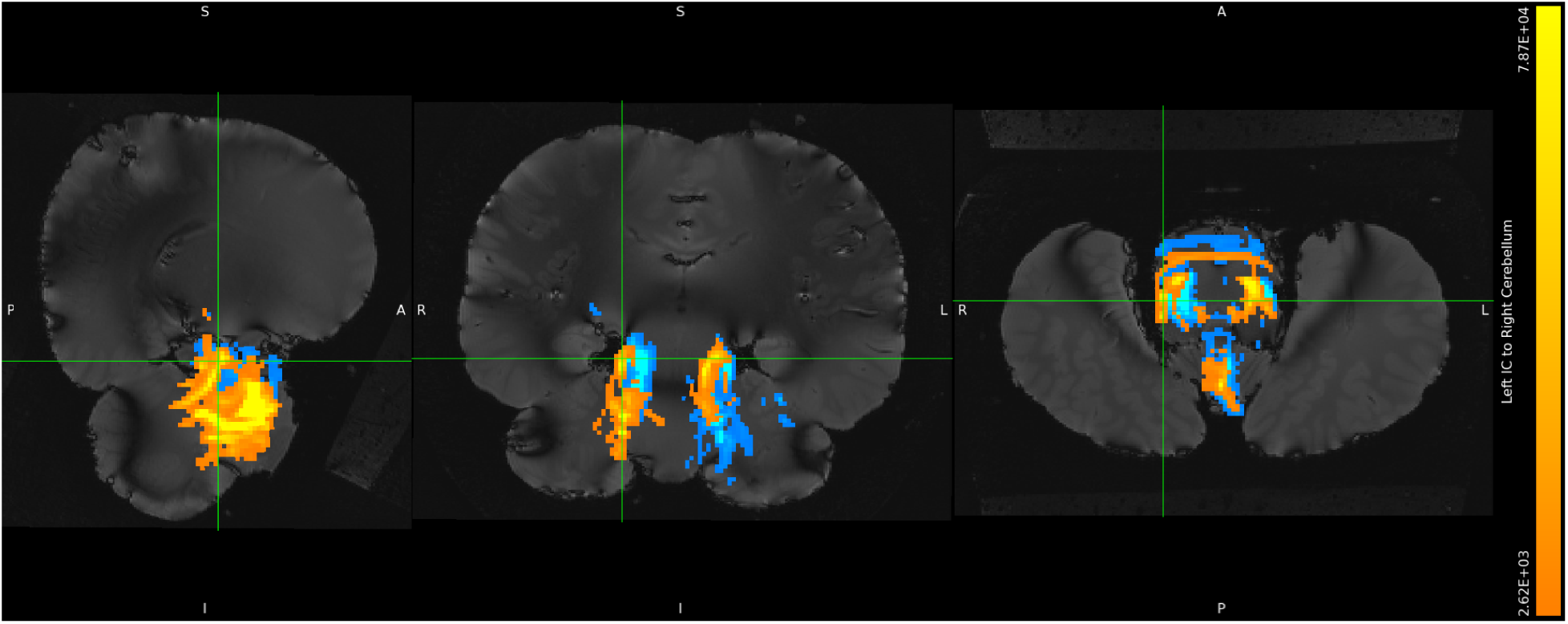
*S. attenuata,* left IC to right cerebellum tracts shown in orange, right IC to left cerebellum tracts shown in turquoise, minimum threshold set to 1% and maximum threshold set to 30% of waytotals. Orthographic view.

**Figure S6B2.**
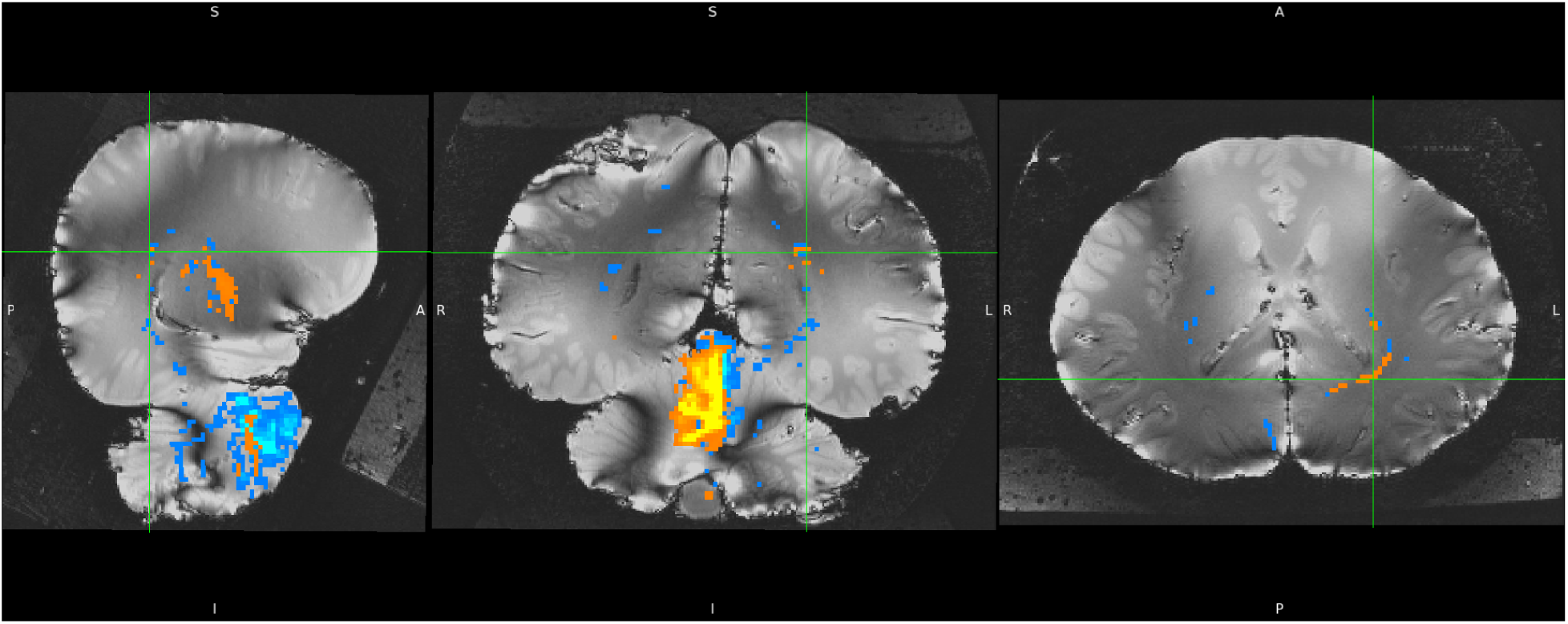
*S. attenuata,* left IC to right cerebellum tracts shown in orange, right IC to left cerebellum tracts shown in turquoise, set to a more liberal threshold of minimum 0.1% and maximum 5% of waytotals. Orthographic view.

**Figure S6B3.**
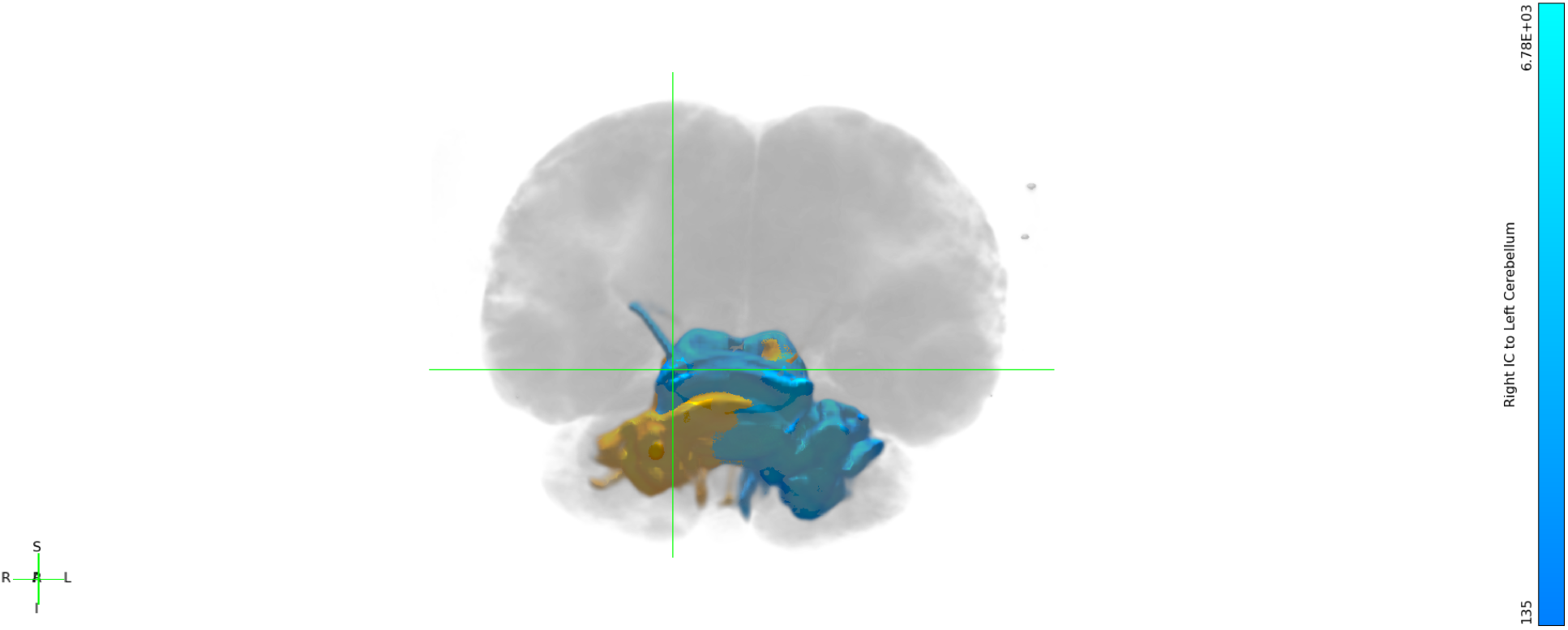
*S. attenuata,* left IC to right cerebellum tracts shown in orange, right IC to left cerebellum tracts shown in turquoise, set to a more liberal threshold of minimum 0.1% and maximum 5% of waytotals. Still 3-dimensional view.

**Figure S6B4.**
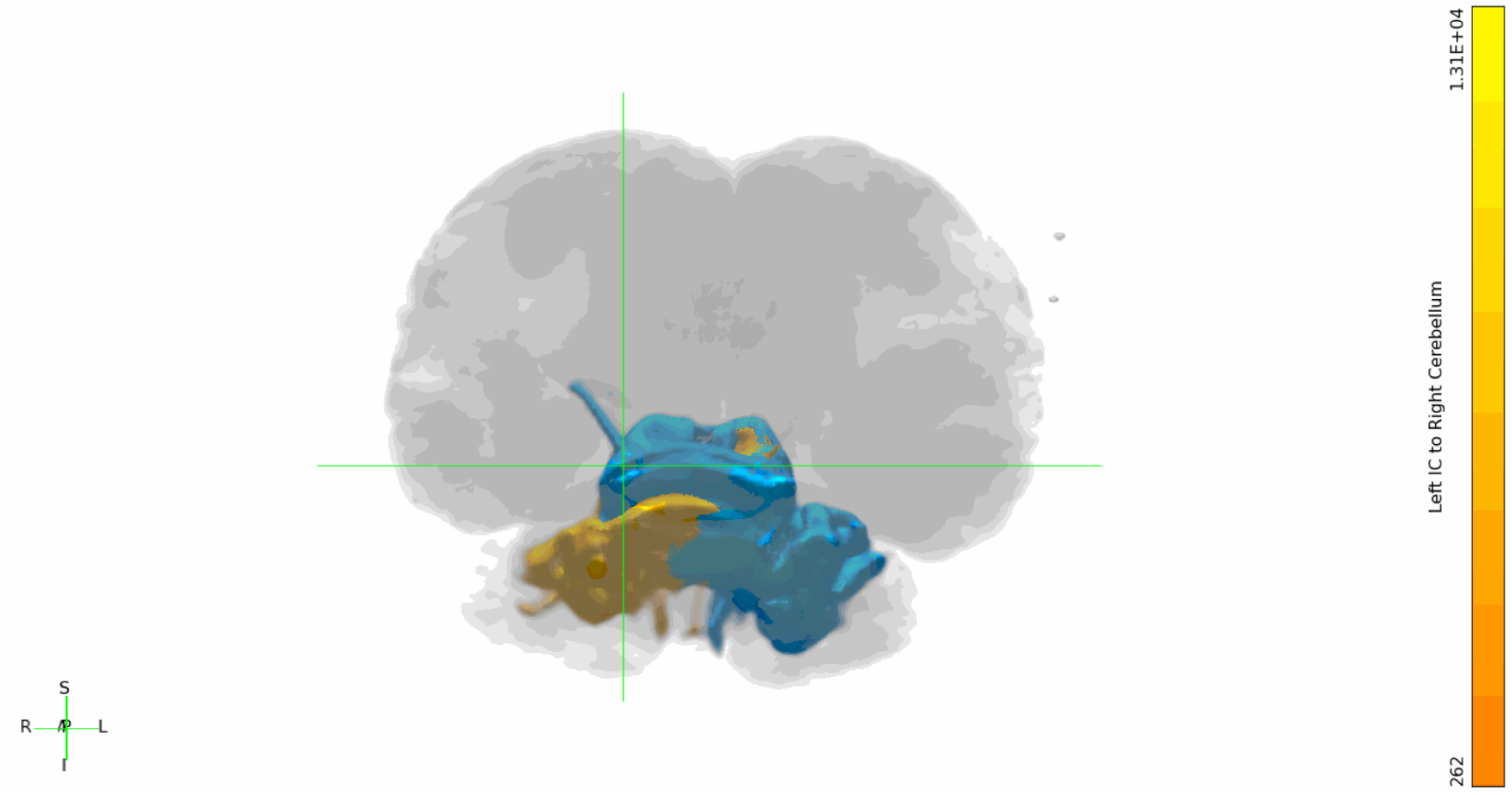
*S. attenuata,* left IC to right cerebellum tracts shown in orange, right IC to left cerebellum tracts shown in turquoise, set to a more liberal threshold of minimum 0.1% and maximum 5% of waytotals. Rotating 3-dimensional view.

**Figure S6C1.**
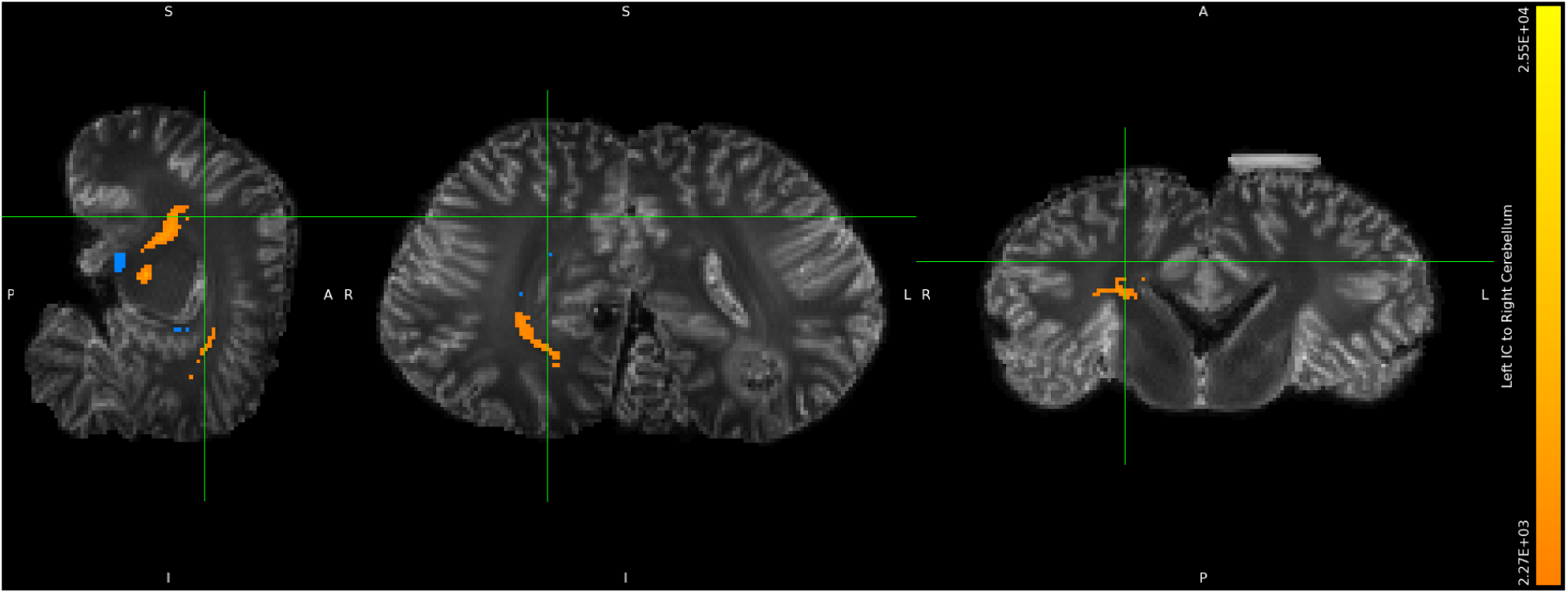
*L. acutus,* left IC to right cerebellum tracts shown in orange, right IC to left cerebellum tracts shown in turquoise, minimum threshold set to 1% and maximum threshold set to 30% of waytotals. Orthographic view.

**Figure S6C2.**
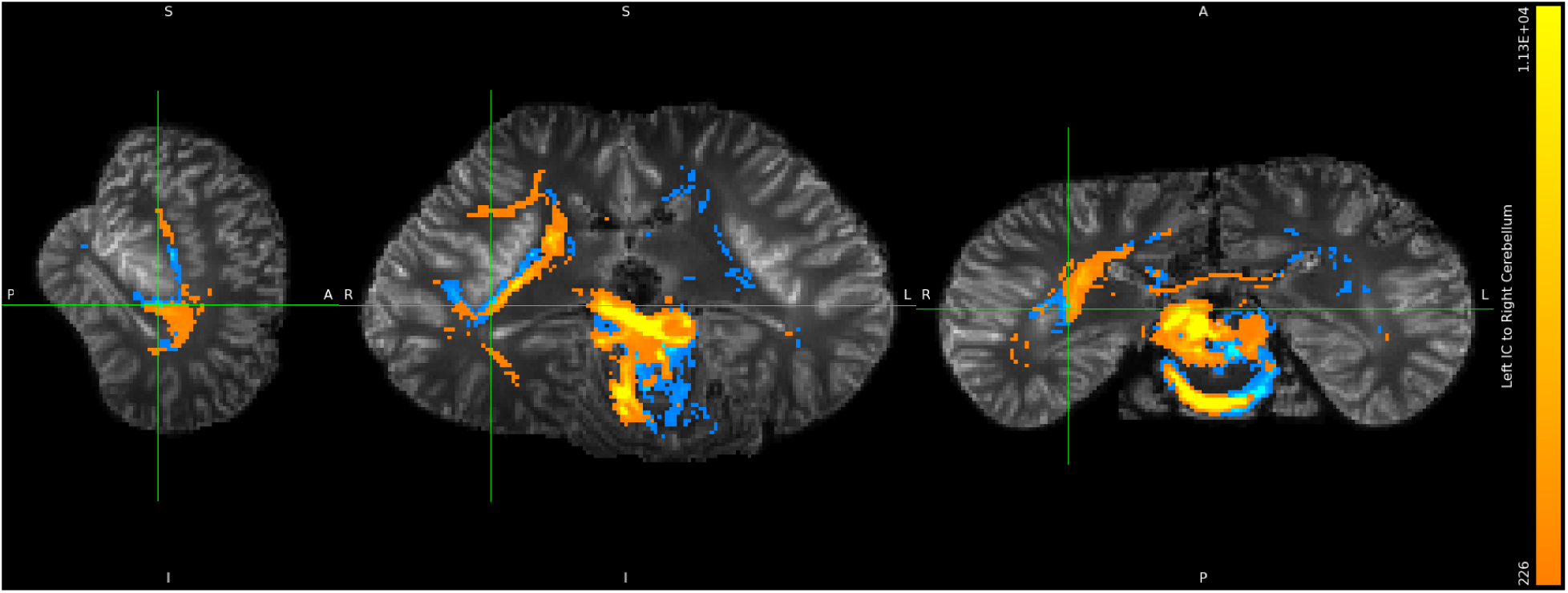
*L. acutus,* left IC to right cerebellum tracts shown in orange, right IC to left cerebellum tracts shown in turquoise, set to a more liberal threshold of minimum 0.1% and maximum 5% of waytotals. Orthographic view.

**Figure S6C3.**
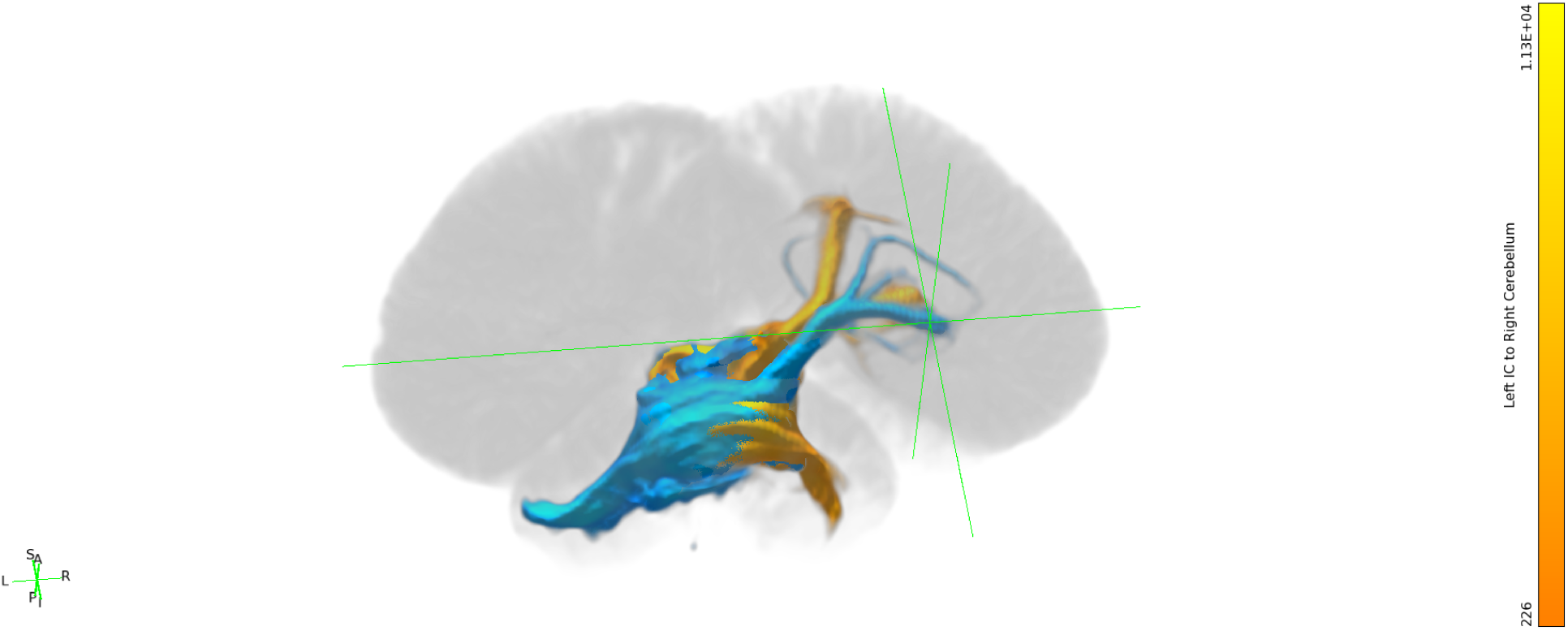
*L. acutus,* left IC to right cerebellum tracts shown in orange, right IC to left cerebellum tracts shown in turquoise, set to a more liberal threshold of minimum 0.1% and maximum 5% of waytotals. Still 3-dimensional view.

**Figure S6C4.**
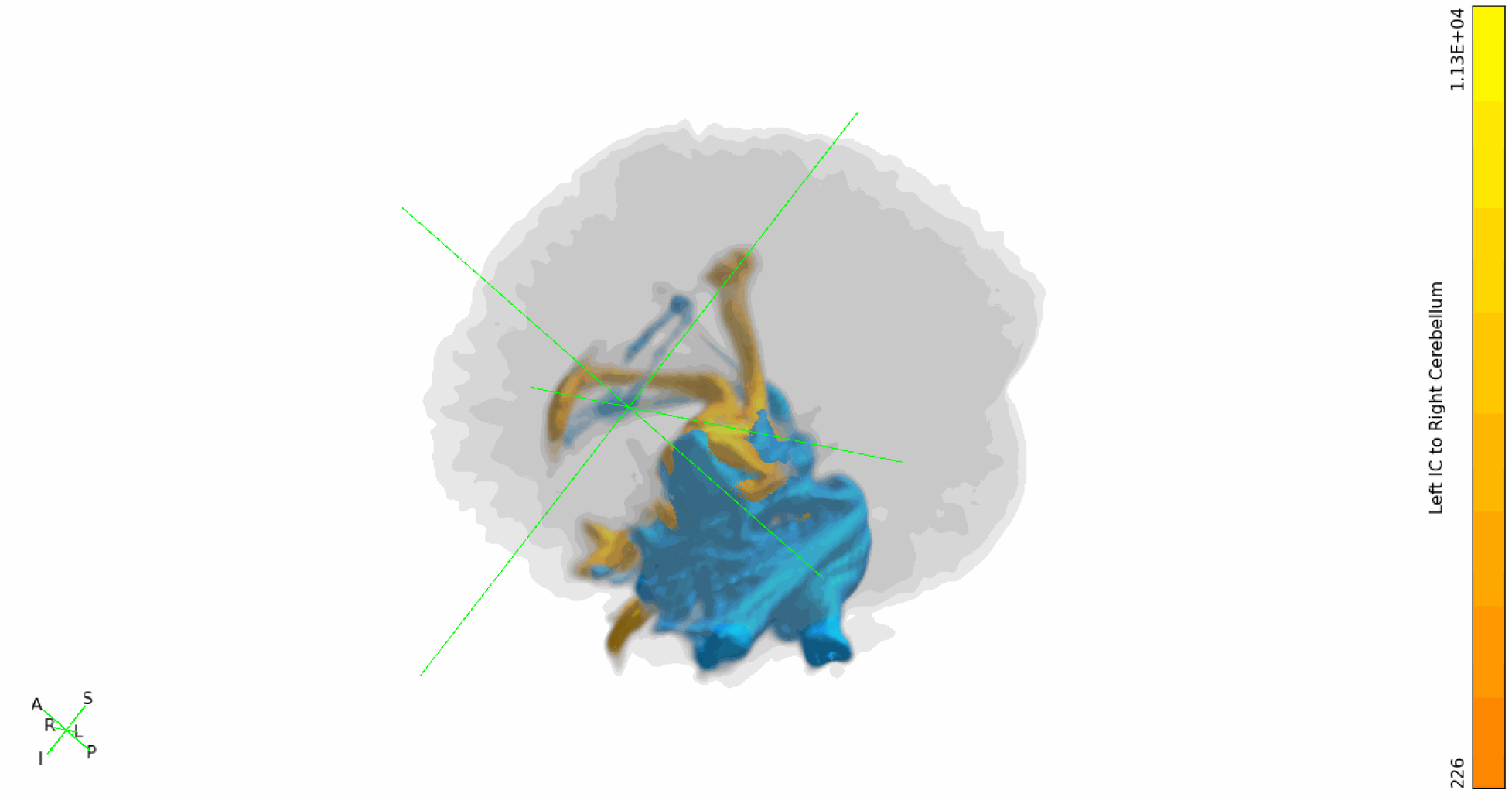
*L. acutus,* left IC to right cerebellum tracts shown in orange, right IC to left cerebellum tracts shown in turquoise, set to a more liberal threshold of minimum 0.1% and maximum 5% of waytotals. Rotating 3-dimensional view.

**Figure S6D1.**
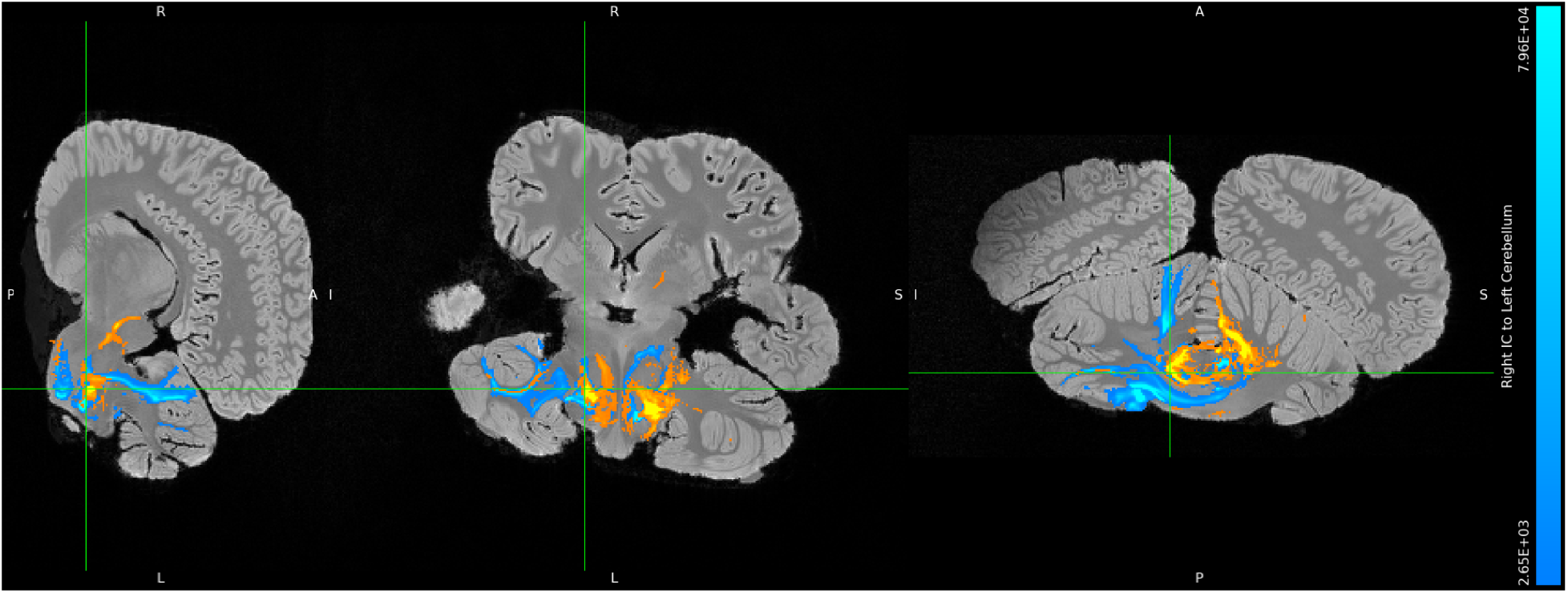
*B. borealis,* left IC to right cerebellum tracts shown in orange, right IC to left cerebellum tracts shown in turquoise, minimum threshold set to 1% and maximum threshold set to 30% of waytotals. Orthographic view.

**Figure S6D2.**
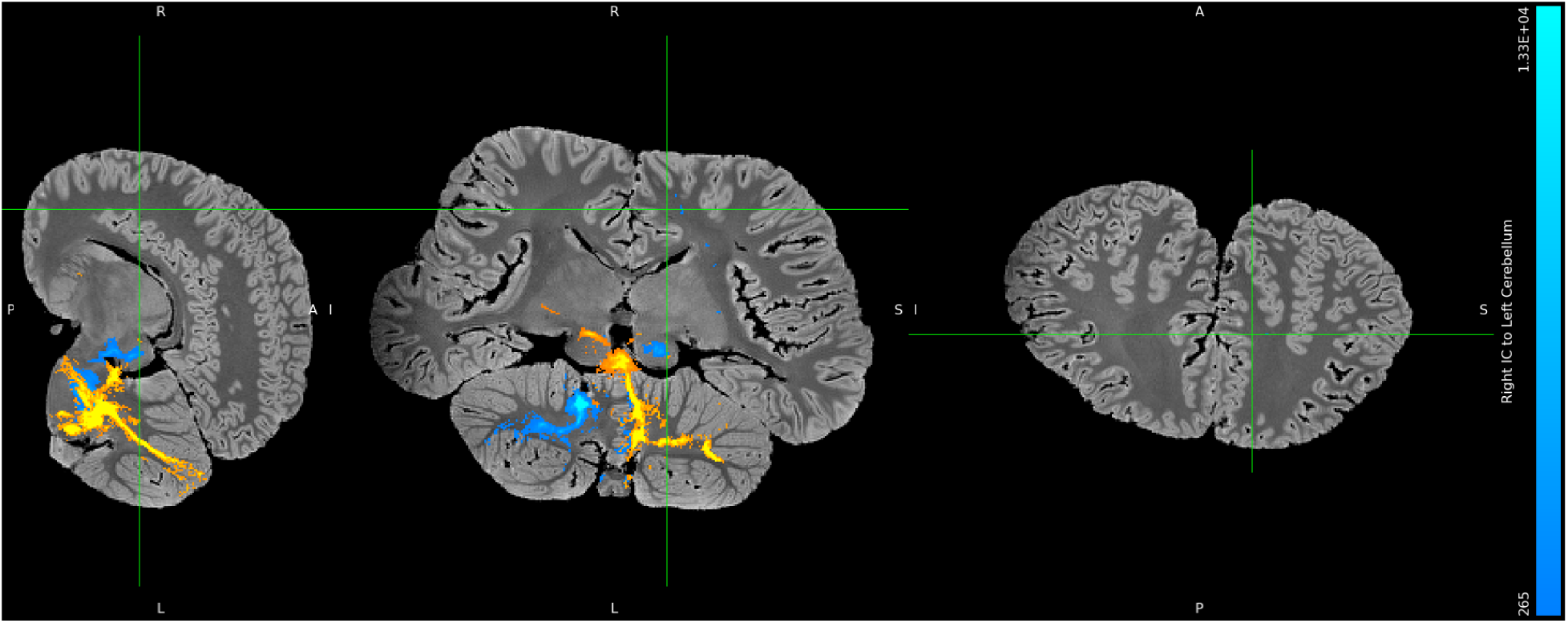
*B. borealis,* left IC to right cerebellum tracts shown in orange, right IC to left cerebellum tracts shown in turquoise, set to a more liberal threshold of minimum 0.1% and maximum 5% of waytotals. Orthographic view.

**Figure S6D3.**
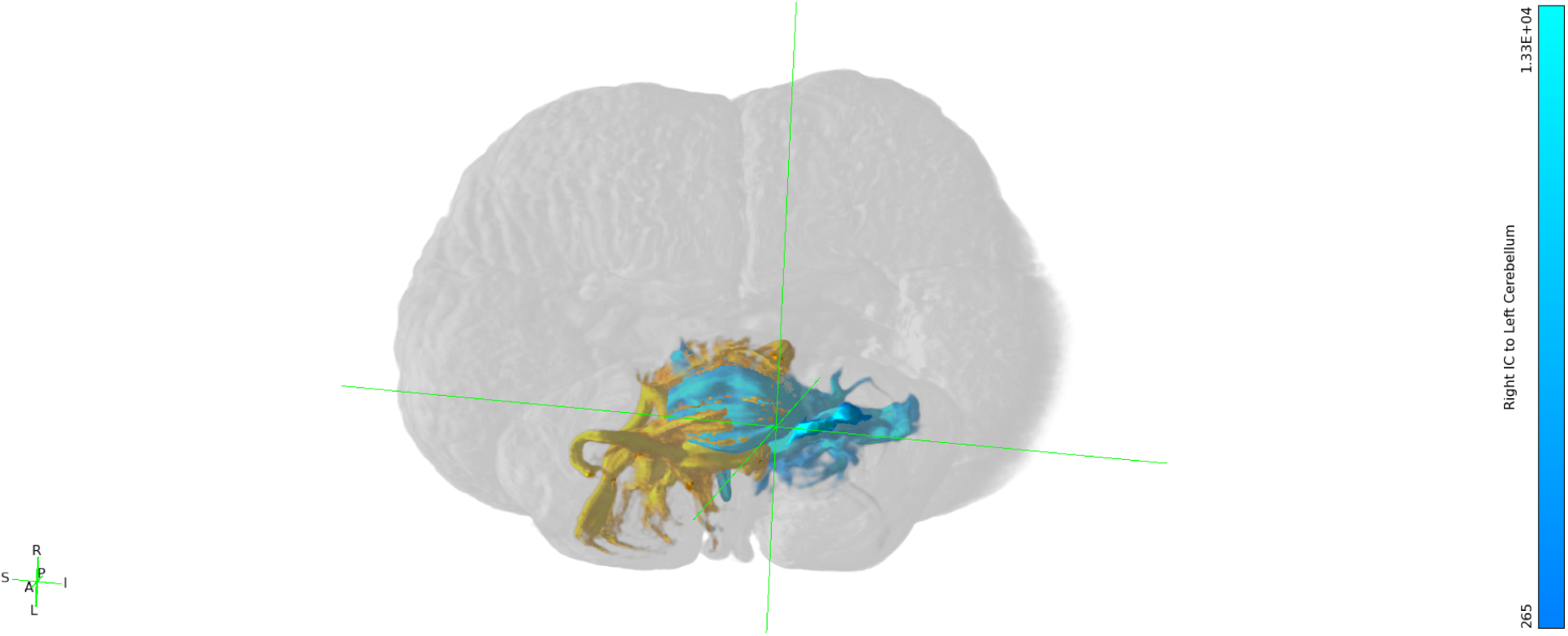
*B. borealis,* left IC to right cerebellum tracts shown in orange, right IC to left cerebellum tracts shown in turquoise, set to a more liberal threshold of minimum 0.1% and maximum 5% of waytotals. Still 3-dimensional view.

**Figure S6D4.**
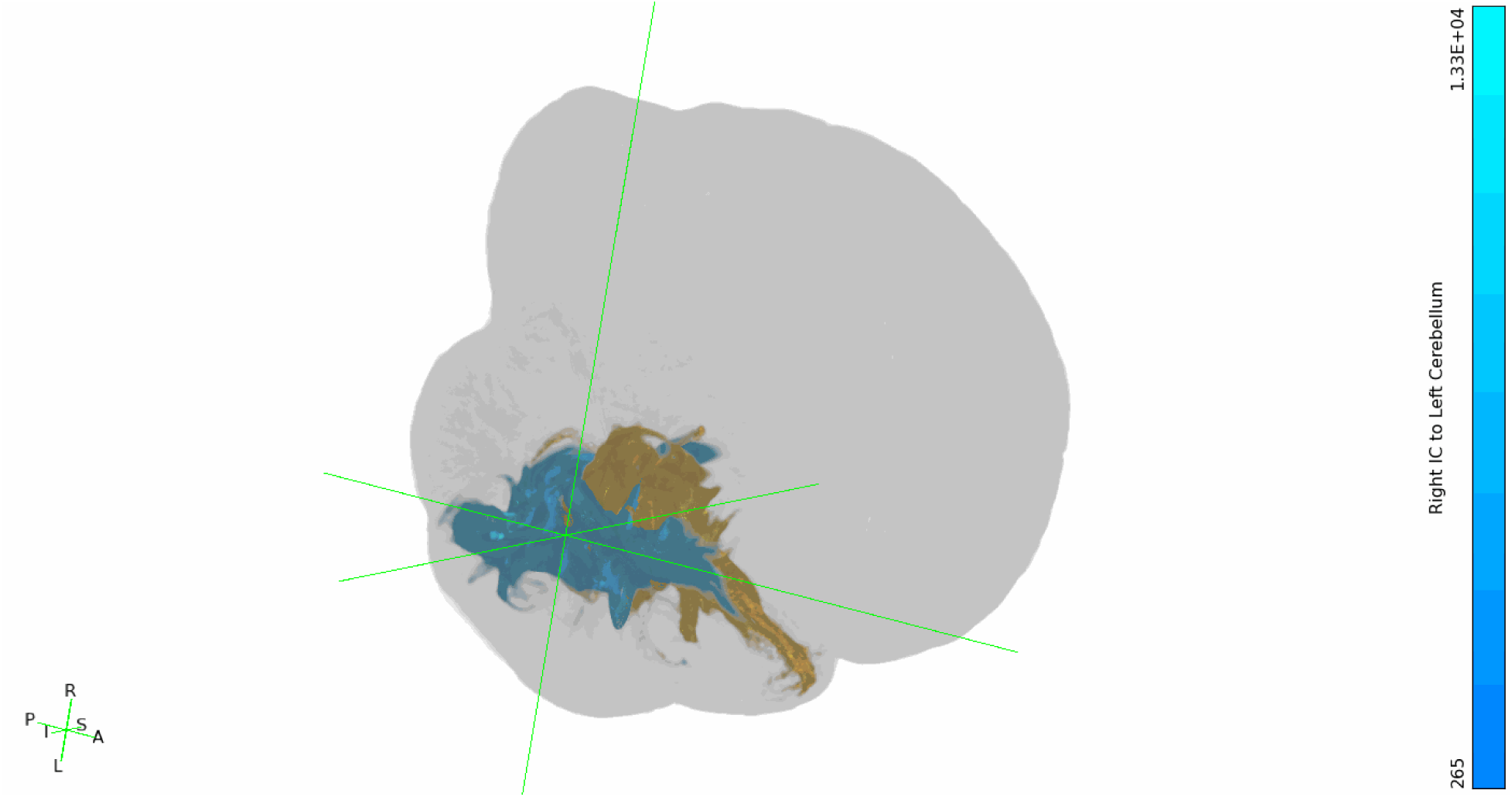
*B. borealis,* left IC to right cerebellum tracts shown in orange, right IC to left cerebellum tracts shown in turquoise, set to a more liberal threshold of minimum 0.1% and maximum 5% of waytotals. Rotating 3-dimensional view.

### S7 Figures. Specific cerebellar targets in IC-cerebellar tractograms

**Figure S7A1.**
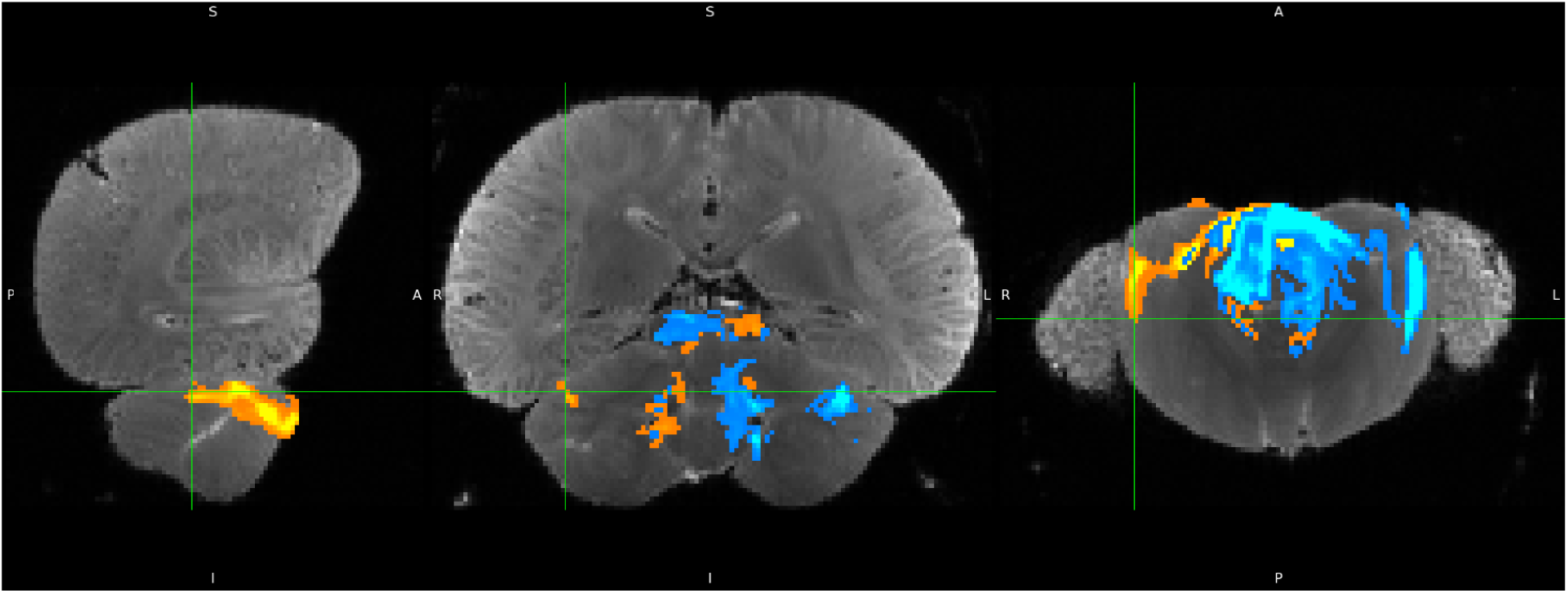
*D. delphis,* left IC to right cerebellum tracts shown in orange, minimum threshold set to 1% and maximum threshold set to 30% of waytotals.

**Figure S7A2.**
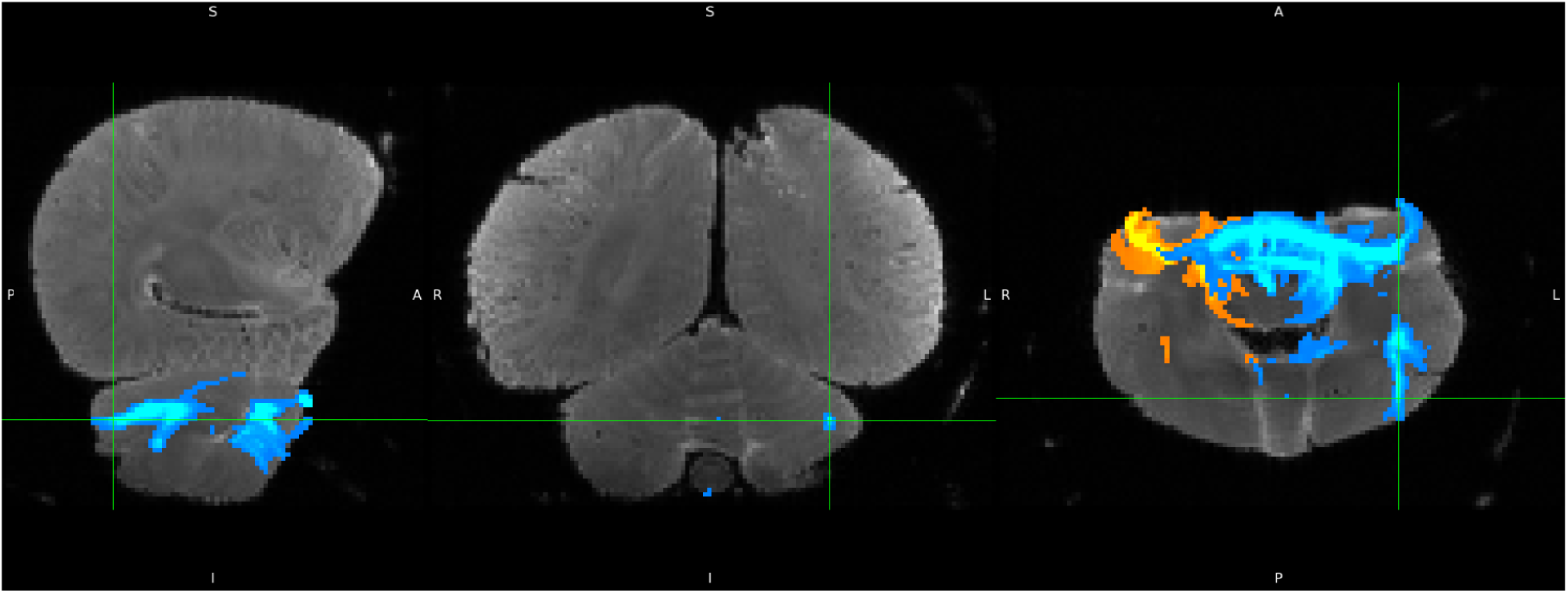
*D. delphis,* right IC to left cerebellum tracts shown in blue, minimum threshold set to 1% and maximum threshold set to 30% of waytotals.

**Figure S7B1.**
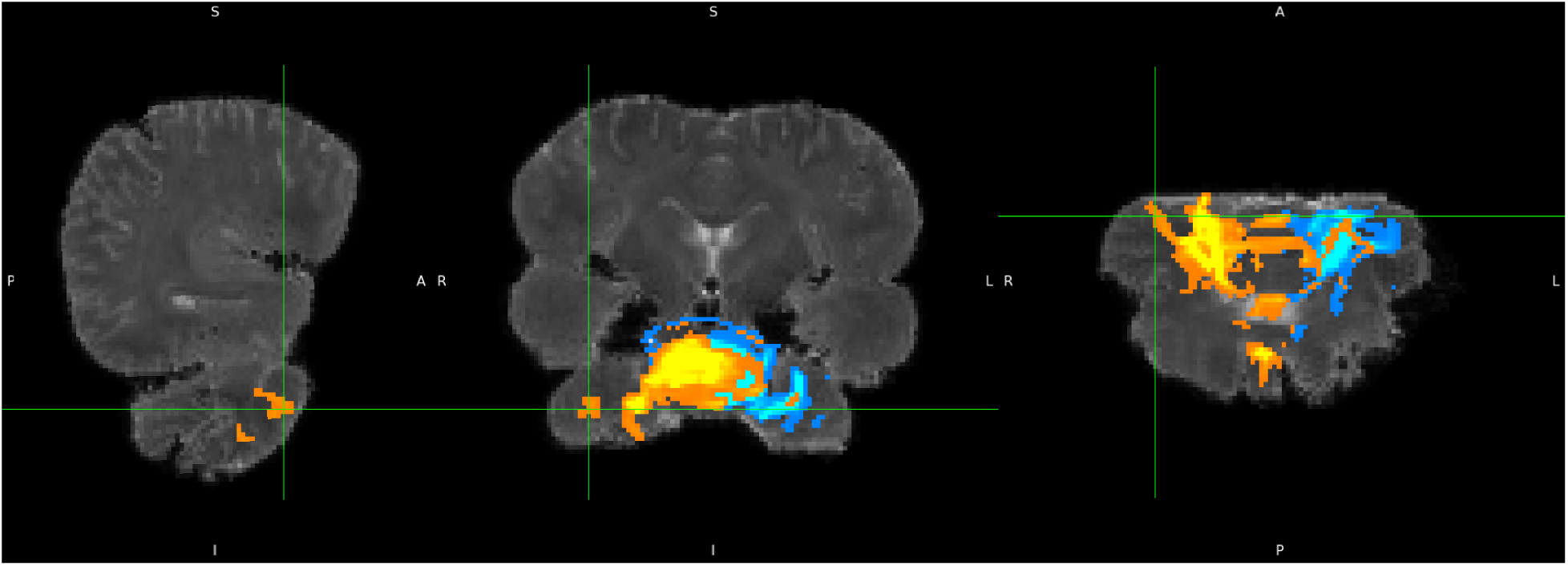
*S. attenuata,* left IC to right cerebellum tracts shown in orange, minimum threshold set to 1% and maximum threshold set to 30% of waytotals.

**Figure S7B2.**
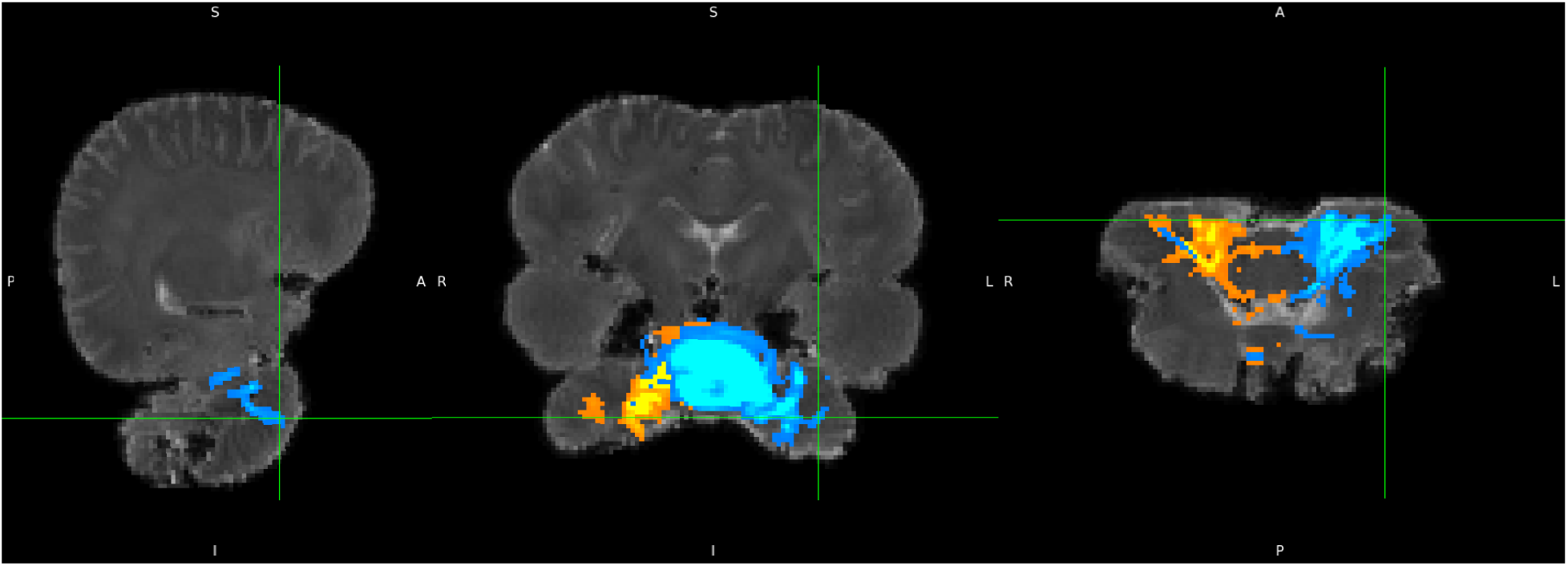
*S. attenuata*, right IC to left cerebellum tracts shown in blue, minimum threshold set to 1% and maximum threshold set to 30% of waytotals.

**Figure S7C1.**
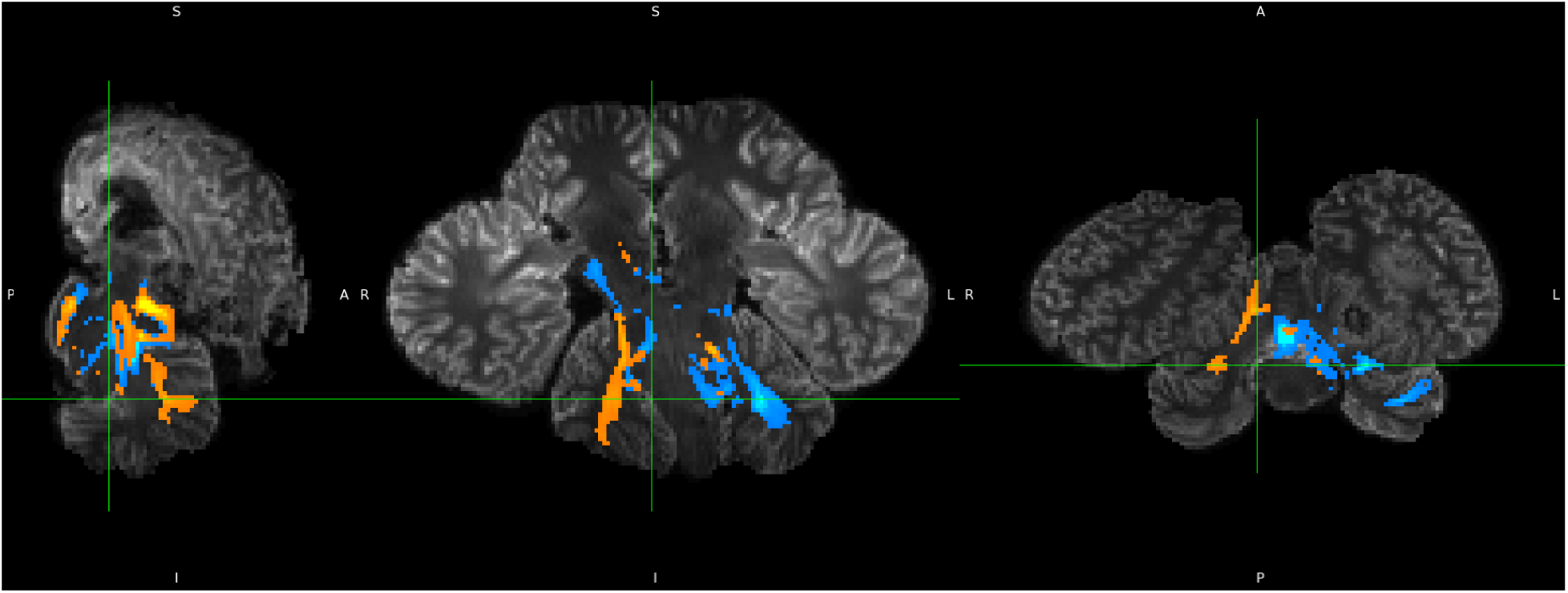
*L. acutus,* left IC to right cerebellum tracts shown in orange, minimum threshold set to 1% and maximum threshold set to 30% of waytotals.

**Figure S7C2.**
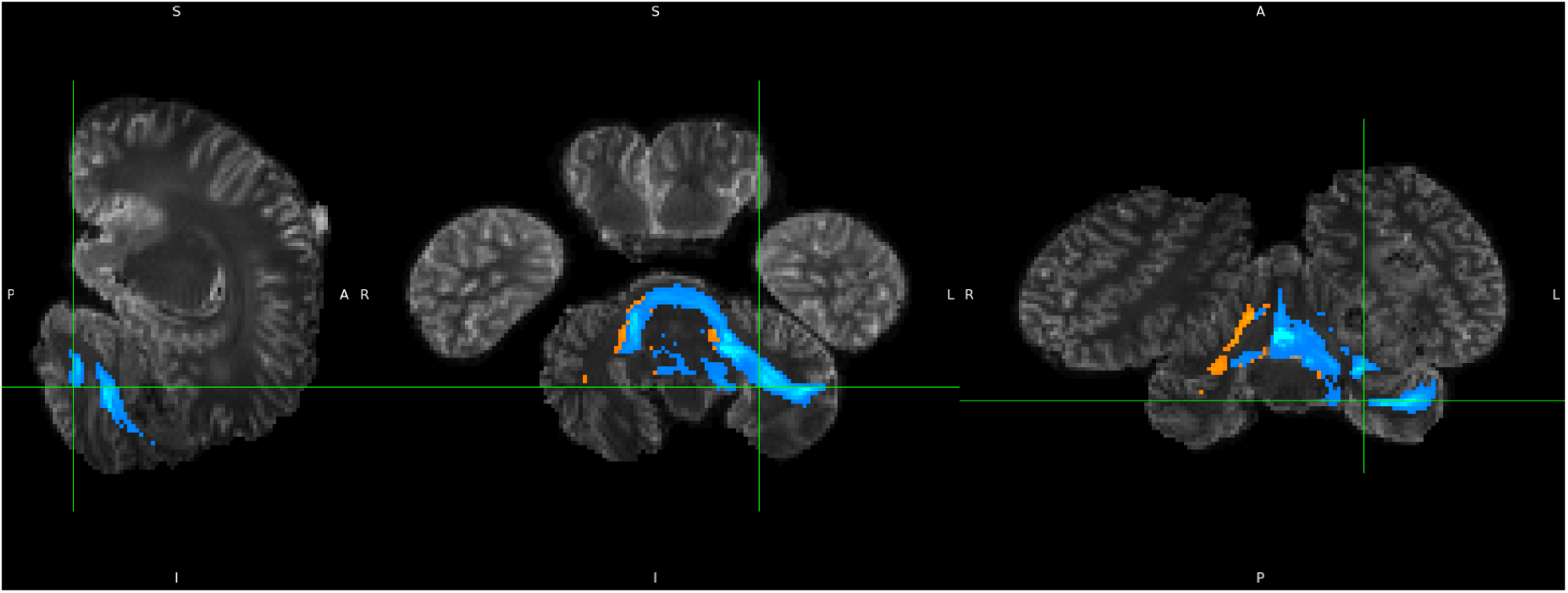
*L. acutus,* right IC to left cerebellum tracts shown in blue, minimum threshold set to 1% and maximum threshold set to 30% of waytotals.

**Figure S7D1.**
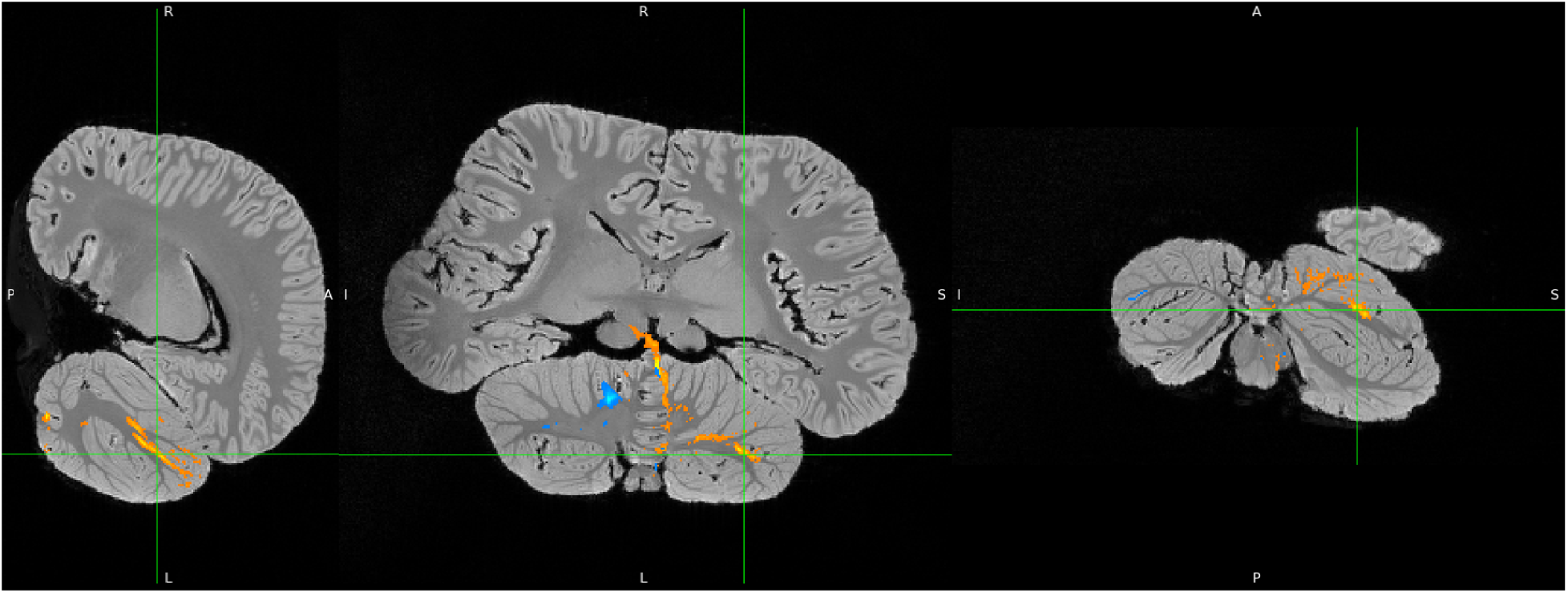
*B. borealis,* left IC to right cerebellum tracts shown in orange, minimum threshold set to 1% and maximum threshold set to 30% of waytotals.

**Figure S7D2.**
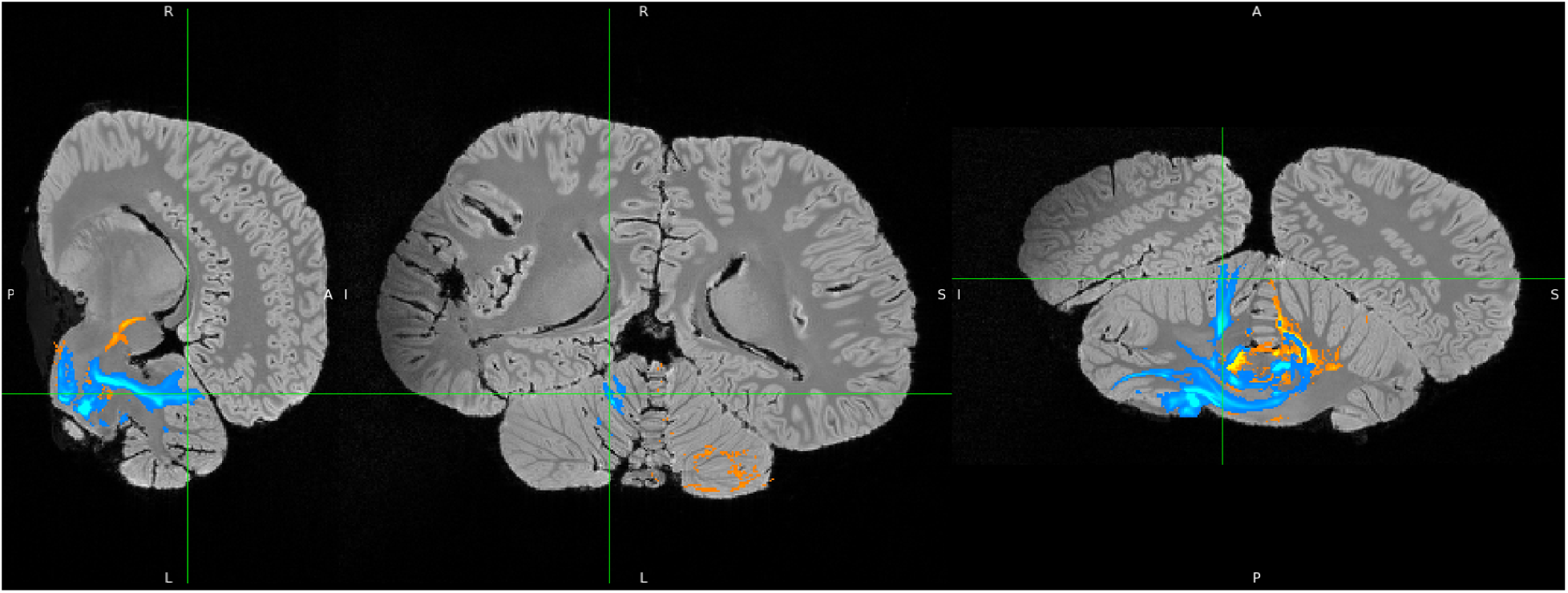
*B. borealis,* right IC to left cerebellum tracts shown in turquoise, minimum threshold set to 1% and maximum threshold set to 30% of waytotals.

### S8 Tables. Results of the systematic tractographical analysis conducted for each specimen

S8 Table 1. Common dolphin.

S8 Table 2. Pantropical spotted dolphin.

S8 Table 3. Atlantic white-sided dolphin.

S8 Table 4. Sei whale.

### S9 Table. Cerebellar projection sites and putative functions

**Table S9.** Display of which cerebellar subregions were targeted in auditory-cerebellar traces, with columns further specifying the species involved, laterality in odontocetes versus mysticete, and putative function as informed by studies in other non-cetacean mammals (Azizi et al., 1985; Brissenden et al., 2018; Burne & Woodward, 1983; Cozzi et al., 2018; Hanson et al., 2013; Suga & Horikawa, 1986; O’Reilly et al., 2009; Kirschen et al., 2010; Klein et al., 2016; Metoki et al., 2022; Nakatani et al., 2022; Van Overwalle et al., 2020; Snider & Stowell, 1944). Asterisks in the “Laterality in Odontoceti” column indicate that the listed laterality was not consistently observed across all specimens, but rather represents the overall lateralization pattern. For example, the asterisk next to the “Left” designation in this column’s “Vermis” row reflects that the evidence for a left-vermis bias in odontocete auditory-cerebellar traces was somewhat mixed, given its left-side targeting in *S. attenuata* and its bilateral targeting in *L. acutus*.

| Subregion: | Implicated in: | Laterality in Odontoceti | Laterality in Mysticete | Function in humans and other mammals |
| --- | --- | --- | --- | --- |
| **I-V** | - | - | - | Somatomotor representation of fore and hind limbs |
| **VI** | *L. acutus*, *B. borealis* | Right | Left | Somatomotor representation of head, neck, lips, tongue |
| **Crus I** | *D. delphis*, *S. attenuata*, *L. acutus*, *B. borealis* | Left | Left | Executive processing, working memory, relational processing, syntactic language comprehension (especially in right lobule) |
| **Crus II** | *D. delphis*, *S. attenuata*, *L. acutus* | Left | - | Executive and associative processing, semantic language comprehension (especially in right lobule), theory of mind, social cognition, and relational processing (especially in left lobule) |
| **VIIb** | *D. delphis*, *B. borealis* | Bilateral | Left | Understudied; *possibly* visuospatial working memory, sensorimotor representation of limbs, or auditory processing |
| **VIIIa** | *D. delphis*, *L. acutus*, *B. borealis* | Right* | Bilateral | Sensorimotor representation, auditory processing in humans and cats, echolocative sound production in bats |
| **VIIIb** | *S. attenuata*, *L. acutus* | Bilateral | - | Sensorimotor representation, auditory processing in humans and cats, echolocative sound production in bats |
| **IX** | *D. delphis*, *S. attenuata*, *L. acutus*, *B. borealis* | Left* | Bilateral | Auditory and visual processing in rodents, echolocative sound production in bats, language and social cognition in humans |
| **X** | *D. delphis* | Bilateral | - | Balance/vestibular processing and eye movement |
| **Vermis** | *S. attenuata*, *L. acutus* | Left* | - | Social and emotional/affective processing |

## Notes

### Competing Interest Statement

The authors have declared no competing interest.

## References

Adrian, E. (1943). Afferent areas in the brain of ungulates. Brain, 66, 89–103. (doi:10.1093/brain/66.2.89)

Ames, A., Beedholm, K., & Madsen, P. T. (2020). Lateralized sound production in the beluga whale (*Delphinapterus leucas*). Journal of Experimental Biology, 223(17). 10.1242/jeb.226316

Andrews, R.J., Knight R.T., Kirby, R.P. (1990). Evoked potential mapping of auditory and somatosensory cortices in the miniature swine. Neuroscience Letters, 114, 27–31. (doi:10.1016/0304-3940(90)90423-7)

Au, W. W. L. (1993). *The sonar of dolphins*. Springer Science & Business Media.

Azizi, S. A., Burne, R. A., & Woodward, D. J. (1985). The auditory corticopontocerebellar projection in the rat: Inputs to the paraflocculus and midvermis. an anatomical and physiological study. Experimental Brain Research, 59(1). 10.1007/bf00237663

Basser, P. J., Pajevic, S., Pierpaoli, C., Duda, J., & Aldroubi, A. (2000). In vivo fiber tractography using DT-MRI data. Magnetic Resonance in Medicine, 44(4), 625–632. 10.1002/1522-2594(200010)44:4%3C625::aid-mrm17%3E3.0.co;2-o

Baumann, O., Borra, R. J., Bower, J. M., Cullen, K. E., Habas, C., Ivry, R. B., Leggio, M., Mattingley, J. B., Molinari, M., Moulton, E. A., Paulin, M. G., Pavlova, M. A., Schmahmann, J. D., & Sokolov, A. A. (2014). Consensus paper: The role of the cerebellum in perceptual processes. The Cerebellum, 14(2), 197–220. 10.1007/s12311-014-0627-7

Baumgartner, M. F., Van Parijs, S. M., Wenzel, F. W., Tremblay, C. J., Carter Esch, H., & Warde, A. M. (2008). Low frequency vocalizations attributed to sei whales (*Balaenoptera borealis*). The Journal of the Acoustical Society of America, 124(2), 1339–1349. 10.1121/1.2945155

Beaulieu C. (2002). The basis of anisotropic water diffusion in the nervous system - A technical review. NMR in Biomedicine, 15(7-8), 435–455. 10.1002/nbm.782

Beedholm, K., Ladegaard, M., Madsen, P. T., & Tyack, P. L. (2023). Latencies of click-evoked auditory responses in a harbor porpoise exceed the time interval between subsequent echolocation clicks. The Journal of the Acoustical Society of America, 153(2), 952–960. 10.1121/10.0017163

Berns, G. S., Cook, P. F., Foxley, S., Jbabdi, S., Miller, K. L., & Marino, L. (2015). Diffusion tensor imaging of dolphin brains reveals direct auditory pathway to temporal lobe. Proceedings of the Royal Society B: Biological Sciences, 282(1811). 10.1098/rspb.2015.1203

Branstetter, B. K., Finneran, J. J., Fletcher, E. A., Weisman, B. C., & Ridgway, S. H. (2012). Dolphins can maintain vigilant behavior through echolocation for 15 days without interruption or cognitive impairment. PLoS ONE, 7(10), e47478. 10.1371/journal.pone.0047478

Brissenden, J. A., Tobyne, S. M., Osher, D., Levin, E. E., Halko, M. A., & Somers, D. C. (2018). Topographic cortico-cerebellar networks revealed by visual attention and working memory. Current Biology, 28(21), 3364–3372.e5. 10.1016/j.cub.2018.08.059

Bullock, T.H., & Ridgway, S.H (1972). Evoked potentials in the central auditory system of alert porpoises to their own and artificial sounds. Journal of Neurobiology, 3, 79–99.

Bullock, T. H., & Gurevich, V. S. (1979). Soviet literature on the nervous system and psychobiology of cetacea. International Review of Neurobiology, 21, 47–127. 10.1016/s0074-7742(08)60637-6

Burne, R. A., & Woodward, D. J. (1983). Visual cortical projections to the paraflocculus in the rat. Experimental Brain Research, 49(1). 10.1007/bf00235541

Buxton, R. B. (1993). The diffusion sensitivity of fast steady-state free precession imaging. Magnetic Resonance in Medicine, 29(2), 235–243. 10.1002/mrm.1910290212

Canning, C., Crain, D., Eaton, T. S., Nuessly, K., Friedlaender, A., Hurst, T., … & Weinrich, M. (2011). Population-level lateralized feeding behaviour in North Atlantic humpback whales, *Megaptera novaeangliae*. Animal Behaviour, 82(4), 901–909.

Clark, C., Ellison, W., Southall, B., Hatch, L., Van Parijs, S., Frankel, A., & Ponirakis, D. (2009). Acoustic masking in marine ecosystems: Intuitions, analysis, and implication. Marine Ecology Progress Series, 395, 201–222. 10.3354/meps08402

Cohen-Gadol, A. (2024). *The Neurosurgical Atlas*. The Neurosurgical Atlas. https://www.neurosurgicalatlas.com/

Cook, P. F., Berns, G. S., Colegrove, K., Johnson, S., & Gulland, F. (2018). Postmortem DTI reveals altered hippocampal connectivity in wild sea lions diagnosed with chronic toxicosis from algal exposure. Journal of Comparative Neurology, 526(2), 216–228.

Cook, P. F., Huggenberger, S., & Cozzi, B. (2024). Neurophysiology. In The Physiology of Dolphins (pp. 163–191). Academic Press.

Committee on Taxonomy. (2023). List of marine mammal species and subspecies. Society for Marine Mammalogy, www.marinemammalscience.org, consulted on Nov. 5, 2023.

Coombs, E. J., Clavel, J., Park, T., Churchill, M., & Goswami, A. (2020). Wonky whales: The evolution of cranial asymmetry in cetaceans. BMC Biology, 18(1). 10.1186/s12915-020-00805-4

Coombs, E. J., Felice, R. N., Clavel, J., Park, T., Bennion, R. F., Churchill, M., Geisler, J. H., Beatty, B., & Goswami, A. (2022). The tempo of cetacean cranial evolution. Current Biology, 32(10). 10.1016/j.cub.2022.04.060

Covey, E. (2005). Neurobiological specializations in echolocating bats. *The Anatomical Record Part A: Discoveries in Molecular*, Cellular, and Evolutionary Biology: An Official Publication of the American Association of Anatomists, 287(1), 1103–1116.

Cozzi, B., Huggenberger, S., & Oelschläger, H. A. (2018). Atlas of the anatomy of dolphins and whales. Academic Press.

Cranford, T. W., Amundin, M., & Norris, K. S. (1996). Functional morphology and homology in the odontocete nasal complex: Implications for sound generation. Journal of Morphology, 228(3), 223–285. 10.1002/(sici)1097-4687(199606)228:3%3C223::aid-jmor1%3E3.0.co;2-3

De Vreese, S., Orekhova, K., Morell, M., Gerussi, T., & Graïc, J.-M. (2024). Neuroanatomy of the cetacean sensory systems. Animals, 14(1), 66. 10.3390/ani14010066

Ebner, T. J., & Pasalar, S. (2008). Cerebellum predicts the future motor state. The Cerebellum, 7, 583–588.

Elemans, C. P., Jiang, W., Jensen, M. H., Pichler, H., Mussman, B. R., Nattestad, J., … & Fitch, W. T. (2024). Evolutionary novelties underlie sound production in baleen whales. Nature, 1–7.

Frisina, R. D. (2001). Subcortical neural coding mechanisms for auditory temporal processing. Hearing Research, 158(1-2), 1–27.

Fox, K. C. R., Muthukrishna, M., & Shultz, S. (2017). The social and cultural roots of whale and dolphin brains. Nature Ecology & Evolution, 1(11), 1699–1705. 10.1038/s41559-017-0336-y

Gatesy, J., Geisler, J. H., Chang, J., Buell, C., Berta, A., Meredith, R. W., … & McGowen, M. R. (2013). A phylogenetic blueprint for a modern whale. Molecular Phylogenetics and Evolution, 66(2), 479–506.

Geraci, J. R., & Lounsbury, V. J. (2005). Marine mammals ashore: A field guide for strandings. National Aquarium in Baltimore.

Gerussi, T., Graïc, J. M., Peruffo, A., Behroozi, M., Schlaffke, L., Huggenberger, S., Güntürkün, O., & Cozzi, B. (2023). The prefrontal cortex of the bottlenose dolphin (Tursiops truncatus Montagu, 1821): a tractography study and comparison with the human. Brain Structure & Function, 228(8), 1963–1976. 10.1007/s00429-023-02699-8

Graïc, J. M., Corain, L., Finos, L., Vadori, V., Grisan, E., Gerussi, T., Orekhova, K., Centelleghe, C., Cozzi, B., & Peruffo, A. (2024). Age-related changes in the primary auditory cortex of newborn, adults and aging bottlenose dolphins (*Tursiops truncatus*) are located in the upper cortical layers. Frontiers in Neuroanatomy, 17. 10.3389/fnana.2023.1330384

Hanson, A. M., Grisham, W., Sheh, C., Annese, J., & Ridgway, S. H. (2013). Quantitative examination of the bottlenose dolphin cerebellum. The Anatomical Record, 296(8), 1215–1228. 10.1002/ar.22726

Harper, C. J., McLellan, W. A., Rommel, S. A., Gay, D. M., Dillaman, R. M., & Pabst, D. A. (2008). Morphology of the melon and its tendinous connections to the facial muscles in bottlenose dolphins (*Tursiops truncatus*). Journal of Morphology, 269(7), 820–839. 10.1002/jmor.10628

Hermundstad, A. M., Bassett, D. S., Brown, K., Aminoff, E., Clewett, D., Freeman, S., Frithsen, A., Johnson, A., Tipper, C. M., Miller, M. B., Grafton, S. T., & Carlson, J. M. (2013). Structural foundations of resting-state and task-based functional connectivity in the human brain. Proceedings of the National Academy of Sciences of the United States of America, 110(15), 6169–6174. 10.1073/pnas.1219562110

Herzing, D. L., & Johnson, C. M. (2015). *Making sense of it all: Multimodal dolphin communication*. Dolphin Communication and Cognition: Past, Present, and Future, 139–171.

Hof, P. R., Chanis, R., & Marino, L. (2005). Cortical complexity in cetacean brains. *The Anatomical Record Part A: Discoveries in Molecular*, Cellular, and Evolutionary Biology, 287A(1), 1142–1152. 10.1002/ar.a.20258

Huffman, R. F., & Henson Jr, O. W. (1990). The descending auditory pathway and acousticomotor systems: Connections with the inferior colliculus. Brain Research Reviews, 15(3), 295–323.

Janik, V., & Slater, P. J. B. (1997). Vocal learning in mammals. Advances in the Study of Behavior, 26, 59–99. 10.1016/S0065-3454(08)60377-0

Janik, V. M., Sayigh, L. S., & Wells, R. S. (2006). Signature whistle shape conveys identity information to bottlenose dolphins. Proceedings of the National Academy of Sciences, 103(21), 8293–8297. 10.1073/pnas.0509918103

Janik, V. M. (2014). Cetacean vocal learning and communication. Current Opinion in Neurobiology, 28, 60–65.

Janik, V. M., & Knörnschild, M. (2021). Vocal production learning in mammals revisited. Philosophical Transactions of the Royal Society B, 376(1836), 20200244.

Janvier, P. (2002). Early vertebrates. Clarendon Press, Oxford University Press. (Original work published 1996).

Jbabdi, S., Behrens, T. E., & Smith, S. M. (2010). Crossing fibres in tract-based spatial statistics. Neuroimage, 49(1), 249–256.

Jen, P. H. S., & Schlegel, P. A. (1980). Neurons in the cerebellum of echolocating bats respond to acoustic signals. Brain Research, 196(2), 502–507.

Kamada, T., & Jen, P. H. S. (1990). Auditory response properties and directional sensitivity of cerebellar neurons of the echolocating bat, *Eptesicus fuscus*. Brain Research, 528(1), 123–129.

Kanwal, J. S. (2012). Right–left asymmetry in the cortical processing of sounds for social communication vs. navigation in mustached bats. European Journal of Neuroscience, 35(2), 257–270.

Karenina, K., Giljov, A., Ivkovich, T., & Malashichev, Y. (2016). Evidence for the perceptual origin of right-sided feeding biases in cetaceans. Animal Cognition, 19, 239–243.

Kirschen, M. P., Chen, S. H. A., & Desmond, J. E. (2010). Modality specific cerebro-cerebellar activations in verbal working memory: An fMRI study. Behavioural Neurology, 23(1-2), 51–63. 10.3233/BEN-2010-0266

Klein, A. P., Ulmer, J. L., Quinet, S. A., Mathews, V., & Mark, L. P. (2016). Nonmotor functions of the Cerebellum: An introduction. American Journal of Neuroradiology, 37(6), 1005–1009. 10.3174/ajnr.a4720

Kremers, D., Célérier, A., Schaal, B., Campagna, S., Trabalon, M., Böye, M., Hausberger, M., & Lemasson, A. (2016). Sensory perception in cetaceans: Part I—current knowledge about dolphin senses as a representative species. Frontiers in Ecology and Evolution, 4. 10.3389/fevo.2016.00049

Knolle, F., Schröger, E., Baess, P., & Kotz, S. A. (2012). The cerebellum generates motor-to-auditory predictions: ERP lesion evidence. Journal of Cognitive Neuroscience, 24(3), 698–706.

Kössl, M., Hechavarria, J., Voss, C., Macias, S., Mora, E., & Vater, M. (2014). Neural maps for target range in the auditory cortex of echolocating bats. Current Opinion in Neurobiology, 24, 68–75. 10.1016/j.conb.2013.08.016

Ladygina, T. F., & Supin, A. Y. (1977). Localization of the sensory projection areas in the cerebral cortex of the dolphin, Tursiops truncatus. PubMed, 13(6), 712–718.

Ladygina, T. F., Mass, A. M., & Supin, A. I. (1978). Multiple sensory projections in the dolphin cerebral cortex. Zhurnal Vysshei Nervnoi Deiatelnosti Imeni I P Pavlova, 28(5), 1047–1053. https://pubmed.ncbi.nlm.nih.gov/716593/

Laeta, M., Oliveira, J. A., Siciliano, S., Lambert, O., Jensen, F. H., & Galatius, A. (2023). Cranial asymmetry in odontocetes: A facilitator of sonic exploration? Zoology, 160, 126108–126108. 10.1016/j.zool.2023.126108

Lazari, A., & Lipp, I. (2021). Can MRI measure myelin? Systematic review, qualitative assessment, and meta-analysis of studies validating microstructural imaging with myelin histology. NeuroImage, 230, 117744. 10.1016/j.neuroimage.2021.117744

Leliveld, L. M. (2019). From science to practice: A review of laterality research on ungulate livestock. Symmetry, 11(9), 1157.

Lyamin, O. I., Mukhametov, L. M., Siegel, J. M., Nazarenko, E. A., Polyakova, I. N., & Шпак, О. В. (2002). Unihemispheric slow wave sleep and the state of the eyes in a white whale. Behavioural Brain Research, 129(1-2), 125–129. 10.1016/s0166-4328(01)00346-1

Lyamin, O. I., Manger, P. R., Ridgway, S. H., Mukhametov, L. M., & Siegel, J. M. (2008). Cetacean sleep: An unusual form of mammalian sleep. Neuroscience & Biobehavioral Reviews, 32(8), 1451–1484.

Madsen, P. T., Lammers, M., Wisniewska, D., & Beedholm, K. (2013). Nasal sound production in echolocating delphinids (*Tursiops truncatus* and *Pseudorca crassidens*) is dynamic, but unilateral: Clicking on the right side and whistling on the left side. Journal of Experimental Biology, 216(21), 4091–4102. 10.1242/jeb.091306

Malkemper, E. P., Oelschläger, H. H. A., & Huggenberger, S. (2012). The dolphin cochlear nucleus: Topography, histology and functional implications. Journal of Morphology, 273(2), 173–185.

Malmierca, M. S. (2004). The inferior colliculus: A center for convergence of ascending and descending auditory information. Neuroembryology and Aging, 3(4), 215–229

Marino, L., Rilling, J. K., Lin, S. K., & Ridgway, S. H. (2000). Relative volume of the cerebellum in dolphins and comparison with anthropoid primates. Brain Behavior and Evolution, 56(4), 204–211.

Marino, L., Sudheimer, K. D., Pabst, D. A., Mclellan, W. A., Filsoof, D., & Johnson, J. I. (2002). Neuroanatomy of the common dolphin (*Delphinus delphis*) as revealed by magnetic resonance imaging (MRI). The Anatomical Record, 268(4), 411–429. 10.1002/ar.10181

Marino, L., Connor, R. C., Fordyce, R. E., Herman, L. M., Hof, P. R., Lefebvre, L., Lusseau, D., McCowan, B., Nimchinsky, E. A., Pack, A. A., Rendell, L., Reidenberg, J. S., Reiss, D., Uhen, M. D., Van der Gucht, E., & Whitehead, H. (2007). Cetaceans have complex brains for complex cognition. PLoS Biology, 5(5), e139. 10.1371/journal.pbio.0050139

Marx, F. G., & Fordyce, R. E. (2015). Baleen boom and bust: A synthesis of mysticete phylogeny, diversity and disparity. Royal Society Open Science, 2(4), 140434. 10.1098/rsos.140434

Marx, F., Lambert, O., & Uhen, M. D. (2016). Cetacean paleobiology. Wiley-Blackwell.

McLachlan, N. M., & Wilson, S. J. (2017). The contribution of brainstem and cerebellar pathways to auditory recognition. Frontiers in Psychology, 8, 265.

McKenna, M. F., Cranford, T. W., Berta, A., & Pyenson, N. D. (2011). Morphology of the odontocete melon and its implications for acoustic function. Marine Mammal Science, 28(4), 690–713. 10.1111/j.1748-7692.2011.00526.x

McNab, J. A., Jbabdi, S., Deoni, S. C. L., Douaud, G., Behrens, T. E. J., & Miller, K. L. (2009). High resolution diffusion-weighted imaging in fixed human brain using diffusion-weighted steady state free precession. NeuroImage, 46(3), 775–785. 10.1016/j.neuroimage.2009.01.008

Mennink, L. M., van Dijk, J. Marc C., & van Dijk, P. (2020). The cerebellar (para)flocculus: A review on its auditory function and a possible role in tinnitus. Hearing Research, 398, 108081. 10.1016/j.heares.2020.108081

Metoki, A., Wang, Y., & Olson, I. R. (2022). The social cerebellum: A large-scale investigation of functional and structural specificity and connectivity. Cerebral Cortex, 32(5), 987–1003. 10.1093/cercor/bhab260

Miller, K. L., McNab, J. A., Jbabdi, S., & Douaud, G. (2012). Diffusion tractography of post-mortem human brains: optimization and comparison of spin echo and steady-state free precession techniques. NeuroImage, 59(3), 2284–2297.

Montie, E. W., Schneider, G. E., Ketten, D. R., Marino, L., Touhey, K. E., & Hahn, M. E. (2008). Volumetric neuroimaging of the Atlantic white-sided dolphin (*Lagenorhynchus acutus*) brain from in situ magnetic resonance images. The Anatomical Record, 291, 263–282.

Montie, E. W., Wheeler, E., Pussini, N., Battey, T. W. K., Barakos, J., Dennison, S., Colegrove, K., Gulland, F. (2010). Magnetic resonance imaging quality and volumes of brain structures from live and postmortem imaging of California sea lions with clinical signs of domoic acid toxicosis. Diseases of Aquatic Organisms, 91, 243–256.

Morgane, P. J., Jacobs, M. S., & McFarland, W. L. (1980). The anatomy of the brain of the bottlenose dolphin (*Tursiops truncatus*). Surface configurations of the telencephalon of the bottlenose dolphin with comparative anatomical observations in four other cetacean species. Brain Research Bulletin, 5(3), 1–107. 10.1016/0361-9230(80)90272-5

Morgane, P. J., Glezer, I. I., & Jacobs, M. S. (1988). Visual cortex of the dolphin: An image analysis study. Journal of Comparative Neurology, 273(1), 3–25. 10.1002/cne.902730103

Moss, C. F., Ortiz, S. T., & Wahlberg, M. (2023). Adaptive echolocation behavior of bats and toothed whales in dynamic soundscapes. Journal of Experimental Biology, 226(9). 10.1242/jeb.245450

Murdoch, B. E. (2010). The cerebellum and language: Historical perspective and review. Cortex, 46(7), 858–868.

Nakatani, H., Nakamura, Y., & Okanoya, K. (2022). Respective involvement of the right cerebellar crus I and II in syntactic and semantic processing for comprehension of language. The Cerebellum, 22(4), 739–755. 10.1007/s12311-022-01451-y

Nummela, S., Thewissen, J. G. M., Bajpai, S., Hussain, S. T., & Kumar, K. (2004). Eocene evolution of whale hearing. Nature, 430(7001), 776–778.

Oelschläger, H. H. A., Haas-Rioth, M., Fung, C., Ridgway, S. H., & Knauth, M. (2007). Morphology and evolutionary biology of the dolphin (*Delphinus* sp.) brain – MR imaging and conventional histology. Brain, Behavior and Evolution, 71(1), 68–86. 10.1159/000110495

Oelschläger, H. H. A. (2008). The dolphin brain—A challenge for synthetic neurobiology. Brain Research Bulletin, 75(2-4), 450–459. 10.1016/j.brainresbull.2007.10.051

Ohyama, T., Nores, W. L., Murphy, M., & Mauk, M. D. (2003). What the cerebellum computes. Trends in Neurosciences, 26(4), 222–227.

Orekhova, K., Selmanovic, E., De Gasperi, R., Gama, M. A., Wicinski, B., Maloney, B., Seifert, A. C., Alipour, A., Balchandani, P., Gerussi, T., Graïc, J. M., Centelleghe, C., Di Guardo, G., Mazzariol, S., & Hof, P. R. (2022). Multimodal assessment of bottlenose dolphin auditory nuclei using 7-Tesla MRI, immunohistochemistry and stereology. Veterinary Sciences, 9(12), 692–692. 10.3390/vetsci9120692

Park, T., Fitzgerald, E. M. G., & Evans, A. R. (2016). Ultrasonic hearing and echolocation in the earliest toothed whales. Biology Letters, 12(4), 20160060. 10.1098/rsbl.2016.0060

Paulin, M. G. (1993). The role of the cerebellum in motor control and perception. *Brain*, Behavior and Evolution, 41(1), 39–50.

Pettit, S. G., & McCulloch, S. P. (2023). Odontocetes (“toothed whales”): Cognitive science and moral standing – are dolphins persons? Journal of Applied Animal Ethics Research, 5(1), 1–36. 10.1163/25889567-bja10033

Pidoux, L., Le Blanc, P., Levenes, C., & Leblois, A. (2018). A subcortical circuit linking the cerebellum to the basal ganglia engaged in vocal learning. Elife, 7, e32167.

Plogmann D., Kruska D. (1990). Volumetric comparison of auditory structures in the brains of European wild boars (*Sus scrofa*) and domestic pigs *(Sus scrofa* f. dom.). Brain, Behavior and Evolution, 35, 146–155. doi:10.1159/000115863

Popov, V.V., Ladygina, T.F. & Supin, A.Y. (1986). Evoked potentials of the auditory cortex of the porpoise, *Phocoena phocoena*. Journal of Comparative Physiology. A 158, 705–711. doi:10.1007/BF00603828.

Powell, E. W., & Hatton, J. B. (1969). Projections of the inferior colliculus in cat. Journal of Comparative Neurology, 136(2), 183–192.

Raghanti, M. A., Wicinski, B., Meierovich, R., Warda, T., Dickstein, D. L., Reidenberg, J. S., Tang, C. Y., George, J. C., Hans Thewissen, J. G. M., Butti, C., & Hof, P. R. (2018). A comparison of the cortical structure of the bowhead whale (*Balaena mysticetus*), a basal mysticete, with other cetaceans. The Anatomical Record, 302(5), 745–760. 10.1002/ar.23991

Reddi, S. J. (2017). Understanding the relationship between functional and structural connectivity of brain networks. The Neuroscientist, 21.

Revishchin, A. V., & Garey, L. J. (1990). The thalamic projection to the sensory neocortex of the porpoise, *Phocoena phocoena*. Journal of Anatomy, 169, 85–102. https://pubmed.ncbi.nlm.nih.gov/2384340/

Ridgway, S. H., Bullock, T. H., Carder, D. A., Seeley, R. L., Woods, D., & Galambos, R. (1981). Auditory brainstem response in dolphins. Proceedings of the National Academy of Sciences, 78(3), 1943–1947.

Ridgway, S. H. (1990). The central nervous system of the bottlenose dolphin. In S. Leatherwood & R. R. Reeves (Eds.), The Bottlenose Dolphin (pp. 69–97). Academic Press.

Ridgway, S. H., Houser, D., Finneran, J., Carder, D., Keogh, M., Van Bonn, W., Smith, C., Scadeng, M., Dubowitz, D., Mattrey, R., & Hoh, C. (2006). Functional imaging of dolphin brain metabolism and blood flow. Journal of Experimental Biology, 209(15), 2902–2910. 10.1242/jeb.02348

Ridgway, S. H., & Hanson, A. C. (2014). Sperm whales and killer whales with the largest brains of all toothed whales show extreme differences in cerebellum. *Brain*, Behavior and Evolution, 83(4), 266–274. 10.1159/000360519

Ridgway, S., Samuelson Dibble, D., Van Alstyne, K., & Price, D. (2015). On doing two things at once: Dolphin brain and nose coordinate sonar clicks, buzzes and emotional squeals with social sounds during fish capture. Journal of Experimental Biology, 218(24), 3987–3995.

Ridgway, S. H., Carlin, K. P., Van Alstyne, K. R., Hanson, A. C., & Tarpley, R. J. (2017). Comparison of dolphins’ body and brain measurements with four other groups of cetaceans reveals great diversity. Brain Behavior and Evolution, 88(3-4), 235–257.

Ruesch, A., Acharya, D., Bulger, E., Cao, J., McKnight, J., Manley, M., Fahlman, A., Shinn-Cunningham, B., & Kainerstorfer, J. M. (2023). Evaluating feasibility of functional near-infrared spectroscopy in dolphins. Journal of Biomedical Optics, 28(07). 10.1117/1.jbo.28.7.075001

Ryabov, V. A. (2023). The role of asymmetry of the left and right external ear of bottlenose dolphin (*Tursiops truncatus*) in the spatial localization of sound. Acoustical Physics, 69(1), 119–131.

Saldaña, E., & Merchań, M. A. (1992). Intrinsic and commissural connections of the rat inferior colliculus. Journal of Comparative Neurology, 319(3), 417–437.

Schilling, K. G., Daducci, A., Maier-Hein, K., Poupon, C., Houde, J. C., Nath, V., … & Descoteaux, M. (2019). Challenges in diffusion MRI tractography–Lessons learned from international benchmark competitions. Magnetic Resonance Imaging, 57, 194–209.

Schmahmann, J. D., Doyon, J., McDonald, D., Holmes, C., Lavoie, K., Hurwitz, A. S., Kabani, N., Toga, A., Evans, A., & Petrides, M. (1999). Three-dimensional MRI atlas of the human cerebellum in proportional stereotaxic space. NeuroImage, 10(3), 233–260. 10.1006/nimg.1999.0459

Schmahmann, J.D. (2005). Cerebellar motor and cognitive functions. In H. Leiguardia, H. J. Freund, M. Jeannerod, M. Hallett (Eds.), Higher order motor disorders: From neuroanatomy and neurobiology to clinical neurology (pp. 105–121). Oxford University Press.

Schulmeyer, F. J., Adams, J. C., & Oelschläger, H. H. (2000). Specialized sound reception in dolphins—A hint for the function of the dorsal cochlear nucleus in mammals. Historical Biology, 14(1-2), 53–56.

Shepherd, T. M., Flint, J. J., Thelwall, P. E., Stanisz, G. J., Mareci, T. H., Yachnis, A. T., & Blackband, S. J. (2009). Postmortem interval alters the water relaxation and diffusion properties of rat nervous tissue--Implications for MRI studies of human autopsy samples. NeuroImage, 44(3), 820–826. 10.1016/j.neuroimage.2008.09.054

Sitek, K., Calabrese, E., Johnson, G. A., Ghosh, S., & Chandrasekaran, B. (2022). Structural connectivity of human inferior colliculus subdivisions using in vivo and post mortem diffusion MRI tractography. Frontiers in Neuroscience, 16. 10.3389/fnins.2022.751595

Snider, R. S., & Stowell, A. (1944). Receiving areas of the tactile, auditory, and visual systems in the cerebellum. Journal of Neurophysiology, 7, 331–357.

Sokolov V.E., Ladygina T.F. & Supin A.Y. (1972) Localization of sensory zones in the dolphin’s cerebral cortex. Dokl. Akad. Nauk. SSSR 202, 490–493. PubMed.

Suga, N., & Horikawa, J. (1986). Multiple time axes for representation of echo delays in the auditory cortex of the mustached bat. Journal of Neurophysiology, 55(4), 776–805.

Sun, X., Jen, P. H. S., & Kamada, T. (1983). Mapping of the auditory area in the cerebellar vermis and hemispheres of the mustache bat, *Pteronotus parnellii parnellii*. Brain Research, 271(1), 162–165.

Tanaka, H., Ishikawa, T., Lee, J., & Kakei, S. (2020). The cerebro-cerebellum as a locus of forward model: A review. Frontiers in Systems Neuroscience, 14, 19.

Tarpley, R. J., & Ridgway, S. H. (1994). Corpus callosum size in delphinid cetaceans. Brain, Behavior and Evolution, 44(3), 156–165

Toro, R. (2015). *Bottlenose dolphin*. Braincatalogue.org. https://braincatalogue.org/Bottlenose_dolphin

Thaploo, D., Joshi, A., Georgiopoulos, C., Warr, J., & Hummel, T. (2022). Tractography indicates lateralized differences between trigeminal and olfactory pathways. NeuroImage, 261, 119518. 10.1016/j.neuroimage.2022.119518

Thornton, S. W., Mclellan, W. A., Rommel, S. A., Dillaman, R. M., Nowacek, D. P., Koopman, H. N., & Ann Pabst, D. (2015). Morphology of the nasal apparatus in pygmy (*Kogia breviceps*) and dwarf (*K. sima*) sperm whales. The Anatomical Record, 298(7), 1301–1326.

Tyack, P. L., & Sayigh, L. S. (1997). Vocal learning in cetaceans. In C. T. Snowdon & M. Hausberger (Eds.), Social influences on vocal development (pp. 208–233). Cambridge University Press. 10.1017/CBO9780511758843.011

Van Overwalle, F., Ma, Q., & Heleven, E. (2020). The posterior crus II cerebellum is specialized for social mentalizing and emotional self-experiences: A meta-analysis. Social Cognitive and Affective Neuroscience, 15(9), 905–928. 10.1093/scan/nsaa124

Vernes, S. C., Janik, V. M., Fitch, W. T., & Slater, P. J. B. (2021). Vocal learning in animals and humans. Philosophical Transactions of the Royal Society B: Biological Sciences, 376(1836). 10.1098/rstb.2020.0234

Vernooij, M. W., Smits, M., Wielopolski, P. A., Houston, G. C., Krestin, G. P., & van der Lugt, A. (2007). Fiber density asymmetry of the arcuate fasciculus in relation to functional hemispheric language lateralization in both right- and left-handed healthy subjects: A combined fMRI and DTI study. NeuroImage, 35(3), 1064–1076. 10.1016/j.neuroimage.2006.12.041

Vogler, D. P., Robertson, D., & Mulders, W. H. A. M. (2016). Influence of the paraflocculus on normal and abnormal spontaneous firing rates in the inferior colliculus. Hearing Research, 333, 1–7. 10.1016/j.heares.2015.12.021

Wang, Y., Xue, C., Liu, B., Liu, W., & Shiffrin, R. M. (2020). Understanding the relationship between human brain structure and function by predicting the structural connectivity from functional connectivity. IEEE Access, 8, 209926–209938. 10.1109/access.2020.3039837

Whitehead, L., & Banihani, S. (2013). The evolution of contralateral control of the body by the brain: Is it a protective mechanism? *Laterality: Asymmetries of Body*, Brain and Cognition, 19(3), 325–339. 10.1080/1357650x.2013.824461

Wright, A. K., Theilmann, R. J., Ridgway, S. H., & Scadeng, M. (2018). Diffusion tractography reveals pervasive asymmetry of cerebral white matter tracts in the bottlenose dolphin (*Tursiops truncatus*). Brain Structure & Function, 223(4), 1697–1711. 10.1007/s00429-017-1525-9

Zook J. M., Jacobs M. S., Glezer I., Morgane P. J. (1988). Some comparative aspects of auditory brainstem cytoarchitecture in echolocating mammals: Speculations on the morphological basis of time-domain signal processing. In P. E. Nachtigall & P. W. B. Moore (Eds.), Animal Sonar: Processes and Performance (pp. 311–316). NATO ASI Science; Springer.

