## Supplementary figures and images for "Lateralized cerebellar connectivity differentiates auditory pathways in echolocating and non-echolocating whales"

### 3D rotation of atlantic white sided dolphin ascending auditory tracts

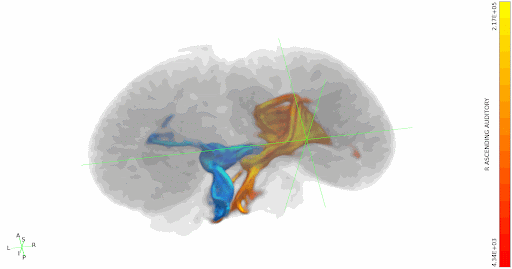

### 3D rotation of atlantic white sided dolphin IC-contralateral cortex tracts

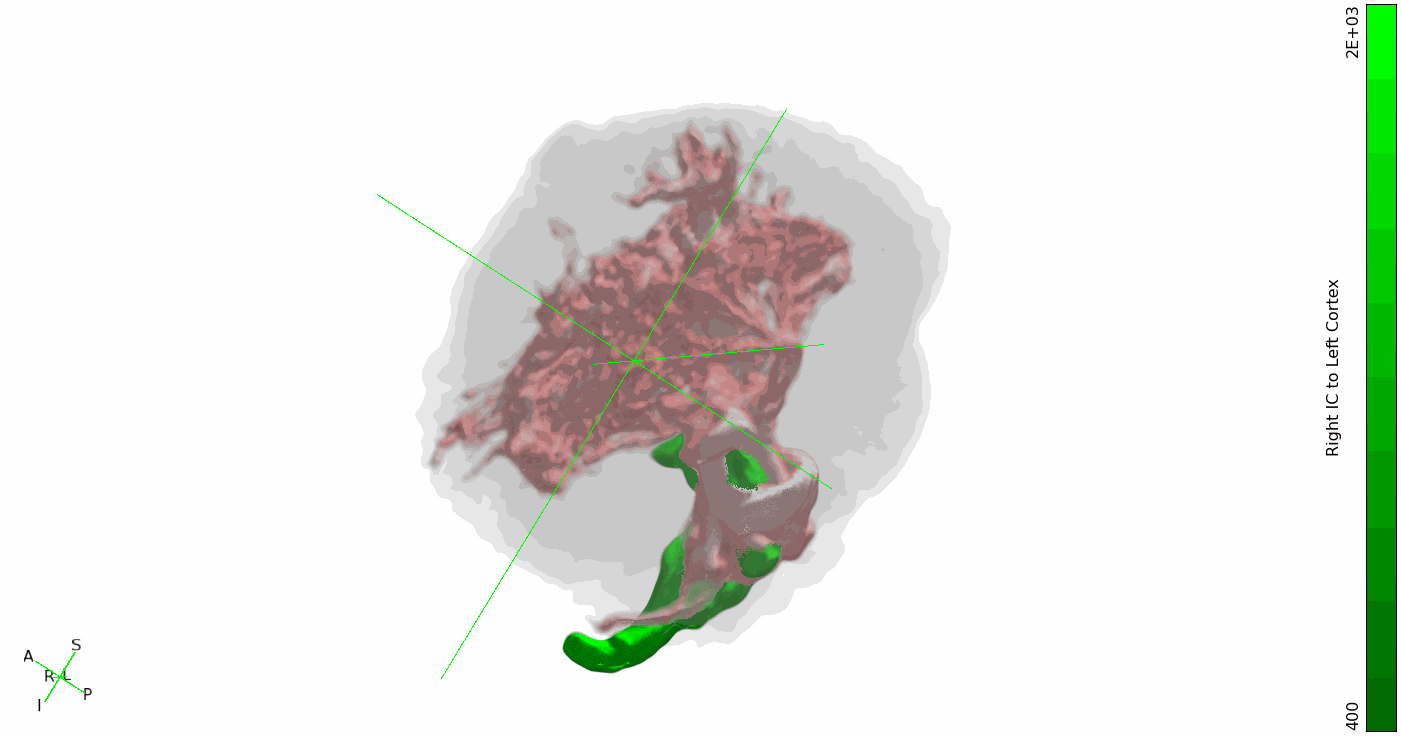

### 3D rotation of atlantic whited dolphin IC-cerebellum tracts

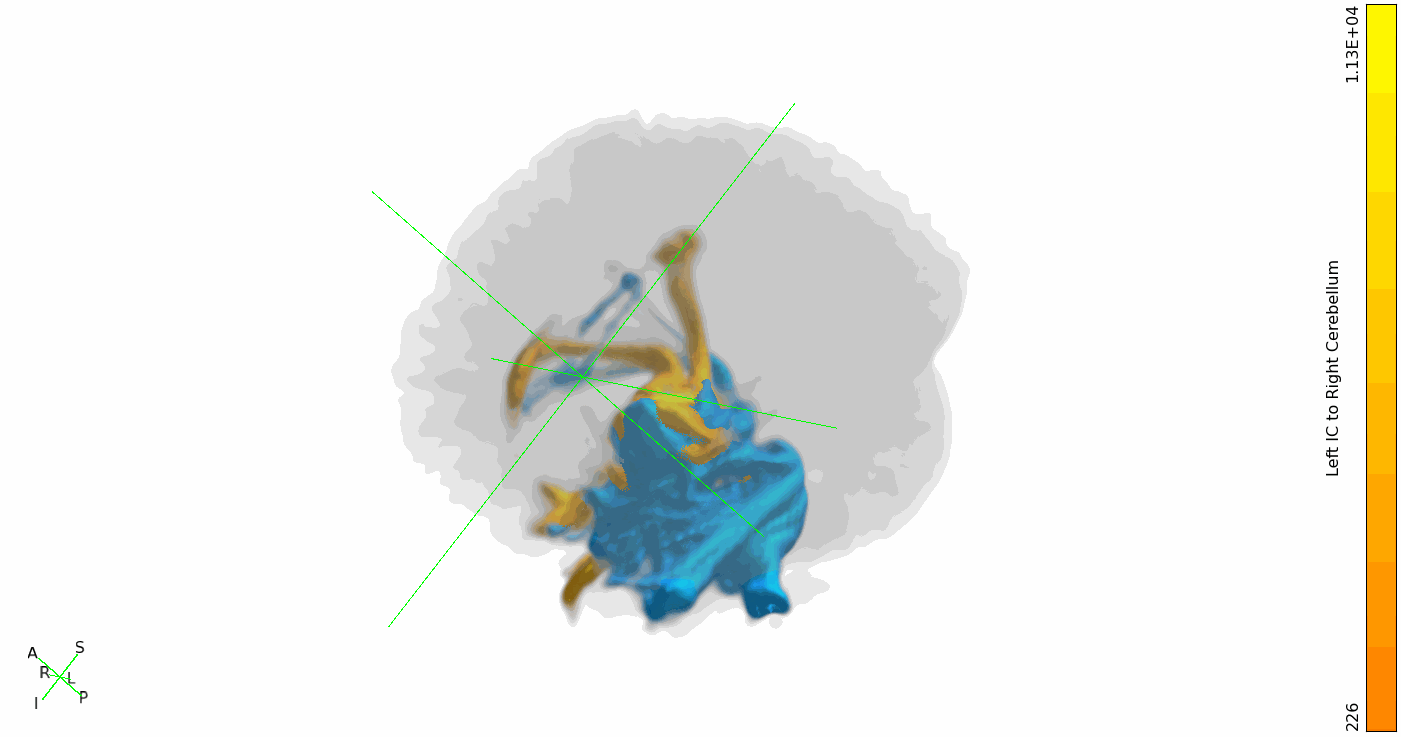

### 3D rotation of common dolphin ascending auditory tracts

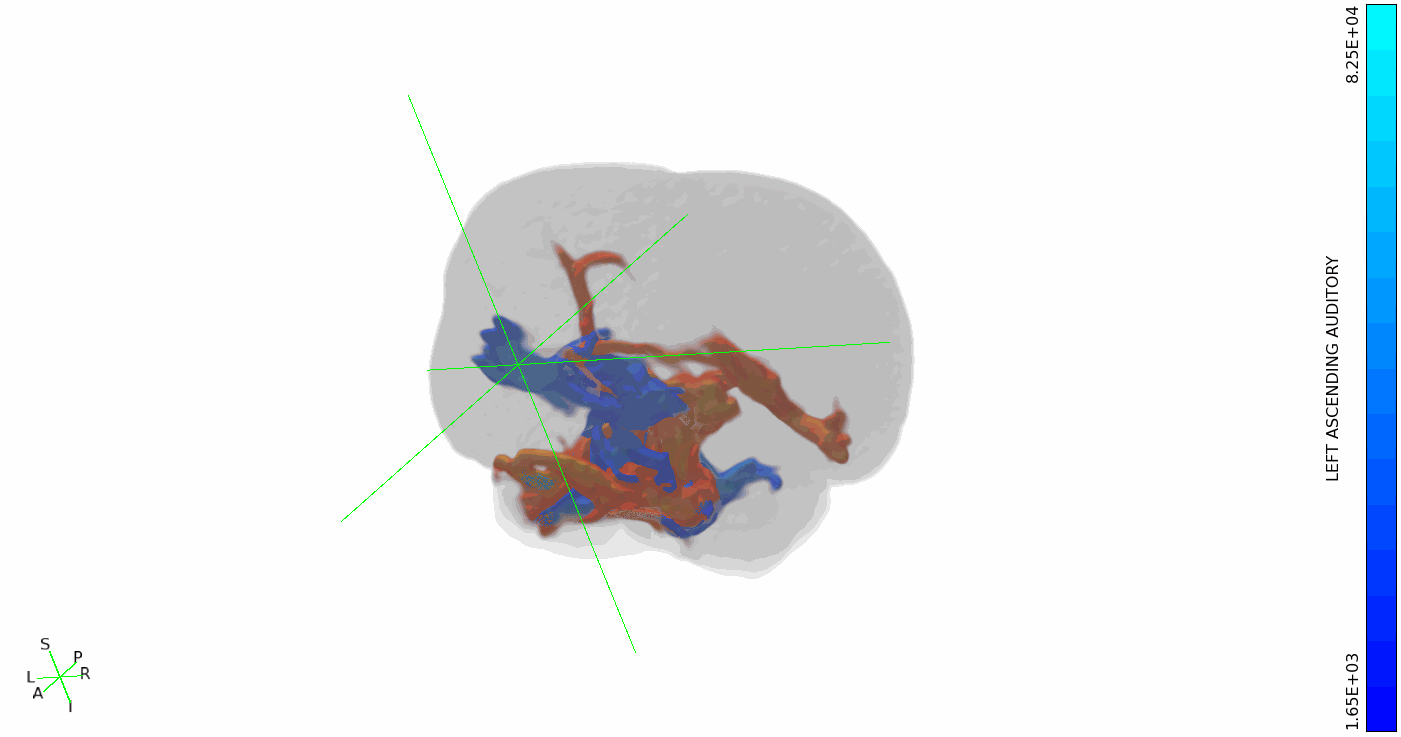

### 3D rotation of common dolphin IC-cerebellum tracts

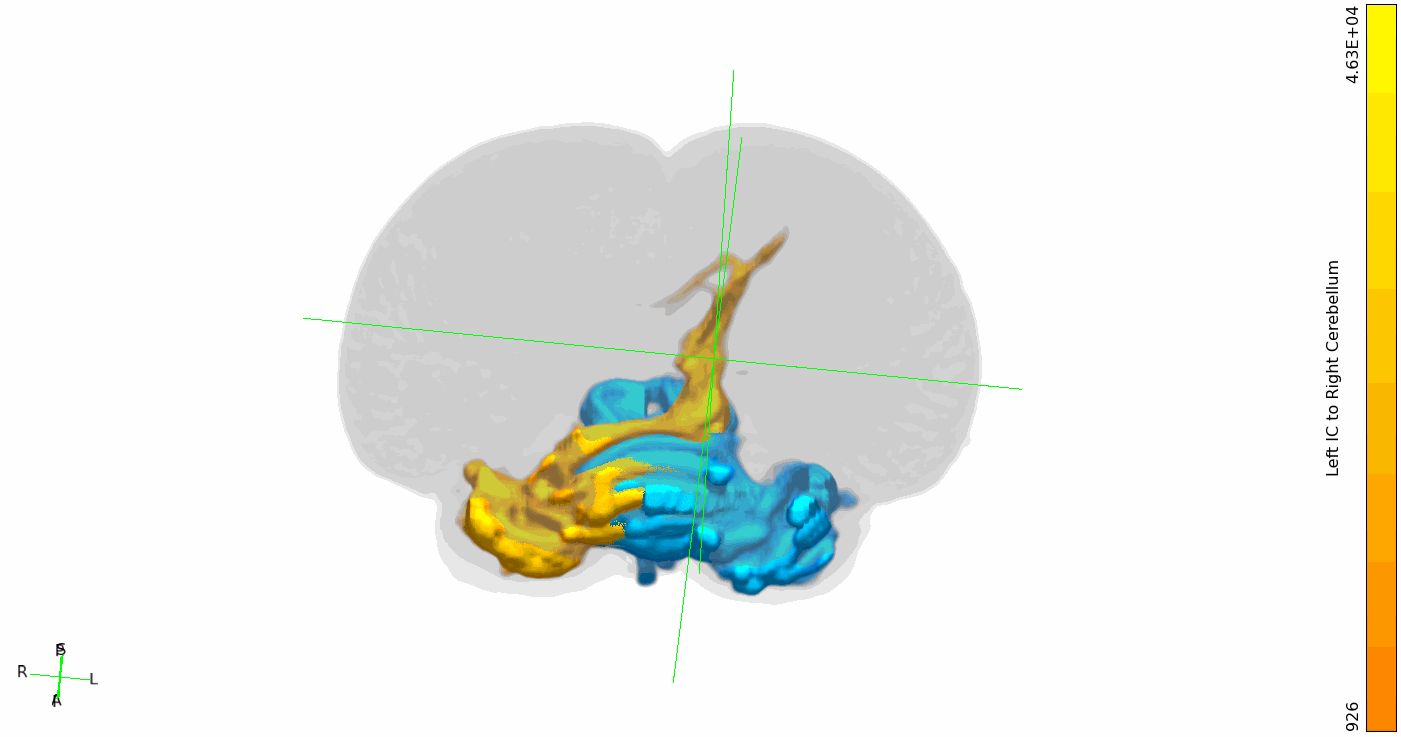

### 3D rotation of common dolphin IC-contralateral cortex tracts

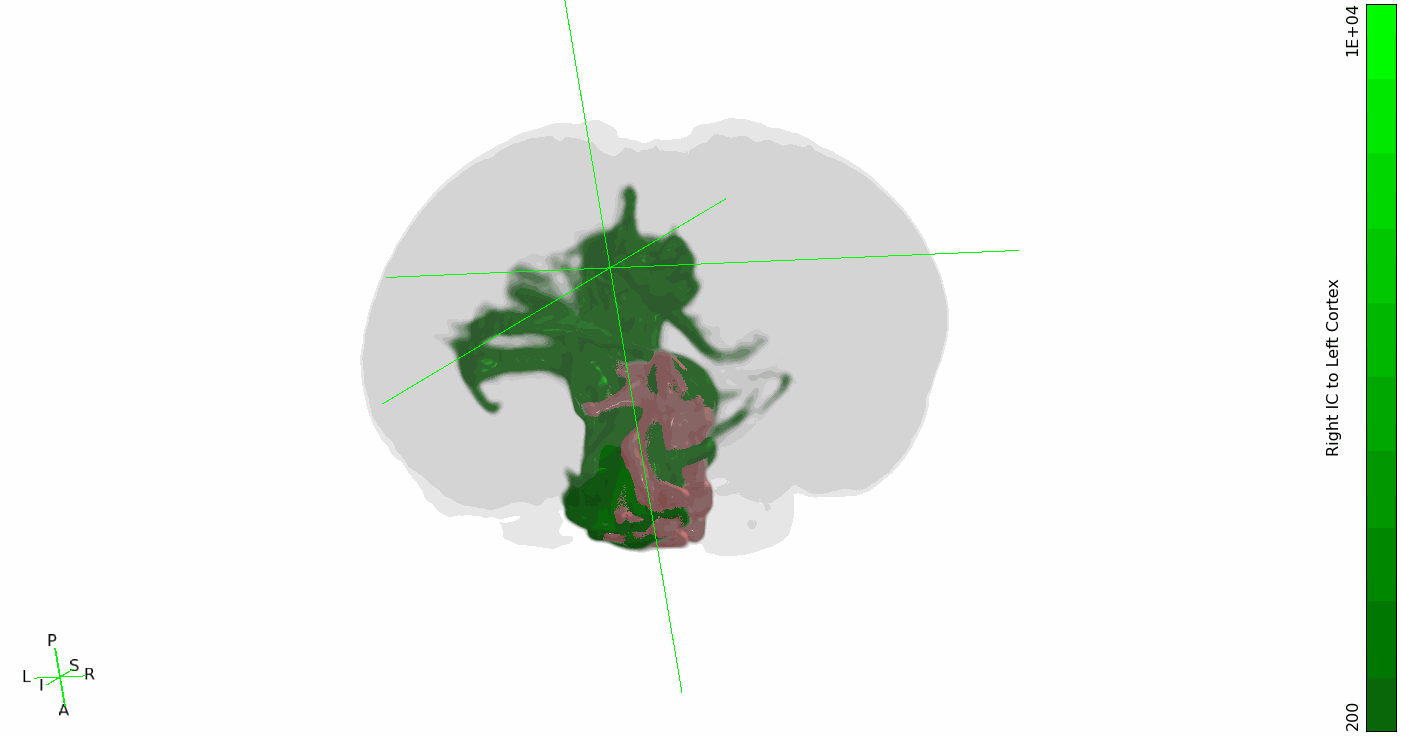

### 3D rotation of pantropical spotted dolphin ascending auditory tracts

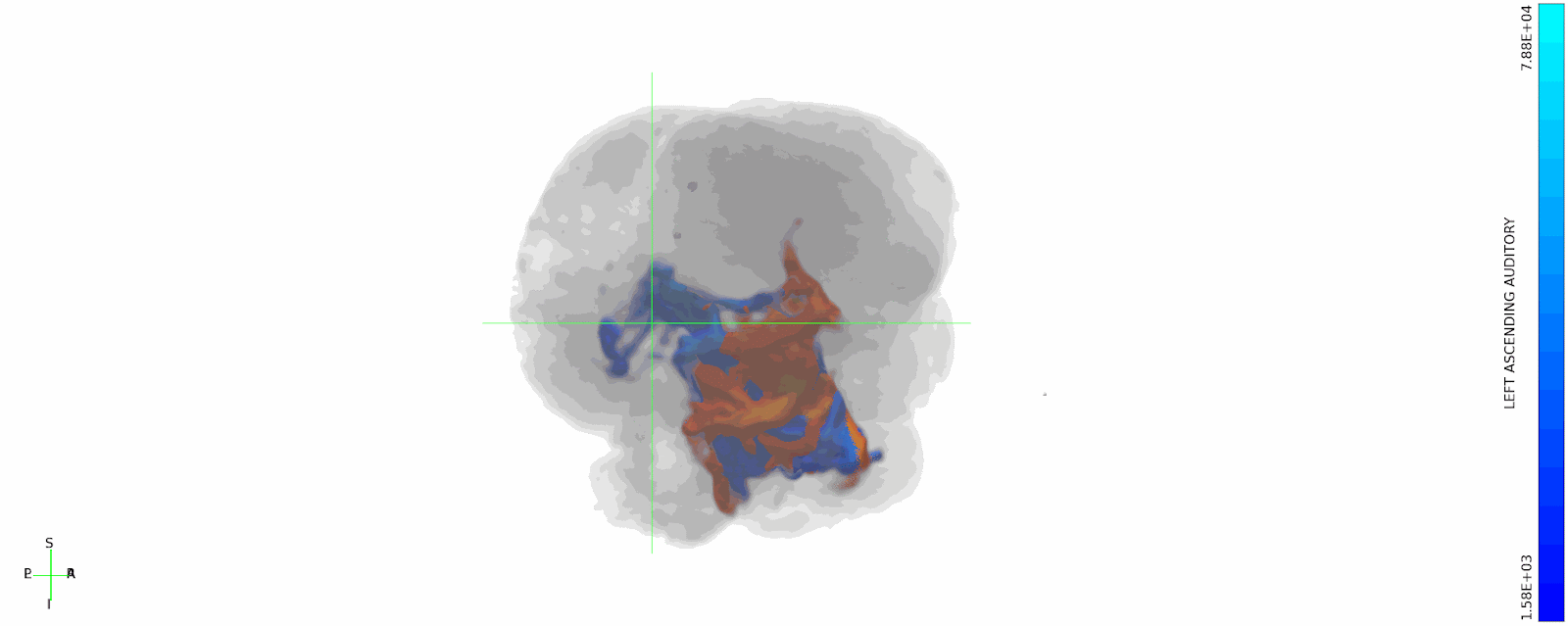

### 3D rotation of pantropical spotted dolphin IC-cerebellum tracts

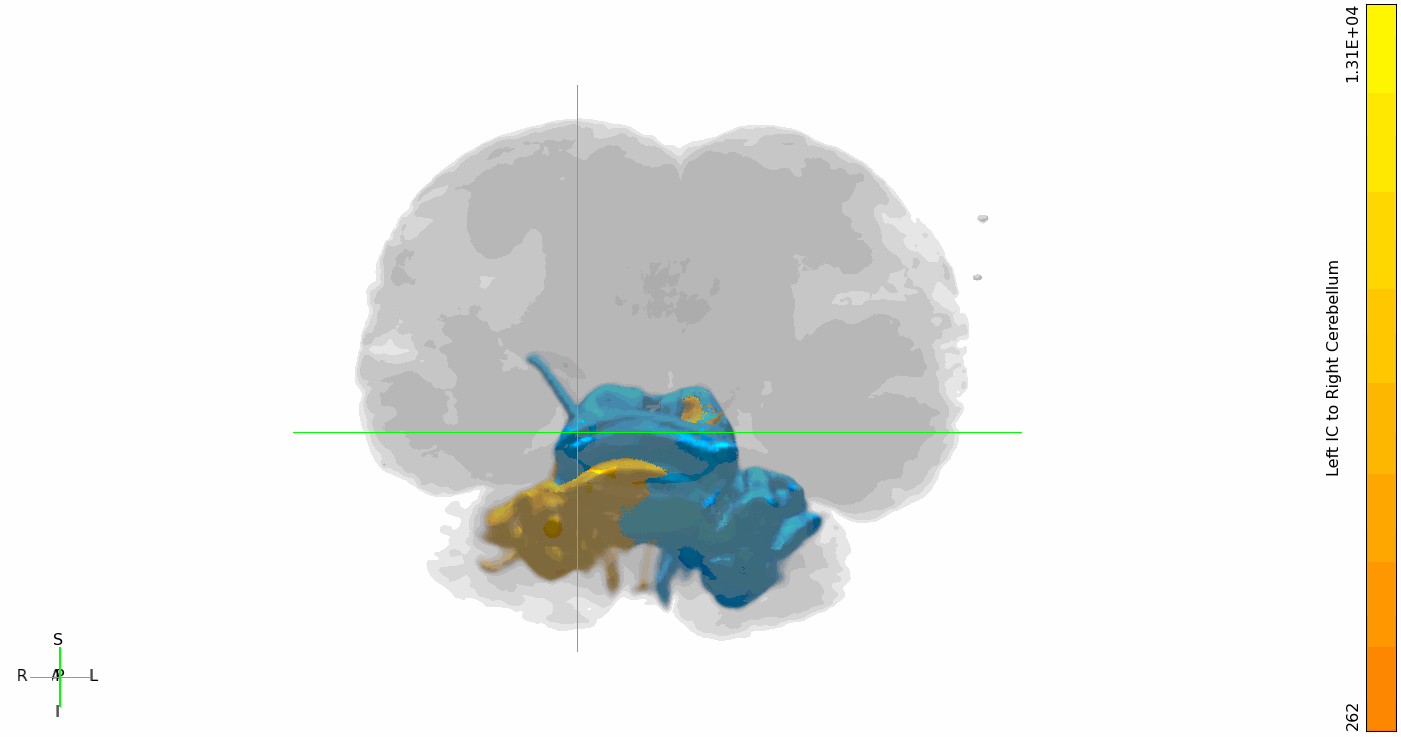

### 3D rotation of pantropical spotted dolphin IC-contralateral cortex tracts

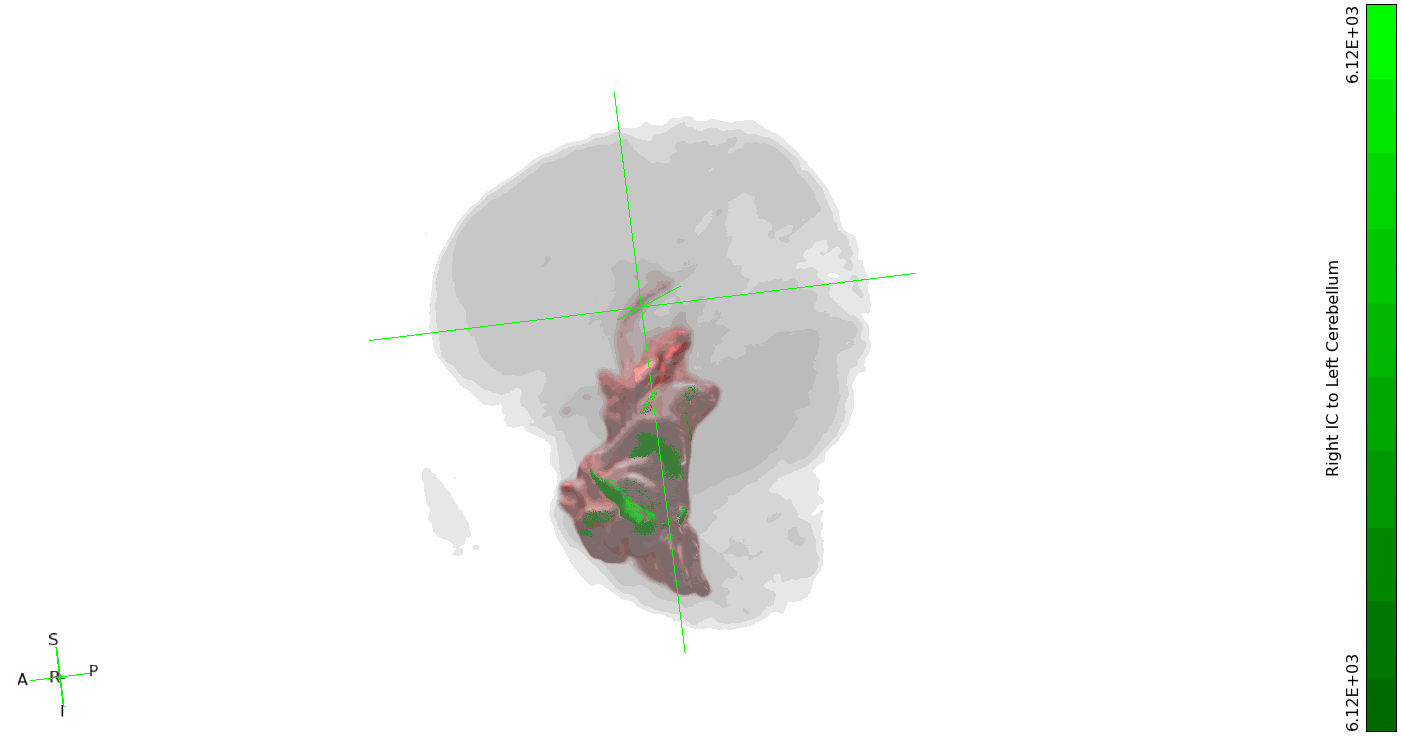

### 3D rotation of sei whale ascending auditory tracts

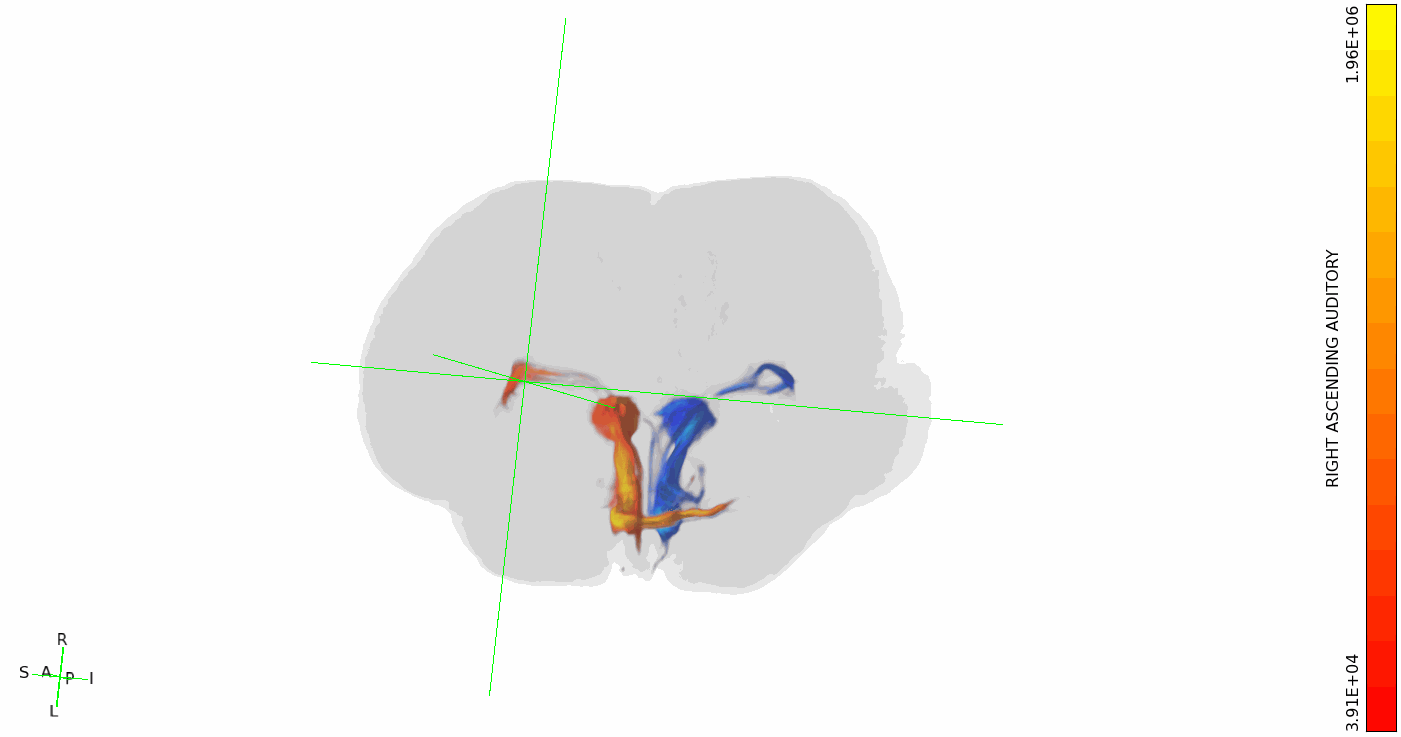

### 3D rotation of sei whale IC-cerebellum tracts

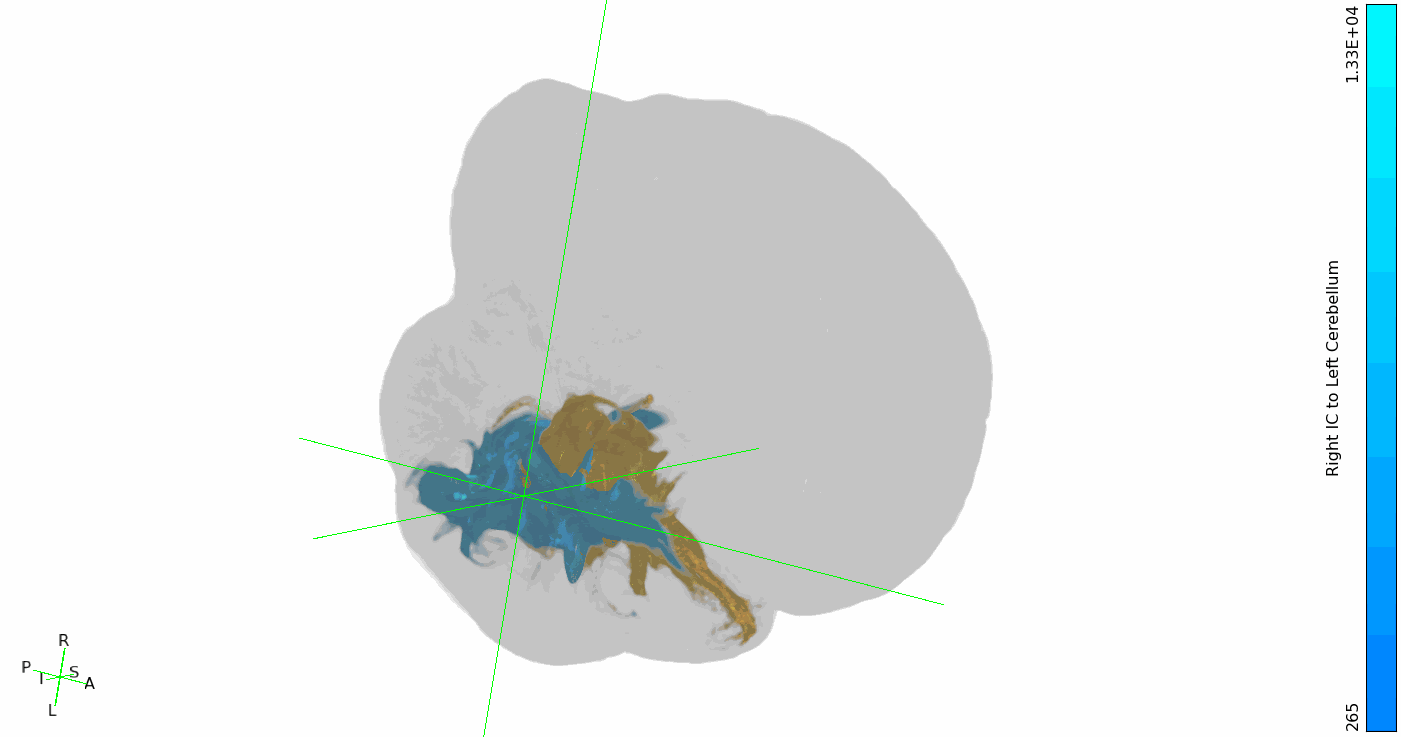

### 3D rotation of sei whale IC-contralateral cortex tracts

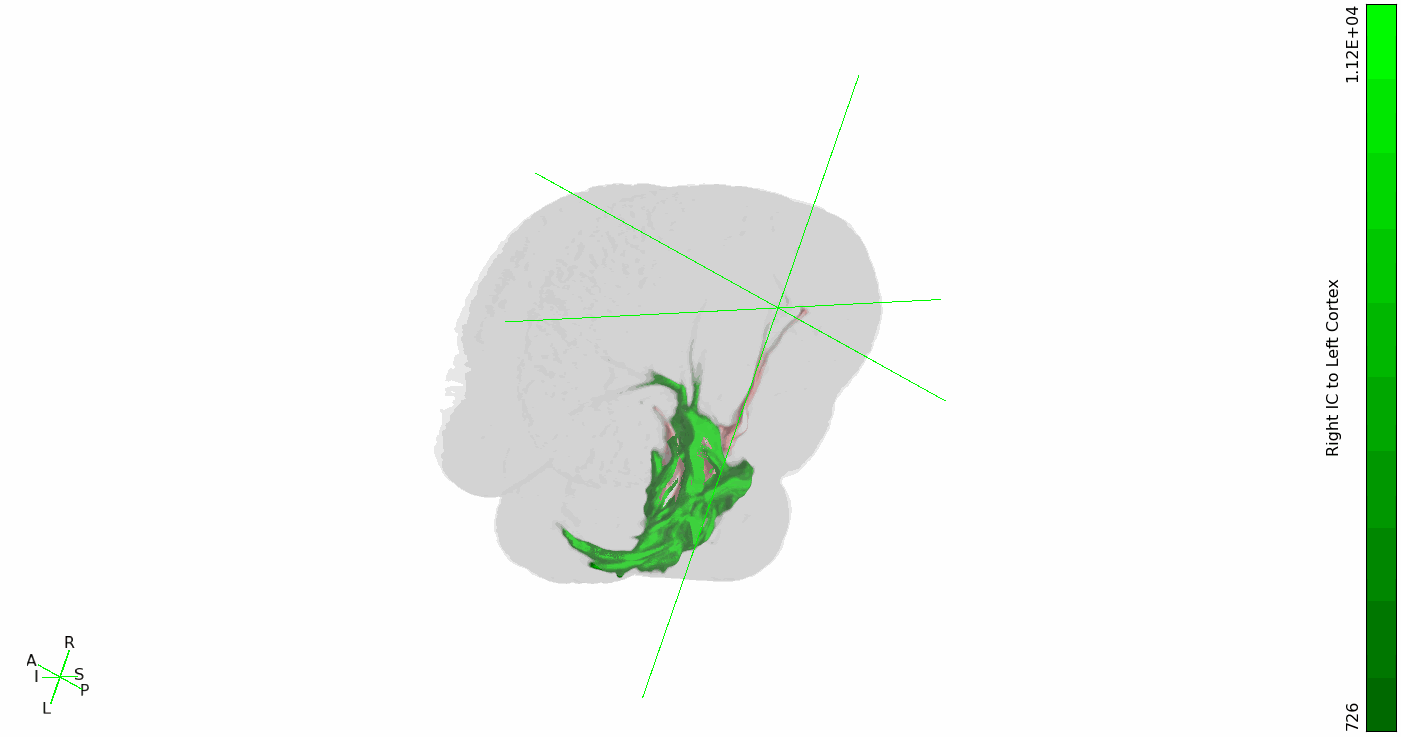
